# Parallel molecular computation on digital data stored in DNA

**DOI:** 10.1101/2022.08.17.504328

**Authors:** Boya Wang, Siyuan S. Wang, Cameron Chalk, Andrew D. Ellington, David Soloveichik

## Abstract

DNA is an incredibly dense storage medium for digital data, but computing on the stored information is expensive and slow (rounds of sequencing, *in silico* computation, and DNA synthesis). Augmenting DNA storage with “in-memory” molecular computation, we use strand displacement reactions to algorithmically modify data stored in the topological modification of DNA. A secondary sequence-level encoding allows high-throughput sequencing-based readout. We show multiple rounds of binary counting and cellular automaton Rule 110 computation on 4-bit data registers, as well as selective access and erasure. Avoiding stringent sequence design, we demonstrate large strand displacement cascades (122 distinct steps) on naturally-occurring DNA sequences. Our work merges DNA storage and DNA computing, setting the foundation of entirely molecular algorithms for parallel manipulation of digital information kept in DNA.

---

DNA is an incredibly dense (up to 455 exabytes per gram, 6 orders of magnitude denser than magnetic or optical media) and stable (readable over millennia) digital storage medium (*2–4*). Storage and retrieval of up to gigabytes of digital information in the form of text, images, and movies have been successfully demonstrated. DNA’s essential biological role ensures that the technology for manipulating DNA will never succumb to obsolescence. However, performing computation on the stored data typically involves sequencing the DNA, electronically computing the desired transformation, and synthesizing new DNA. This expensive and slow loop limits the applicability of DNA storage to rarely accessed data (cold storage).

In contrast to traditional (passive) DNA storage, schemes for dynamic DNA storage allow access and modification of data without sequencing and/or synthesis. Upon binding to molecular probes, files can be accessed selectively (*5*) and modified through PCR amplification (*6*). Introducing or inhibiting binding of molecular probes with existing data barcodes can rename or delete files (*7*). Information encoded in the hybridization pattern of DNA can be written and erased (*8*) and can even undergo basic logic operations such as AND and OR (*9*) using strand displacement. By encoding information in the topology (nicks) of naturally-occurring DNA (a.k.a. native DNA (*10*)) data can be modified through ligation or cleavage (*11*). With image similarities encoded in the binding affinities of DNA query probes and data, similarity searches on databases can be performed through DNA hybridization (*12, 13*). These advances allow information to be directly accessed and edited within the storage medium, albeit with limited or no computation.

In addition to storing information, DNA has been used as a programmable material for computation, primarily via “strand displacement” reactions. With the understanding of the kinetics and thermodynamics of DNA strand displacement (*14–16*), a variety of rationally designed molecular computing devices have been engineered. These include molecular implementations of logic circuits (*17–19*), neural networks (*20, 21*), consensus algorithms (*22*), dynamical systems including oscillators (*23*), and cargo-sorting robots (*24*). Despite the above achievements, the current DNA storage paradigm has not made use of the computational power of strand displacement systems.

Here we propose a new paradigm called SIMD||DNA (Single Instruction Multiple Data ^1^ computation with DNA) which integrates DNA storage with in-memory computation by strand displacement (Figure 1A). Inspired by methods of storing information in DNA topology (nick positions) (*10, 11*), SIMD||DNA encodes information in a multi-stranded DNA complex with a unique pattern of nicks and exposed single-stranded regions (a *register*). Although storage density is somewhat reduced (approximately a factor of 30, see Discussion), encoding information in topology still achieves orders of magnitude higher density than competing magnetic and optical technologies. To manipulate information, an instruction (a set of strands) is applied in parallel to all registers (Figure 1B). The strand composition of a register changes when the applied instruction strands trigger strand displacement reactions within that register. Non-reacted instruction strands and reaction waste products are washed away via magnetic bead separation to prepare for the next instruction. Each instruction can change every bit on every register, yielding a high level of parallelism—our experiments routinely manipulated 10^11^ registers in parallel, each of which could in principle store distinct data.

**Figure 1:**
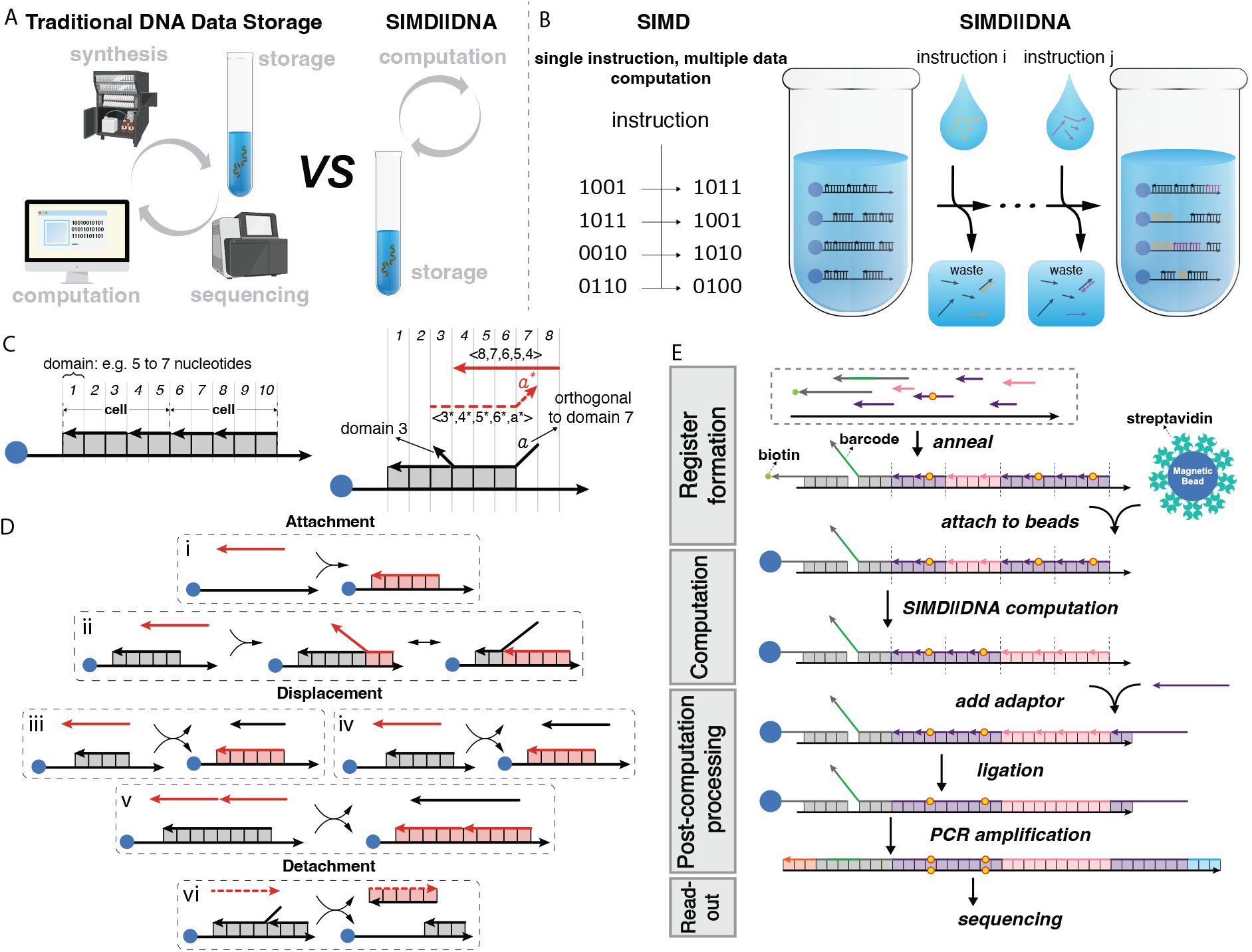
Overview of SIMD||DNA. (A) Instead of outsourcing the computation process to an electronic computer with required sequencing and synthesis steps, SIMD||DNA allows in-memory computation performed by DNA itself. (B) Analogous to SIMD computation in classical computing, in SIMD||DNA a set of instruction strands is added to the solution to simultaneously react with all the registers (multi-stranded complexes with information encoded in the pattern of nicks and exposed single-stranded regions). Magnetic beads (blue) labeling of registers allows separation of unreacted instruction strands and displaced reaction products. (C) Notation. Vertically aligned top and bottom domains are complementary unless specifically labeled (e.g. *a* and *a\**, with “*” indicating complementarity). We assume that domains bind only their complements. Dashed strands share the same sequence as their vertically aligned bottom strands. (D) Instruction strands induce three types of events: attachment, displacement and detachment. The toehold exchange reaction in (iv) is considered irreversible since instruction strands are at high concentration. (E) Experimental workflow. Registers are assembled through annealing and attached to magnetic beads, followed by computation. The post-computation process of ligation and PCR amplification prepares for sequencing readout. Yellow dots indicate top-bottom strand mismatches, allowing data readout after ligation erases topological information (secondary sequence-based encoding).

Additional important features of registers include unique identifiers (“barcodes”) that label the information contained in the register (for example, in putative medical record applications, the barcodes could identify specific patients). Adding a *query* strand with the complement of the barcode displaces the register from the magnetic bead, allowing retrieval or erasure of specific information (e.g., information pertaining to particular patients) without disturbing the rest of the registers. High-throughput readout is achieved via another feature of the registers: a redundant sequence-based encoding that does not play a role in computation but is readable by next generation sequencing (NGS).

Although some DNA strand displacement systems have been adapted to naturally-occurring sequences such as for diagnostic applications (*26*), sophisticated sequence design incompatible with natural sequences is typically used (*27–29*). Naturally-occurring DNA may lead to undesired binding, triggering spurious displacement or preventing displacement from completing. Surprisingly, we show that SIMD||DNA algorithms may be programmed while eschewing careful sequence design and the high costs and low yield of long oligonucleotide phosphoramidite synthesis. M13, a mainstay of DNA origami (*30*), provides a large storage space for parallel SIMD computation on multiple registers. To date we constructed the largest strand displacement system using naturally-occurring DNA sequences: using SIMD||DNA, we implemented in total 122 distinct strand displacement reactions.

## SIMD||DNA

In SIMD||DNA, molecular algorithms manipulate digital information stored in DNA strand topology through DNA strand displacement reactions (Figure 1). Each data register contains a long *bottom* strand and multiple bound short *top* strands. *Domains* represent consecutive nucleotides that act as a functional unit. Strand displacement is initiated by hybridization at a single-stranded *toehold* domain followed by displacement of the incumbent strand. The domain lengths are chosen so that ideally: (1) each domain can initiate strand displacement (i.e. can act as a toehold), (2) strands bound by a single domain readily dissociate, and (3) strands bound by two or more domains cannot dissociate. The bottom strand is partitioned into sets of consecutive domains called *cells*, each representing a bit of information (Figure 1C).

Each *instruction* of a program corresponds to the addition of a set of DNA strands at high concentration to registers bound to magnetic beads. Instruction strands are allowed to react for a short time before washing to prevent waste products from interacting with registers.

The instruction strands cause three types of events (Figure 1D): **Attachment** events attach new top strands to the register, preserving all the previously-bound strands. Attachment can include partial displacement of a pre-existing top strand on the register (Figure 1D(ii)). In **displacement** events, an instruction strand displaces and replaces a previously-bound top strand. This event can occur only if a toehold is available for the displacement. Note that since the displaced strands are in relatively low concentration, we assume they do not subsequently react with the registers. (We assume toehold exchange (*14*) is irreversible in our model as shown in Figure 1D(iv).) Two instruction strands can also cooperatively (*31*) displace strands on the register (Figure 1D(v)). In a **detachment** event, an instruction strand displaces a complementary top strand from a register without introducing a new top strand. This event can only occur if there is a toehold on the top strand to initiate the displacement.

A secondary sequence-level encoding enables parallel reading out of the data stored in a heterogeneous pool of registers. It consists of single-base mismatches on a subset of the top strands compared to the bottom strand, allowing us to distinguish different top strands even after after ligation and PCR amplification. To ensure fast kinetics of displacement events, any mismatch is on the displaced strand (*32*); see Section S3.4. We subsequently use NGS to read proportionally amplified computation products.

## Binary Counting Program

The binary counting SIMD||DNA program treats each register as a binary integer, and increments all the registers by one. Abstractly, a binary increment operation flips the right-most (least significant) zero and all bits to its right. At the high-level, our SIMD algorithm does the following (Figure 2A): Starting from the rightmost domain (designed to initiate displacement), the program erases all 1’s in between the rightmost cell and the rightmost value-0 cell (Instructions 1 and 2), and changes those cells to 0 at Instructions 4 and 5. The rightmost value-0 cell is first marked (Instruction 3), and then changed to value 1 (Instructions 6 and 7). (See Section S1 for the proofs of correctness of our programs.)

**Figure 2:**
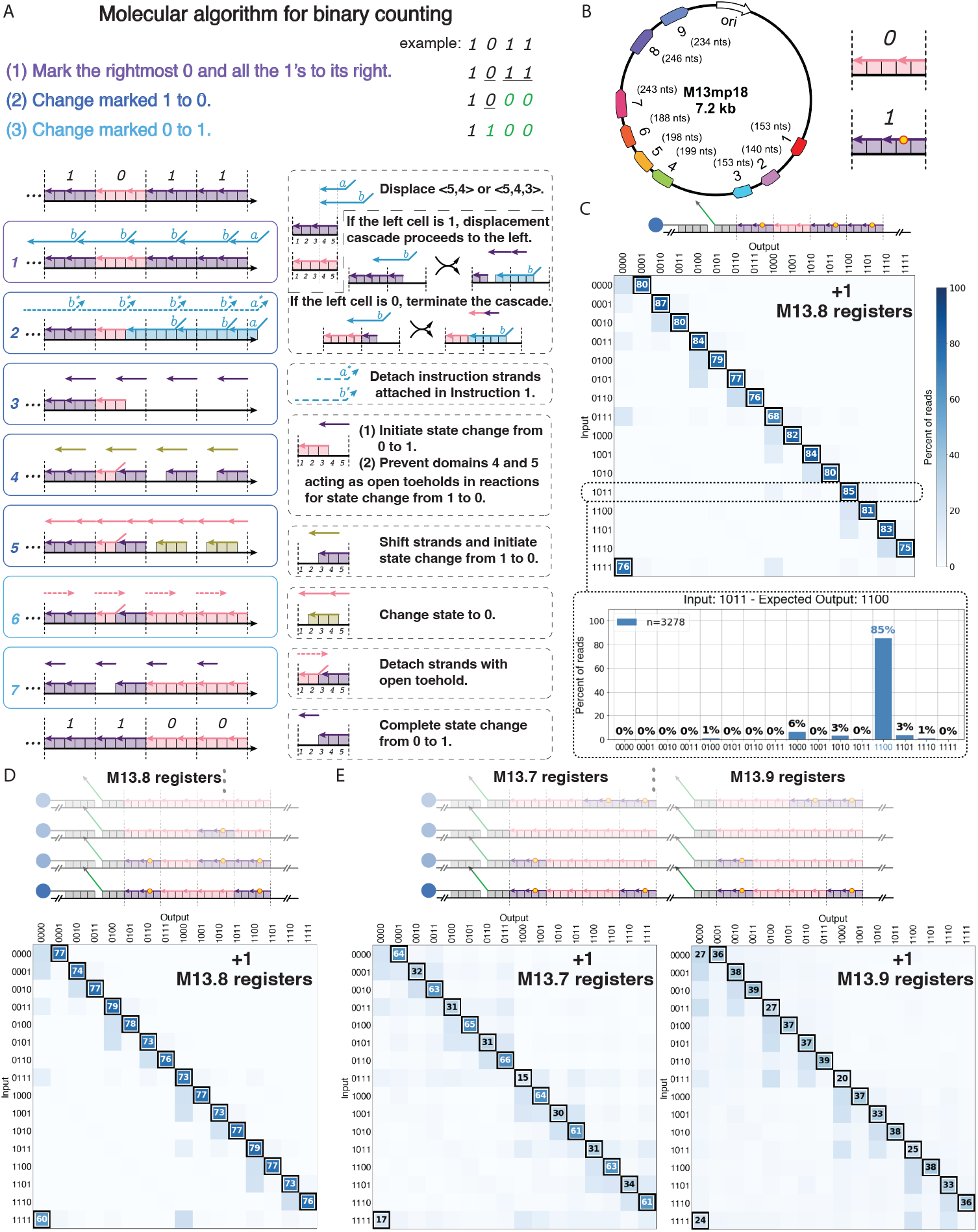
Binary counting program on naturally-occurring (M13) sequences. (A) Molecular program implementing binary counting. The example register (top) encoding the initial value updates to the new value (bottom) after 7 instructions. Strands are labeled by colors: value 1 (purple), value 0 (pink), intermediates (others). Solid boxes show the instruction strands and the configuration of the register before reacting. Dashed boxes explain the logical meaning of the instructions. (B) Locations of registers on M13mp18 and the encoding of 0 and 1. In the text, M13.i refers to registers at location choice *i*. (C) NGS results of SISD binary counting (40 °C) on M13.8 registers. For each initial value, the distribution of the output values are represented in the heat-map matrix. Lower bar plot shows an example of the distribution of output values in one row (input 1011) of the heat map. (D) NGS results of SIMD binary counting (40 °C) on M13.8 registers. (E) NGS results of SIMD binary counting (50 °C) on M13.7 and M13.9 registers in parallel. In the heat maps, black borders indicate the correct output value; the percent value is shown if the output value is at least 25% of all reads for a given sample.

We show that strand displacement computation can proceed directly on naturally-occurring M13mp18 plasmid (M13) sequence space with minimal sequence optimization (Figure 2). To choose the parts of the M13 plasmid to use, we first eliminated areas with 4 consecutive C’s or G’s and hairpin “[A]” (*33*), and then selected 9 random locations as candidates at which we encoded registers (Figure 2B). As a scalable technique to alleviate spurious interactions in naturally-occurring DNA while keeping desired binding intact, we increased reaction temperature and the strength of domain binding (increasing their length). Partitioning the sequence from the chosen locations into domains of different lengths according to the binding energy, M13.1, M13.2, M13.3 registers were designed with weak (short) domains, M13.4, M13.5, M13.6 registers with medium domains, and M13.7, M13.8, M13.9 registers with strong (long) domains (details in Section S3.2). We tested initial values 0010 and 0011 with different reaction temperatures (Section S6) on these 9 registers, and then picked 5 of them for further experiments.

We first performed SISD (single-instruction, single-data) computation on M13.8 registers for each of the 16 4-bit initial values, where each test tube contained only one initial value. After NGS sequencing, the reads were organized according to the barcode sequences associated with their encoded initial values, and the percentage of reads representing the correct value was calculated (Methods in Section S5.2). More than 99% of M13.8 registers that were assembled, processed, and sequenced contained the expected initial value (Figure S4A). After a round of binary counting computation, registers corresponding to all 16 initial values show the correct output as the dominant output (Figure 2C), with the minimum correct percent at 68%. We observed similar results for M13.3 registers (Figure S4B).

To demonstrate SIMD computation, we executed the binary counting program on a pool of M13.8 registers with all 16 initial values in the same test tube (Figure S5A), obtaining a similar correct fraction of computation as for SISD (see Figure 2D). We also achieved similar SIMD computation results on M13.7 and M13.9 registers (Figure S5B, C) at a higher temperature.

The 7.2kb long M13mp18 plasmid is, in principle, capable of accommodating SIMD||DNA computation on several hundreds of bits. As a proof of scaling, we investigated the ability to store (Figure S6) and compute (Figure 2E) simultaneously on M13.7 and M13.9 registers assembled on the same M13 molecules. As shown in Figure 2E, most registers produced the highest readcount for the correct output, albeit with more errors for some inputs. Note that different programs could be simultaneously executed on different registers or registers of orthogonal sequence, thereby potentially generalizing SIMD||DNA to MIMD (multiple instruction, multiple data).

## Cellular Automaton Rule 110 Program

Expanding the diversity of functions computable by SIMD||DNA, we implemented a Turing universal program based on cellular automaton Rule 110 (the full program is shown in Section S1.2). The Turing universality of Rule 110 (*34*) argues that, in principle SIMD||DNA is capable of performing any computation that can be performed on any computer.

An abstract elementary cellular automaton (*35*), one of the simplest models of computation, consists of an infinite set of cells with two states, 0 or 1. At each time step, updates to a cell depend on the states of its left and right neighbors. A simple two-rule characterization of Rule 110 is as follows: 0 updates to 1 if and only if the state to its right is a 1, and 1 updates to 0 if and only if both neighbors are 1. The SIMD||DNA program for implementing a time-step is shown in Figure S1.

To implement the Rule 110 program, we used artificially designed sequences (sequence design explained in Section S3.1) as well as M13. The encoding of bit value 1 for the Rule 110 program contains an exposed (toehold) region; thus to enable ligation and sequencing, “seal” strands were added at the completion of computation to fill in the gaps on the patterns of the top strands (Figure 3A). We performed SIMD computation for all 16 initial values taking the bits to the left and right of the four register bits as 0 (the implementation of this boundary condition is explained in Section S10). The sequencing readout shows that the correct values are the dominant output: Figure 3B shows the data for registers composed of artificially designed sequences (the control without computation is shown in Figure S8A). We achieved similar results using the M13 sequence as seen in Figure S8B.

**Figure 3:**
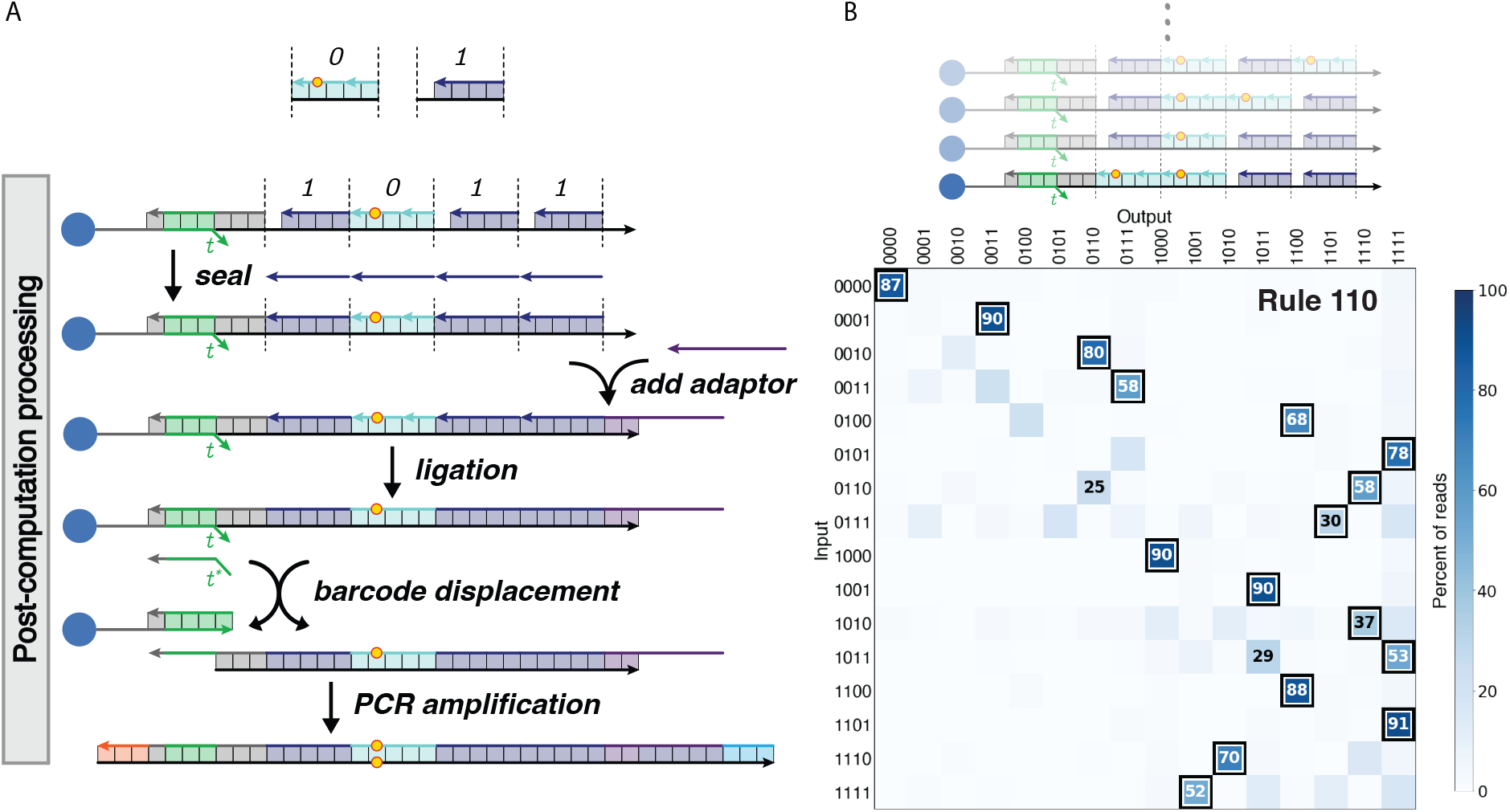
SIMD cellular automaton Rule 110 computation. (A) Post-computation process for the Rule 110 program with chemically synthesized DNA. After computation, “seal” strands are added to fill in the gap for cells representing bit 1 for the following ligation step. Strands complementary to the barcode sequences are added to selectively displace the register with certain initial values. (B) NGS results of SIMD Rule 110 computation (25 °C) on 16 registers with unique initial values. The leftmost and rightmost cells of the input only have one neighbor; thus they undergo a modified Rule 110 update which proceeds as if the missing neighbor were 0 (see Section S10 for the implementation of this boundary condition). The registers were composed of chemically synthesized DNA. The correct output value is indicated with a black border; values that appear in at least 25% of all reads for a given sample are printed.

## Random Access (Selective Retrieval and Selective Erasure)

Random access is the selective reading, modification, or erasure of data within a database, and importantly allows access to a subset of information without expending resources (such as NGS) to read the entire repository each time. Other DNA storage schemes typically use PCR to selectively amplify data (*5, 6*) or selectively pull out information by tuning the binding affinity between sequences (*12, 36*). However, designing sequences or multiplexed orthogonal PCR probes with high specificity can be challenging. In contrast, strand displacement achieves specificity through kinetically and energetically favorable reactions that displace a pre-existing strand (*37*).

To accommodate random access in SIMD||DNA, registers holding different data are prepared with specific barcode sequences that can serve as a point of access. After computation on all registers together, query strands complementary to the barcode sequences can be added to elute registers with the matching barcodes (Figure 3A). Thus, specific registers can be accessed separately for read out from the register mix.

We experimentally demonstrated SIMD computation combined with random access for both the Rule 110 and the binary counting programs (Figure 4). To test random access, we assigned registers barcodes corresponding to their initial inputs. After Rule 110 computation on a mix of registers with all 16 inputs, we added a query strand with the barcode corresponding to input 0011 and processed the displaced registers (ligation, PCR amplification, sequencing). Next, we queried the remaining registers with the barcode corresponding to initial value 1001 in the same way. Finally, the remaining registers were queried with all 16 different barcodes. The sequencing results confirmed that, for the first and second queries, the desired registers accounted for 90% and 67% of the registers displaced from the mix (Figure S9). In random access combined with the binary counting computation (Figure 4B), all queries showed high specificity for the chosen barcode (at least 77%, Figure S10).

**Figure 4:**
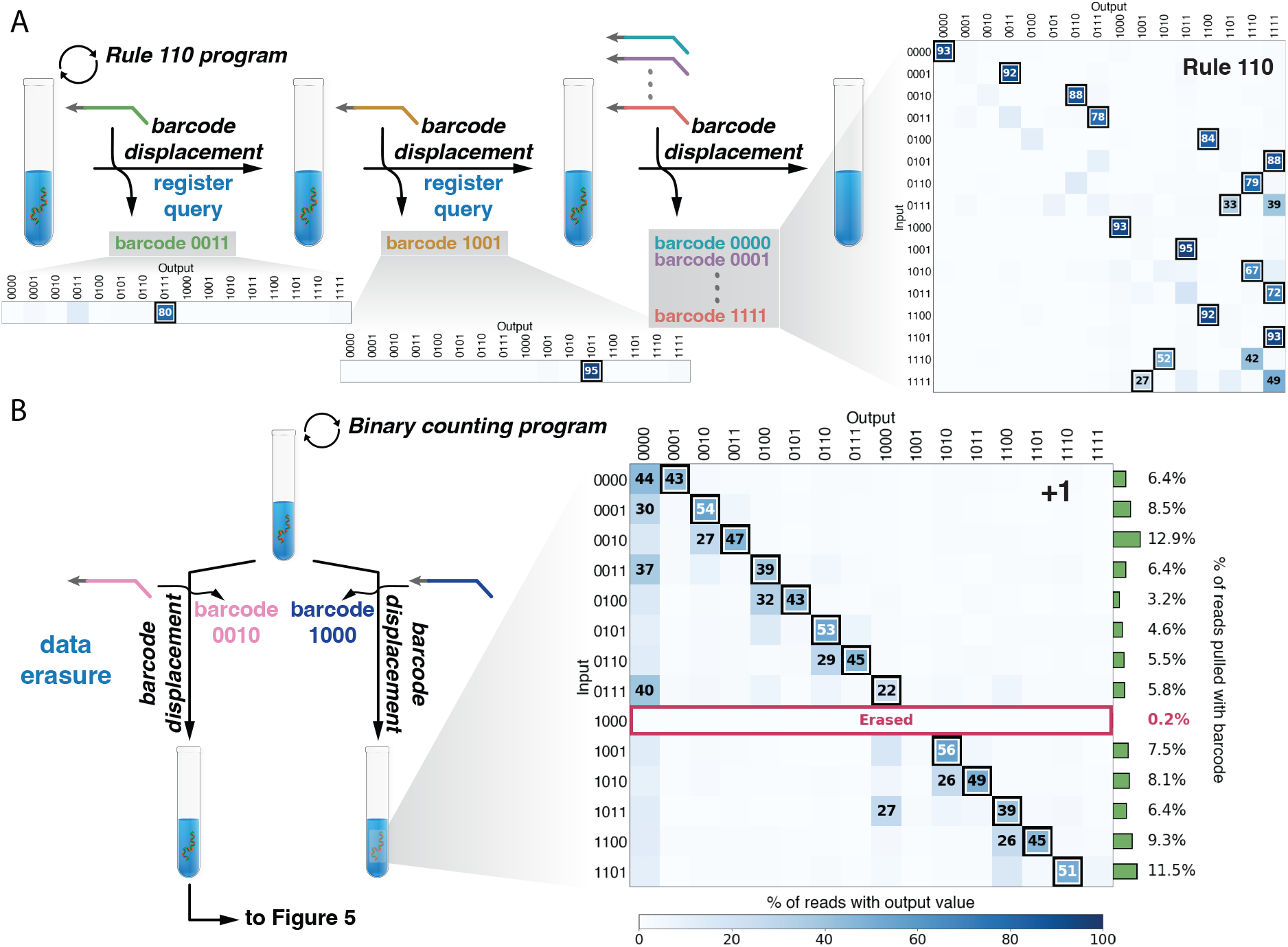
NGS results for random access. (A) Series of selective retrieval operations following Rule 110 computation and (B) Selective erasure after the binary counting computation. (A) Registers with initial value 0011 were accessed first (left), 1001 second (middle), and all remaining values last (right) by adding the query strands corresponding to their barcodes. The query strand concentration was lower than the estimated register concentration; thus displacement achieved partial extraction of registers. Computation was performed at 25 °C for all instructions. (B) The remaining registers after selective erasure are plotted. The query strand concentration was higher than the estimated register concentration; thus displacement achieved full erasure of registers. The green bars indicate the fraction of the read out registers with the corresponding barcode. Computation was performed at 40 °C for Instruction 1, and 25 °C for other instructions. For both (A) and (B), the registers were composed of chemically synthesized DNA.

Another feature of our random access scheme is the targeted deletion of selected data, which may be crucial for the secure management of sensitive information. For example, in medical records application, the information about a particular set of patients can be specifically removed. This contrasts with prior methods of erasure in which the erased information remains intact but is marked as erased (e.g., (*7*)). We demonstrate selective erasure by adding excess query strands, and reading out the values from the remaining mix. We observed that reads corresponding to the displaced registers were almost completely removed (Figure 4B, also Figure S11).

## Multiple Rounds of Computation

The promise of dynamic DNA storage in removing slow and expensive read-write cycles requires that the computation cycle can be applied multiple times. Toward this goal, we performed three rounds of Rule 110 computation on an input mix containing 5 initial values (Figure 5A). In the first round of computation, for all inputs, at least 83% of the read out registers updated correctly. In the second and third rounds of computation, the correct percent decreased more for some register values (e.g., 2nd column) than others (see Discussion). For the binary counting program, we took the input mix that underwent one round of computation and selective erasure (from Figure 4B), and performed another round of computation (Figure 5B). The correct value was present in at least 17% of all read out registers. Similar results were achieved for a register mix where a different barcode was erased (Figure S12). These experiments demonstrate the feasibility of not only multiple computational rounds, but the compatibility of selective retrieval, erasure and computation in the same register pool.

**Figure 5:**
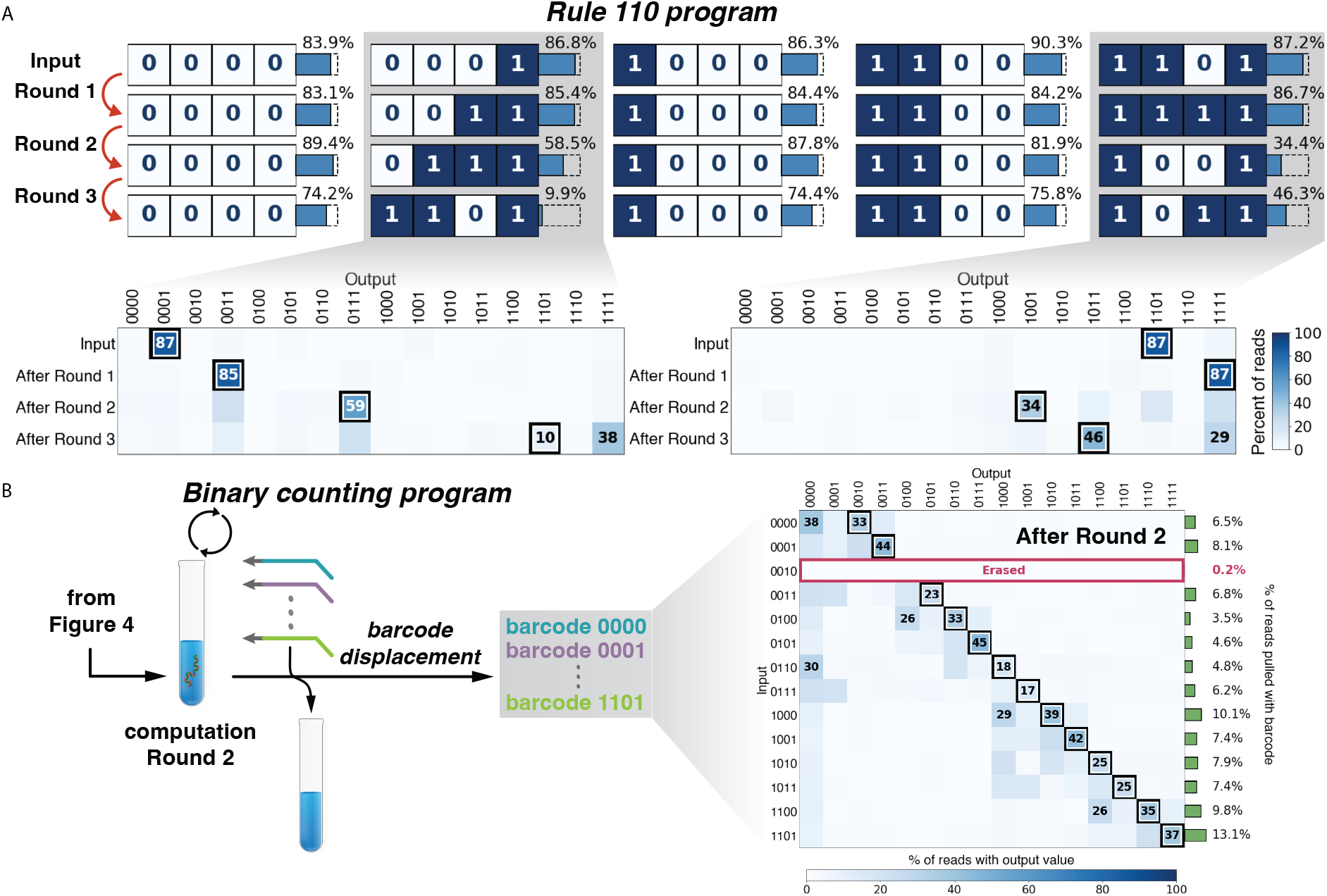
Multiple rounds of SIMD computation. (A) Three rounds of Rule 110 computation were performed on an input mix with 5 distinct initial values (columns). Lower panels show the distribution of output values for initial values 0001 and 1101. Computation was performed at 25 °C for all instructions. The reaction time for every instruction was 2 min except for instruction 2 which was 10 min. (B) The mix of inputs that underwent selective erasure from Figure 4B went through another round of binary counting computation. Computation was performed at 40 °C for Instruction 1, and 25 °C for other instructions. For (A) and (B), the registers were composed of chemically synthesized DNA.

## Discussion

SIMD||DNA bridges the fields of DNA computing and DNA storage by allowing computation directly on the storage medium, potentially allowing massively parallel processing on large scale databases. Since the first experiment using DNA to solve instances of NP-complete problems (*38*) in the 1990s, there have been more than twenty years of study of parallel DNA computing machines, with many models relying on enzymes to introduce covalent modification on DNA (*39–42*). However, these models are incompatible with current DNA storage paradigms, and other enzyme-free models have limits and implementation challenges (for example the sticker model (*43*) requires well-tuned binding affinities to allow for differential melting^2^).

Strand displacement is one of the most versatile mechanisms to modify chemically-encoded information (*17–24*), and theoretical work has shown that strand displacement can in principle perform Turing universal computation on information-storing polymers (*45*). Building on these insights we developed the framework of SIMD||DNA, implemented two algorithms (binary counting and Rule 110) and demonstrated random access, erasure, and multiple rounds of computation. Binary counting is a fundamental function in computer programming, and could be used to track events in applications such as updating medical records. Rule 110 demonstrates that, in principle, our paradigm is capable of executing arbitrary algorithms since any algorithm can be ‘compiled’ to Rule 110 (*34*). Additional efficient algorithms can be envisioned for bit shifting and parallel bubble sorting (*46*) (fundamental algorithms in computer science), as well as more efficient ways to ‘compile’ other algorithms (*47*). Algorithms can be recast to SIMD||DNA in many distinct ways; this diversity motivates the investigation of input encoding, running time, space usage, parallelism and randomization (see Section S2).

Crucially for scalability, SIMD||DNA ‘software’ can be run on different underlying ‘hardware’ choices. We constructed registers and achieved strand displacement computation for both SIMD||DNA programs using either chemically synthesized or naturally-occurring M13 DNA. Indeed, we show that large strand displacement systems (122 distinct displacement steps) can operate on naturally-occurring sequences without much sequence optimization—comparable to the most complex strand displacement systems using rationally designed sequences (*21*). Due to the key role that the M13 plasmid plays in structural DNA nanotechnology (*30, 33*), recent work has developed low-cost biological production of short single-stranded DNA with M13 subsequences (*48*). Thus it may now be possible to biologically produce both registers and instruction strands at scale, something that would finally allow dynamic information storage with low cost.

Importantly, we demonstrated high-throughput readout of the dynamically manipulated information through NGS sequencing. In almost all cases, the correct outputs had the highest fraction of all sequencing reads. In those few instances where there was only a substantial, but not majority, count for the right answer, it is likely that further refinement will yield higher accuracy; hypotheses about attendant errors and potential for improvement are discussed in Section S17. (See Section S15 for additional forms of product loss not captured by read counts, including loss of registers due to washing and incomplete ligation.)

Data storage in the nicking patterns does trade data density and stability for computability. Our current encodings in SIMD||DNA store data at a density of approximately 0.03 bit per nucleotide, a decrease from traditional storage schemes that encode information in the DNA sequence itself (theoretical maximum data density of 2 bits per nucleotide). In principle, data density can be increased by using different encoding schemes, such as allowing overhangs on the top strands to encode information. Additionally, compared to information stored in the DNA sequence, which could be stable for thousands of years (*2*), information stored in the pattern of nicks may be more prone to change since the patterns of nicks is more readily disrupted (e.g. via undesired 4-way branch migration between different registers, temperature fluctuations). Indeed, the temperature instability of topological DNA storage could enable easy and complete erasure of information through heating (*49*). For applications requiring longevity, however, in addition to using specialized materials for the protection of DNA (*50, 51*), it is possible to seal the nicks reversibly through light-induced photochemical ligation (*52*).

Several follow-up approaches could be used to further scale up SIMD||DNA. Recently developed Nanopore sequencing methods could potentially read information encoded in nicks and single-stranded gaps in double-stranded DNA directly in a high-throughput manner without PCR amplification (*53*). To increase both the yield and scale of computation, registers can be affixed to the surface of a microfluidic chip to achieve autonomous control of instruction strand addition and elution. Further, as discussed in Section S2.1, our programs in principle only require a small set of orthogonal domains (7 for Rule 110 and 8 for binary counting) independent of the length of the register. Reusing domains in this manner would also allow each instruction to be implemented with up to two distinct strands, potentially reducing chemical synthesis cost significantly.

SIMD||DNA ultimately adds a ‘wet’ CPU to the opportunities inherent in DNA encoding for ‘cold’ data storage. Such dynamic DNA storage could revolutionize the DNA storage architecture for applications that involve multiple computation cycles and parallel computation, especially since SIMD||DNA circumvents repeated sequencing and synthesis of oligonucleotides. The demonstrations that distinct strand displacement cascades can be read out in parallel, that long strand displacement cascades may not require extensive sequence design and can operate with naturally-occurring sequences, and that barcoded registers can be separately processed is important when we begin to consider DNA data storage beyond purely archival purposes—for example, for the storage and updating of medical records for specific patients. In particular, more advanced querying could be used to select a population of patients satisfying certain criteria (such as receiving a combination of diagnostics or treatment plans), and such algorithmic queries could be practically realized via SIMD||DNA programs that control the accessibility of the toehold for the query strand as the output of the computation (Figure S17).

## Acknowledgments

We thank Marc Riedel, Olgica Milenkovic, and Tonglin Chen for invaluable discussions. We also thank Marc Riedel for suggesting the analogy to Single-Instruction, Multiple-Data Computers. We thank the Genomic Sequencing and Analysis Facility at UT-Austin for providing sequencing services. We thank Cheulhee Jung and Bingling Li for sharing experience of working with magnetic beads, and David Doty for discussion and suggestion of using ViennaRNA.

## Authors contributions

B.W., C.C., and D.S. devised the project. B.W. and C.C. designed the programs and proved the correctness. B.W. designed the experiments, performed initial fluorescence experiments, and experiments for register preparation, computation, post-computation ligation and displacement. S.W. performed qPCR assays, post-computational library preparation, and data analysis. B.W. and S.W. prepared the figures. B.W. wrote the initial draft of the manuscript. All authors edited the paper. D.S. and A.E. provided guidance throughout the project.

## Data and materials availability

All materials are available upon request.

## Supplementary materials

Theoretical proofs and questions

Materials and Methods

Supplementary Text

Figs. S1 to S17

Tables S1 to S3

References (*54–63*)

## Footnotes

1 Single instruction, multiple data (SIMD) is one of the four classifications in Flynn’s taxonomy (*25*). The taxonomy captures computer architecture designs and their parallelism. The four classifications are the four choices of combining single instruction (SI) or multiple instruction (MI) with single data (SD) or multiple data (MD). SI versus MI captures the number of processors/instructions modifying the data at a given time. SD versus MD captures the number of data registers being modified at a given time, each of which can store different information. Our scheme falls under SIMD, since many registers, each with different data, are affected by the same instruction.

2 In a sense, we realize an extension of the sticker model envisioned by (*44*): “Recent research suggests that DNA ‘strand invasion’ might provide a means for the specific removal of stickers from library strands. This could give rise to library strands that act as very powerful read-write memories. Further investigation of this possibility seems worthwhile.”

