## Supporting Information for "Parallel molecular computation on digital data stored in DNA"

### Parallel molecular computation on digital data stored in DNA
### Supporting Information

Boya Wang[1¶,*], Siyuan S. Wang[2¶], Cameron Chalk[1], Andrew D. Ellington[2], and David Soloveichik[1,*]

[1]*Department of Electrical and Computer Engineering, University of Texas at Austin, Austin, TX 78712, USA*
[2]*Department of Chemistry and Biochemistry, Institute for Cellular and Molecular Biology, University of Texas at Austin, Austin, TX 78712, USA*

*To whom correspondence should be addressed;
 (B.W.); (D.S.)

**S6 Characterizing register locations on the M13mp18 plasmid for subsequent experiments** **20**

**S7 Binary counting: single instruction single data results for M13.8 and M13.3 registers** **22**

**S8 Binary counting: single instruction multiple data results for M13.7, M13.8 and M13.9 registers** **23**

**S9 Binary counting: single instruction multiple data initial values for M13.7 and M13.9 registers on the same M13 molecules** **24**

**S10 Rule 110 boundary implementation** **25**

**S11 Rule 110: single instruction multiple data** **26**

**S12 Random-access specificity and selective erasure** **27**
S12.1 Random access . . . 27
S12.2 Selective erasure . . . 29

**S13 Multiple rounds of binary counting program** **30**

**S14 Washing efficiency** **31**

**S15 Investigating register loss during SIMD||DNA computation** **32**

**S16 Investigation of instruction completeness through fluorescence** **34**

**S17 Hypotheses about error** **35**

**S18 The number of strand displacement steps performed on naturally-occurring sequences** **36**

**S19 Algorithmic query** **37**

**S20 Sequences** **38**

#### S1 SIMD||DNA Programs

We consider sequences of instructions, called *programs*. We design programs for binary functions over $n$ bits $f : \{0,1\}^n \to \{0,1\}^n$ so that, given a register encoding any input $s = \{0,1\}^n$, after applying all instructions in the program sequentially as in Figure 1B, the resulting register encodes $f(s)$. Here we give our two programs: binary counting and simulation of elementary cellular automaton Rule 110.

##### S1.1 Binary counting

The counting program computes $f(s) = s + 1$. Binary counting is captured by changing all the 1s to 0 from the least significant bit to more significant bits until the first 0, and changing that 0 to 1. All the bits more significant than the rightmost 0 remain the same. For example, $f(1011) = 1100$, and $f(1000) = 1001$. In the case of overflow, we rewrite the register to all 0s. In other words, on inputs of all 1s, we output all 0s: $f(1111) = f(0000)$.

Binary counting allows one SIMD||DNA program to move data through a number of states exponential in the size of the register. We consider this a requirement of any useful data storage/computation suite: if instead not all configurations of the register were reachable from some initial configuration via some program, then the useful density of the storage would be reduced.

The full program is in Figure 2. Each state-0 cell is fully covered by two strands, with one covering the first three domains and the other one covering the last two domains. Each state-1 cell is fully covered by two strands, with one covering the first two domains and the other one covering the last three domains. One extra domain is included to the right of the rightmost cell which is used to initiate displacement. The program contains seven instructions. It erases all the 1's in between the rightmost cell and the rightmost state-0 cell at Instructions 1 and 2, and changes those cells to 0 at Instructions 4 and 5. It marks the rightmost state-0 cell at Instruction 3, and change the marked state-0 cell to state 1 at Instructions 6 and 7.

To prove correctness, we first argue that all the 1's from the least significant bit to the rightmost 0 update to 0. Then we argue that rightmost 0 updates to 1. Assume the bit string has length $n$ (and so the least significant bit is at cell $n$), and the rightmost 0 is at cell $m$ ($m \leqslant n$). We let $j_k$ denote the $k$th domain on cell $j$ (from left to right).

In the main text, we defined the SIMD||DNA model with the assumption that all cells are orthogonal in sequence since this best corresponds to our experimental implementation. However, this assumption is not necessary for the proof of correctness of our programs, and we allow all cells to share the same sequences, only assuming that each domain within a cell is orthogonal (i.e., $j_k$ can be the same sequence as $j'_k$ but must be orthogonal from $j_{k'}$). Additionally, the extra domain to the right of the rightmost cell is orthogonal to all other domains. (See Section S2.1 for a discussion of domain orthogonality and whether algorithms rely on it.)

**Claim S1.1.** *All state 1 cells to the right of the rightmost 0 cell change to a 0.*

*Proof.* Instruction 1 initiates a series of sequential reactions from the least significant bit $n$ to the rightmost 0. First the instruction strand with overhang domain $a$ displaces the strand covering domains $n_4$ and $n_5$. If the least significant bit is 1 ($m < n$), the domain $n_3$ becomes unbound after this displacement reaction. Then the domain $n_3$ serves as an open toehold to initiate another displacement reaction with the instruction strand with overhang domain $b$. Similar displacement reactions proceed until cell $m$. By assumption, cell $m$ is a state-0 cell, so the domain $m_3$ will not be open after the displacement step, and thus the displacement cascade stops. Then the strands added in Instruction 2 detach the strands from Instruction 1, leaving the cells from the $(m+1)$th bit to the $n$th bit free. In Instruction 3, every applied instruction strand from cell $m+1$ to $n$ attaches to the register. Instruction 4 shifts those strands added in Instruction 3 one domain to the left, which opens toeholds for the cooperative displacement in Instruction 5. After those cells change to state-0 in Instruction 5, the strands added in Instruction 6 and 7 do not change them, so they remain in state 0. □

**Claim S1.2.** *The rightmost state* 0 *cell changes to a* 1*.*

*Proof.* Instruction 1 initiates a series of sequential reactions from the least significant bit to the rightmost 0 at cell $m$. The domain $m_3$ will not be open after the instruction strand displaces the strand covering domains $m_4$ and $m_5$ and no more strand displacement can proceed to the left. Then the strands added in Instruction 2 detach the strands from Instruction 1, leaving the domains $m_4$ and $m_5$ free. The strands added in Instruction 3 serve as two purposes: (1) They correspond to one of the strands representing state 1. Thus they help cell $m$ to transition to state 1 and they partially displace the strand at domain $m_3$. (2) They serve as a placeholder by binding at domains $m_4$ and $m_5$ to prevent cell $m$ from being modified in Instructions 4 and 5. Instruction 6 detaches the strand originally bound from domain $m_1$ to $m_3$, leaving the domains $m_1$ and $m_2$ open. In Instruction 7, the instruction strand attaches to the register at domains $m_1$ and $m_2$, which completes the state changing from 0 to 1. □

**Claim S1.3.** *The cells to the left of the rightmost state* 0 *cell stay the same.*

*Proof.* Note that no open toeholds are exposed at cells to the left of cell $m$, and the displacement cascade does not pass to the left of cell $m$, thus no changes are made to the states of those cells. □

#### S1.2 Cellular automaton (CA) Rule 110

An elementary one-dimensional cellular automaton consists of an infinite set of cells $\{\ldots, c_{-1}, c_0, c_1, \ldots\}$. Each cell is in one of two states, 0 or 1. Each cell changes state in each timestep depending on its left and right neighbor's states. Rule 110 is defined as follows: the state of a cell at time $t+1$, denoted $c_i(t+1)$, is $f(c_{i-1}(t), c_i(t), c_{i+1}(t))$, where $f$ is the following:

$$
\begin{aligned}
f(0,0,0) &= 0 \qquad & f(1,0,0) &= 0\\
f(0,0,1) &= 1 \qquad & f(1,0,1) &= 1\\
f(0,1,0) &= 1 \qquad & f(1,1,0) &= 1\\
f(0,1,1) &= 1 \qquad & f(1,1,1) &= 0
\end{aligned}
$$

Note that a simple two-rule characterization of $f$ is as follows: 0 updates to 1 if and only if the state to its right is a 1, and 1 updates to 0 if and only if both neighbors are 1. This characterization is useful for proving correctness of the program [34].

The instructions implementing one timestep evolution are shown in Figure S1. Each state-0 cell is fully covered by two strands, one of length three and one of length two. Each state-1 cell is partially covered by a length-five top strand and has an open toehold at the leftmost domain. The program consists of six instructions. The program first marks the string "01" (Instruction 1)—here, the 0 will change to 1 later. Then it erases the internal 1's in any string of at least three consecutive 1's (Instructions 2 and 3). These are the 1's with two neighboring 1's, which should be updated to 0, so the program fills in the empty cells with 0 (Instruction 4). Finally it removes the markers from Instruction 1 and changes previously marked 0's to 1's (Instructions 5 and 6).

We claim that this program enforces the two-rule characterization of Rule 110. We first argue that 1 updates to 0 if and only if both neighbors are 1. Then we argue that 0 updates to 1 if and only if the state to its right is a 1. As for binary counting, let $i_k$ denote the $k$th domain on cell $i$ (from left to right). As for binary counting, all cells can share the same sequences, but we assume that each domain within a cell is orthogonal.

**Claim S1.4.** *A cell $i$ initially in state 1 updates to a 0 if cells $i+1$ and $i-1$ are initially 1.*

*Proof.* During Instruction 1, the instruction strands cannot displace the strands in state-1 cells. In Instruction 2, the strand on cell $i$ is displaced cooperatively only if the toeholds on both the left and the right side of the strand are open. By assumption, cell $i+1$ is a 1, so the toehold immediately to the right of cell $i$, $(i+1)_1$, is free. Since cell $i-1$ is in state 1, domain $i_1$ is not covered after Instruction 1 ($i_1$ would be covered if cell $i-1$ were 0). Thus the strand on cell $i$ can be displaced by the instruction 2 strands. In Instruction 3, the instruction 2 strands in cell $i$ are detached, so every domain in cell $i$ is free. Then in Instruction 4 we attach the strands corresponding to a state 0, updating cell $i$ to 0. Instructions 5 and 6 do not introduce any instruction reaction on cell $i$, so cell $i$ remains in state 0. □

**Claim S1.5.** *A cell $i$ initially in state 1 stays in state 1 if either cell $i+1$ or $i-1$ is initially 0.*

*Proof.* During Instruction 1, the instruction strands cannot displace the strands in state-1 cells. In Instruction 2, the strand on state-1 cells is displaced cooperatively only if the toeholds on both the left and the right side of the strand are open. By assumption that the left or right cell is a 0, the toeholds required for this, $i_1$ or $(i+1)_1$, will be covered: First consider that cell $i-1$ is a 0. Then in Instruction 1, the instruction strand displaces one strand at cell $i-1$ and covers the toehold $i_1$. On

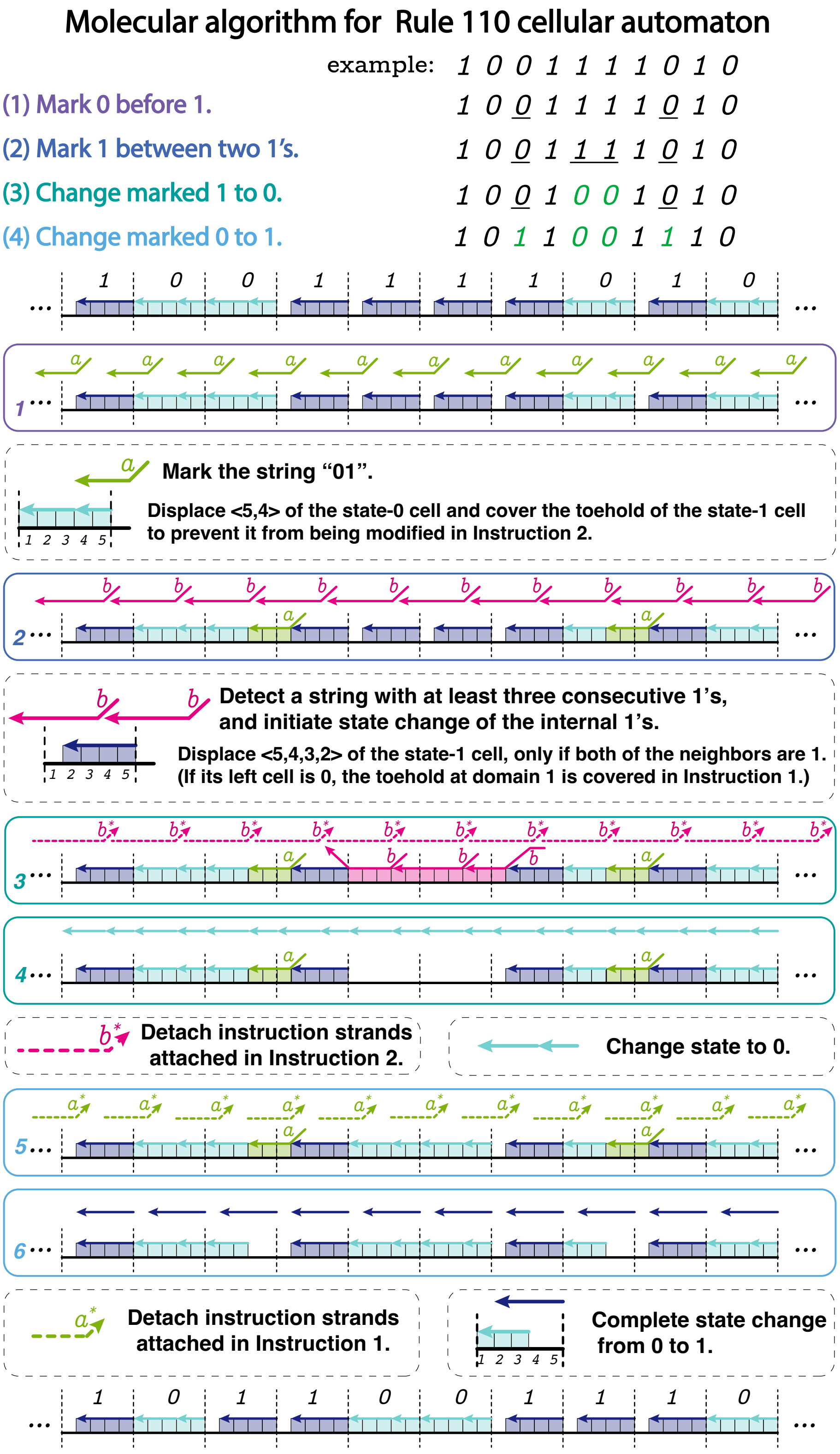

**Figure S1:** The program implementing one timestep of Rule 110 shown on an example register. The top register shows the initial state of each cell. After 6 instructions, the register updates to the state shown at the bottom. Strand colors have three information categories: state 1 (dark blue), state 0 (light blue), intermediates (other colors). Solid boxes show the instruction strands and the state of the register before the strands are applied. Dashed boxes explain the logical meaning of the instructions. The overhang domains $a$ and $b$ are orthogonal to other domains vertically aligned with them.

the other hand, if cell $i+1$ is $0$, then domain $(i+1)_1$ is covered since strands for the $0$ state cover all the domains in that cell. So if either neighbor state is $0$, Instruction 2 does not displace the strand on cell $i$. Then note that Instructions $3$ and $4$ do not introduce any instruction reaction at cell $i$. The instruction $5$ strands detach the instruction $1$ strands if cell $i-1$ is $0$, freeing the toehold at $i_1$ and recovering the state-1 cell. Instruction 6 does not change the state-1 cell. □

**Claim S1.6.** *A cell $i$ initially in state $0$ updates to a $1$ if cell $i+1$ is initially a $1$.*

*Proof.* Since cell $i+1$ is in state $1$, the toehold at domain $(i+1)_1$ is available for the instruction strand in Instruction $1$ to bind, and the rightmost strand on cell $i$ is displaced. Then note that Instructions $2$ through $4$ do not introduce any instruction reaction at cell $i$. In Instruction $5$, the instruction strand from Instruction $1$ is detached, freeing domains $i_4$ and $i_5$. In Instruction $6$ the instruction strand binds at domains $i_4$ and $i_5$ and displaces the strand at cell $i$. So after Instruction $6$, cell $i$ is in state $1$. □

**Claim S1.7.** *A cell $i$ initially in state $0$ stays in state $0$ if cell $i+1$ is initially a $0$.*

*Proof.* Simply note that for any instruction, no instruction reaction on cell $i$ occurs. So cell $i$ stays in state $0$. □

These four claims combined verify that the two-rule characterization given at the beginning of this section is satisfied, so the instructions implement one timestep evolution of Rule 110.

Note that the Rule 110 simulation invokes two sources of parallelism. Instruction strands are applied to all registers in parallel, and every cell within a register can update concurrently (see also Section S2.5).

Also note that Rule 110 is defined only for an infinite set of cells or a circular arrangement of finitely many cells. For a finite set of cells arranged linearly, one must define boundary conditions for updating the leftmost and rightmost cells. Boundary conditions can be constant or periodic. For space-bounded computation by Rule 110, it suffices to set periodic boundary conditions based on the periodic initial condition of the CA given in [34]. These periodic boundary states can be implemented by periodic instructions.

Theoretically, SIMD||DNA's in-memory computation model is as powerful as any other space-bounded computing technique. In other words, our space-bounded simulation of Rule 110 immediately gives that any computable function can be computed by a SIMD||DNA program, if the required space is known beforehand.

### S2 Theoretical discussion

#### S2.1 Orthogonal domains

A common assumption for correctness in strand displacement systems is that domains are *orthogonal*, meaning two domains which are not fully complementary do not bind. In experiments, enforcing this assumption requires specialized sequence design. Further, for any fixed domain length, the space of orthogonal domains is limited, restricting the scalability of the system. As shown by their proofs of correctness (see Section S1), our programs only require a small number of unique domains, independent of the register length. In principle, this allows our programs to use a constant set of orthogonal domains (8 for the binary counting program and 7 for the Rule 110 program), simplifying the sequence design problem for experimental implementation. Reusing a fixed set of domains across cells can also potentially reduce synthesis costs if the same instruction strands can be used to compute on every cell (see Uniform and non-uniform instructions below).

#### S2.2 Uniform and non-uniform instructions

We say that an implementation of a SIMD||DNA program is *uniform* if increasing the number of cells in the register does not increase the number of different instruction strands. (We use the term uniform in the computer science sense describing an algorithm that applies on inputs of arbitrary length.) In contrast a *non-uniform* implementation requires more different instruction strands. To create a uniform program and enable a fixed set of instruction strands to suffice for arbitrarily long registers, we need to share sequence space in different parts of the register; e.g., in the extreme case by making all cells have exactly the same sequence (see Orthogonal domains above). Uniform implementations can potentially incur significant cost savings in DNA synthesis costs, as well as simplify experiments since fewer strands are involved. On the other hand, it is possible that relying on non-uniformity might increase the computational power of SIMD||DNA—for example, instructions which target specific cells may give additional flexibility and decrease the overall number of instructions required. The difference in computational power due to non-uniformity remains open.

As we have argued, our binary counting and Rule 110 programs are compatible with uniform implementations, potentially reducing the cost of instruction strand synthesis for large registers (although we did not experimentally demonstrate a uniform implementation).

#### S2.3 Undesired reaction with waste products

In the idealized SIMD||DNA model, we assume that displaced top strands (*waste*) do not subsequently engage in strand displacement because they are in much lower concentration than the instruction strands. Nonetheless, in reality these waste products could interact further within the system, even with relatively slower kinetics. Interestingly, in the two programs we designed in the paper, we conjecture that besides the reverse of the intended reaction (in the case of toehold exchange), the waste products and registers cannot react, and therefore our programs are robust to this type of error. Further, as shown in Figure S13, it is reasonable to assume that washing removes all the waste products and thus they do not accumulate between instructions.

#### S2.4 Determinism and nondeterminism

Our programs are designed with *deterministic* instructions: given one state of the register, after adding the instruction strands, the register is supposed to change to one specific state. Deterministic instructions make it easy to design, predict, reason about, and compose the programs. In contrast to deterministic instructions, one could also construct *nondeterministic* instructions by introducing nondeterminism to the updates of the cells. For example, consider an empty cell with domains $\langle 3^*, 2^*, 1^* \rangle$, and add instruction strands $\langle 1, 2 \rangle$ and $\langle 2, 3 \rangle$. (The bracket notation $\langle\ \rangle$ specifies a strand by listing the constituent domains from 5′ to 3′-end.) Either the first or second strand can bind, but since they displace each other, only one will remain after elution. The probability of which strand remains depends on its relative concentration. In principle, applying nondeterministic instructions allows for implementation of randomized algorithms and simulation of nondeterministic computing machines.

#### S2.5 Running time

The running time of a program depends on two factors: running time per instruction and the number of instructions. The running time per instruction depends on whether the instruction updates the cells through *parallel* or *sequential* reactions. In general, instructions are capable of acting on each cell within each register in parallel. Yet, Instruction 1 of the binary counting program does not have this source of parallelism. A first reaction (displacement) must occur on the rightmost cell prior to a second reaction occurring on the second cell, which must occur prior to a third reaction on the third cell, and so on. Thus, this instruction with sequential reactions loses the speedup given by independent instruction reactions occurring in parallel on each cell within a register. Besides the running time per instruction, the larger the number of instructions per program, the more complex is the experimental procedure. This motivates studying the smallest number of instructions required to achieve a computational task.

#### S2.6 Universal computation

Our registers as proposed are restricted to a finite number of cells. So although Rule 110 on an infinite arrangement of cells can simulate an infinite-tape Turing machine, our scheme is only capable of space-bounded computation. To claim that a system is capable of universal computation, it is required that the data tape—in our case, the number of cells—can be extended as needed as computation proceeds. Since our program consists of uniform instructions, domain orthogonality is only required within a cell. Therefore, in principle, the register can be extended indefinitely during computation without exhausting the space of orthogonal domains. The register's length could perhaps be extended by merging bottom strands with top strand "connectors".

#### S2.7 Space-efficient computation

Although Rule 110 is Turing universal, computing functions through simulation of a Turing machine by Rule 110 does not make use of the full power of SIMD||DNA. First of all, while simulation of a Turing machine by Rule 110 was shown to be time-efficient [54], it is not space-efficient. Precisely, simulating a Turing machine on an input which takes $T$ time and $S \leq T$ space requires $p(T)$ time and $p(T)$ space (where $p(T)$ is some polynomial in $T$). However, Turing machines can be simulated time- and space-efficiently by one-dimensional CA if the automaton is allowed more than two states [55]. Simulating larger classes of CA is a promising approach to space-efficient computation in this model, since our Rule 110 simulation suggests that CA are naturally simulated by SIMD||DNA programs.

#### S2.8 Equalizing encodings

Our two programs use different schemes for encoding binary information in a register. Using some universal encoding would allow applying different consecutive computations to the same registers. Alternatively, we could design programs to inter-convert between different encodings. The reason for suggesting this alternative is that unlike classical machines acting on bits, in SIMD||DNA the way a bit is encoded affects how it can be changed by instruction reactions. For example, in the binary counting program, the encoding ensures that no toeholds except for the rightmost domain are open on the register, which is used to argue correctness. Alternatively, in the Rule 110 program, toeholds must be available throughout the register to achieve the parallel cell updates required by CA. Therefore having one encoding which implements these two different functions seems difficult.

### S3 Sequence design

#### S3.1 Design sequence space for artificial sequences

We generated the sequences for the binary counting program following the procedure described below. The sequences for the binary counting program were directly mapped to the Rule 110 program, and in fact, some of the instruction strand sequences were shared between these two programs. Considering that instruction 1 has a long strand displacement cascade with most of the reactions being reversible, we designed the middle domain (the 3rd domain) in each cell to be 6-nt long to promote strand dissociation, and all other domains were 7-nt long.

1. We generated a pool of length-6 and length-7 domains, with each nucleotide position having an equal probability of A, T, G. The ATG alphabet was used for all the top strands on the register [56, 27], and the complementary ATC alphabet was used for the bottom strands. This was done to minimize possible secondary structures of some intermediate states of the registers especially when there exist a long unbound region of the bottom strand (e.g. the register after instruction 4). The domains must satisfy the following requirements:
   (a) No more than 4 consecutive A's; no more than 4 consecutive T's;
   (b) To reduce the effect of fraying [17, 57], both the first two and the last two nucleotides contain exactly 1 G;
   (c) To ensure the domains are sufficiently strong, the total number of G in each domain is either 2 or 3.
2. We randomly took domains from the pool and assembled them into a long strand (the bottom strand). To avoid synthesis errors, the long strand was also restricted to satisfy the requirement of no more than 4 consecutive A's or T's, no 4 or more consecutive G's, or no more than 7 consecutive A/T's. Among the candidate sequences, we chose the one with the size of the longest repeated segment being as small as possible.

#### S3.2 Assignment of register addresses for naturally-occurring DNA (M13)

We first passed a very simple check described below on the M13mp18 sequence to filter out a few regions. Then we assigned candidate sequences in those regions based on the binding strength of each domain and assembled the domain sequences into strand sequences.

Starting from location 1, we chose a sequence window of 265 nt and screened the sequence of the entire plasmid. The size of 265 was chosen to accommodate the length limit for paired end reads on the Illumina MiSeq platform. We left out sequence windows with (1) 4 or more consecutive G's, and 4 or more consecutive C's to avoid the synthesis error on the corresponding top strands; and (2) 12 or more consecutive A/T's to avoid regions with extremely weak bonds. We selected 9 sequence windows at the following locations: $\{796, 1060\}$, $\{1326, 1590\}$, $\{1591, 1855\}$, $\{2651, 2915\}$, $\{2916, 3180\}$, $\{3181, 3445\}$, $\{3446, 3710\}$, $\{4241, 4505\}$, $\{4771, 5035\}$. Luckily, none of these windows intersects with the region $\{5500, 6000\}$ which is known to form several strong hairpins [33].

To assign sequences to domains, we noticed that some domains—the third and the fifth domain (Figure S2) from left to right in a cell in the binary counting program—need to be weak since they need to serve as the dissociation toehold, while the strength of other domains are allowed to

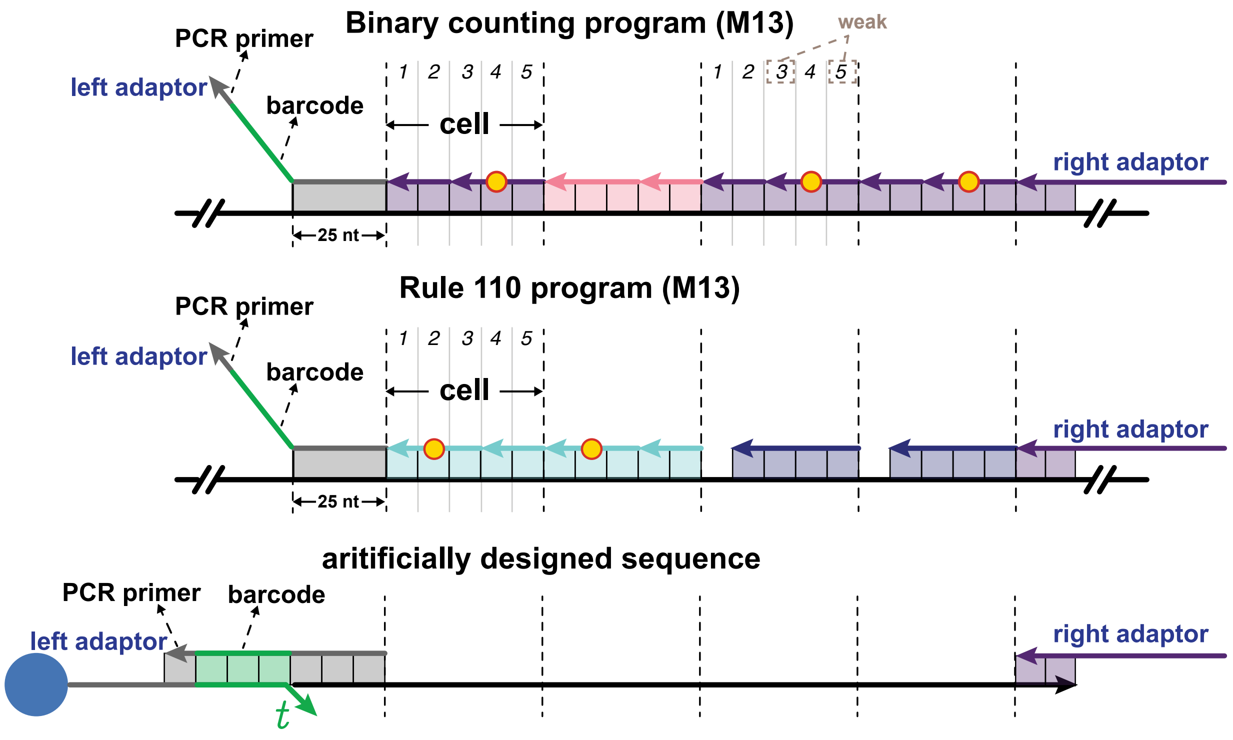

**Figure S2:** Domain layout in registers using naturally-occurring (M13) sequences and artificially designed sequences.

vary. We used NUPACK 3 [16] to calculate the free energy of binding for a domain with its complement. We chose 3 sets of binding stabilities calculated with NUPACK command `/bin/energy -T 25 -material dna`: weak (between −12 kcal/mol and −8.5 kcal/mol), medium (between −17 kcal/mol and −14 kcal/mol) and strong (between −23 kcal/mol and −20 kcal/mol). (Note that the free energy reported by NUPACK uses water concentration with *mole fraction* as the unit instead of molar and thus an appropriate correction was applied [16].) Starting at position 26, the sequences for the domains were assigned according to the binding energy of each domain. The first 25 nt in each region is used to connect the barcode and PCR primer to the selected register sequence.

The Rule 110 program was only implemented with weak binding strength registers (M13.1, M13.2 and M13.3) since that allowed us to reuse some strands from binary counting for Rule 110. (Sharing strands for instruction 1 and 2 in the binary counting program and strands for instruction 2 and 3 in the Rule 110 program requires domains 1, 3, and 5 in cells to be weak in order to readily dissociate.)

#### S3.3 Design of the adaptor region: PCR primers, barcodes, and toeholds

The adaptors were designed to accommodate post-computation processing such as displacement and PCR amplification. Every register has a left and a right adaptor, with regions for the forward and the reverse PCR primer. The left adaptor also contains the barcode sequence encoding the information for the initial value of a register. There are in total 16 unique barcodes corresponding to 16 initial values for a 4-bit register.

The key consideration for barcodes is that they need to be as orthogonal as possible. We selected the sequences for barcodes from the Seesaw sequence pool [18] where there is a collection of 15-nt domains with at least 30% of difference of bases and the shared sequences between each domain are at most 5 nt. For random-access experiments performed on artificially designed sequences, to ensure that the binding strength between the biotin-labelled strand and the left adaptor strand is sufficiently strong at different temperatures, each barcode is composed of two 15-nt domains drawn from the Seesaw pool. For registers constructed on M13, the barcodes were 15-nt long and likewise chosen from the Seesaw pool.

For random-access, a displacement step is required during the post-computation process. The 7-nt toeholds for displacement were generated using the A, T, C alphabet with the following requirements: (1) Each toehold contains no more than 4 consecutive A/T's and no more than 3 consecutive C's; (2) The percentage of C is between 30% to 60%; (3) The hamming distance between each toehold is greater than 2; (4) The binding energy between any of the toehold pairs is greater than $-2.9$ kcal/mol calculated using the *RNAduplex* function (`RNAduplex -P dna_mathews1999.par -T 25 --noGU`) of ViennaRNA [58]. For each unique initial value, a pair of barcode and toeholds were randomly chosen for the biotin-labelled strand and the displacing strand.

##### S3.4 Assignment of mismatches

For the secondary encoding, we assigned mismatches (relative to the bottom strand) to certain domains on some top strands so that $0$ and $1$ can be differentiated after ligation. To determine whether to assign mismatches at strands representing $0$ or representing $1$, we chose the strands not invading any other strands (any mismatch is on the displaced strand) throughout the entire program. If a bit is represented by two strands, the longer strand was preferred due to a larger sequence space.

According to the relative thermodynamic energy of single-base change [37], we chose to change either A or T to G, since they introduce significantly lower disturbances compared to a perfectly matched pair. In screening the sequences of the chosen strands, we preferred to have a mismatch with a smaller change of energy, and at least 4 nt away from the end of a strand (to prevent more fraying at the end of the double-stranded region). We also ensured that no strand has 4 or more consecutive G's (to prevent problems with synthesis).

### S4 Materials and methods

#### S4.1 DNA oligonucleotides

The M13mp18 single-stranded DNA was ordered from NEB (# N4040S). All other DNA strands were synthesized by Integrated DNA Technologies (IDT). For artificially designed sequences, the bottom strands were ordered as PAGE purified Ultramers; the unlabeled oligos for 4-bit registers were ordered PAGE purified; and the fluorophore or phosphate labeled oligos were ordered HPLC purified. For M13 related experiments, the oligos shorter than 60 nt were ordered HPLC purified and in plate; and the oligos longer than 60 nt were ordered PAGE purified.

For oligos ordered in dry form in tubes, we suspended them in nuclease-free water (not DEPC treated), and then quantified the absorabnce at 260 nm using NanoDrop. The concentration ($c$) was calculated as $c = [\text{Absorbance}]/e$, where $e$ is the extinction coefficient provided by IDT. Usually, the nominal stock concentration for single-stranded DNA was $\sim$100 $\mu$M. The oligos delivered in IDT plates were ordered to be normalized to 100 $\mu$M in IDTE pH 8.0 buffer.

#### S4.2 Register preparation

##### S4.2.1 Register Anneal

**Registers with artificially designed sequences.** The bottom strand and all the top strands were mixed and then annealed with $10\%$ excess of top strands. The annealing buffer was TE/Na$^+$ (1 M) buffer ($0.04$ M Tris, $1$ mM EDTA, $1$ M NaCl, pH 8.0, $0.01\%$ Tween 20). The annealing process was performed in a PCR thermocycler: DNA strands were incubated at $95$ °C for $5$ minutes and then slowly cooled down with rate $0.1$ °C/min to $20$ °C.
**Registers with M13 sequence.** The M13 registers were assembled with top strands at $5$ times excess compared with the M13 single-stranded DNA. All the DNA strands were incubated at $95$ °C for $5$ minutes and then slowly cooled down with rate $0.1$ °C/s to $20$ °C.

##### S4.2.2 Register attachment to magnetic beads

The Dynabeads MyOne Streptavidin C1 magnetic beads were purchased from Invitrogen (# 65001). The 96-well super ring magnet separator plate used for elution was purchased from Permagen (SKU:T480). The magnetic beads were first vortexed for 30 sec for suspension. Then $5$ $\mu$L 10 mg/mL beads were transferred to a tube and washed twice with the TE/Na$^+$ (1 M) buffer. The washed beads were incubated with the annealed register (25 $\mu$L at concentration 1 $\mu$M for registers with artificially designed sequences, and 14 nM for M13 registers) for 25 min. The beads were then washed twice by the washing buffer TE/Na$^+$ (0.5 M) buffer ($0.04$ M Tris, $1$ mM EDTA, $0.5$ M Na$^+$, pH 8.0, $0.01\%$ Tween 20) to remove the excess unlabeled register. (The conditions for the washing step were decided based on the results in Section S14.) Finally the beads were suspended with $25$ $\mu$L washing buffer. The register stock concentration was approximately $250$ nM, estimated based on bead capacity.

#### S4.3 Computation

Every computation step had registers with approximate concentration of $50$ nM at a total volume of $25$ $\mu$L. For SIMD experiments with multiple inputs, each input was in equal concentration.

For the binary counting program, the concentrations of the instruction strands are listed in Table S1. The reaction temperatures for Instruction 1 varied from 25 °C to 50 °C. The reaction temperature for all other instructions was 25 °C. After incubating for 10 min, the registers attached to magnetic beads were washed twice by the washing buffer.

| Instruction | Concentration |
|---|---|
| 1 | 3 $\mu$M |
| 2 | 0.5 $\mu$M |
| 3 | 0.5 $\mu$M |
| 4 | 3 $\mu$M |
| 5 | 1 $\mu$M |
| 6 | 0.5 $\mu$M |
| 7 | 0.5 $\mu$M |

**Table S1:** Concentrations of instruction strands for the binary counting program.

For Rule 110 program, the concentrations for instruction strands are listed in Table S2. The reaction temperature for all the instructions was 25 °C. Unless specified elsewhere, the reaction time was 10 min, and then the magnetic beads were washed twice by the washing buffer.

| Instruction | Concentration |
|---|---|
| 1 | 0.5 $\mu$M |
| 2 | 3 $\mu$M |
| 3 | 0.5 $\mu$M |
| 4 | 0.5 $\mu$M |
| 5 | 0.5 $\mu$M |
| 6 | 0.5 $\mu$M |

**Table S2:** Concentrations of instruction strands for the Rule 110 program.

#### S4.4 Post-computation processing

##### S4.4.1 Ligation and preparation for amplification

**Seal strand addition.** For the Rule 110 program, seal strands (25 nM) were mixed with registers for 10 mins at 25 °C prior to adding adaptor strands.

**Adaptor strand addition.** The right adaptor strand (0.5 $\mu$M) was mixed with registers after computation and the mixture was incubated for 10 min at 25 °C. The beads were then washed twice by 1× T4 ligase buffer. The 1× T4 ligase buffer was prepared by diluting the 10× T4 ligase buffer purchased from NEB (# B0202S) with the nuclease-free water and mixing with Tween 20 to reach 0.01%.

**Ligation.** After addition and elution of the adaptor strands, 150 units/$\mu$L of T7 ligase (NEB # M0318S) were incubated with the registers at 25 °C for 30 min. Then the products were washed twice with 1× T4 ligase buffer. (T4 ligase buffer is compatible with T7 ligase.)

**Register displacement from the magnetic beads with query strands.** The query strands were mixed with the ligated products at 25 °C for 10 min. The supernatant was transferred to a new tube and inactivated by heat for 10 min at 65 °C. The concentrations for the query strands were: 20 nM each for Rule 110 random-access experiments and yield investigation; 30 nM each for Rule 110 multiple-rounds-of-computation experiments; 50 nM each for binary counting multiple-rounds-of-computation experiments and random-access experiments.

###### S4.4.2 Library preparation

Prior to amplification, the displaced product was quantified by qPCR with the LightCycler 96 instrument (Roche). Reaction mixtures contained 2.5 $\mu$L of the displaced product, 500 nM each of forward and reverse PCR primers with *NGS adaptors* and *sample-specific NGS barcodes*, 400 $\mu$M dNTP, 1× EvaGreen intercalating dye (Biotium #31000), 0.4 U/$\mu$L Q5 DNA polymerase (NEB #M0491S), 1× Q5 Reaction Buffer (NEB). qPCR was performed on the displaced products using the following protocol: initial melting at 98 °C for 3 minutes, followed by 30 cycles of amplification with melting at 98 °C for 30 sec, annealing at 67 °C for 30 sec, and extension at 72 °C for 30 sec (measurement taken), followed by a final extension at 72 °C for 3 minutes (measurement taken). Once the quantitation cycle ($C_q$) of each sample was determined, PCR was repeated using the same thermocycling protocol in a thermocycler as described for qPCR, except that EvaGreen dye was replaced with nuclease-free water and the number of cycles was set to be $C_q + 5$ for each sample to minimize the amplification of side products. After PCR, equivalent amounts of each PCR product were pooled together and loaded on a 1.8 % NuSieve GTG agarose gel (Lonza #50081). After running the gel, a QIAquick PCR & Gel Cleanup Kit (Qiagen #28506) was used for purification per manufacturer's instructions using the included buffers with the following exceptions: gel fragments were incubated in Buffer QG for at least 20 minutes at 60 °C (instead of 10 minutes at 50 °C), and the column containing products were washed 3 times using Buffer PE (instead of once). The final samples were eluted in nuclease-free water and diluted to a concentration of 5 ng/$\mu$L as measured by Nanodrop.

###### S4.4.3 Next-generation sequencing

Sample libraries were sequenced for 2x261 cycles using Illumina MiSeq 2x250 paired end reagent kits (v2). Because the SIMD||DNA products share the sequence space and exhibit very low base diversity, it was necessary to artificially supplement base diversity to avoid sequencing failure. We added a sample library prepared from HeLa genomic DNA (NEB #N4006) and PhiX genomic DNA (fiducial markers for imaging added by the sequencing core facility). For sequencing runs consisting of more than 30 % SIMD||DNA products, this genomic DNA library was added to a fraction of 50 % of all reads.

##### S4.5 Sanger sequencing

We prepared samples for Sanger sequencing using the same PCR and clean up protocol described above. To improve the quality of the sequencing reads and increase confidence in the base calls, we sequenced samples with both forward and reverse primers.

#### S4.6 Fluorescence experiments

Fluorescence experiments were measured on the BioTek Synergy H1 multi-mode microplate reader. The low volume NBS (non-binding surface) 384 well plates with clear flat bottom were used, purchased from Corning corporation (# 3544). The excitation wavelength was 577 nm and the emission wavelength was 608 nm. The excitation bandwidth was fixed at 9 nm and the emission bandwidth was fixed at 20 nm. Data points were taken every half minute for two minutes, since beads precipitated over time affecting fluorescence signal.

### S5 Data analysis

#### S5.1 Sanger sequencing

Sanger sequencing traces were aligned to the expected SIMD||DNA product sequence using the "Map to Reference" feature in Geneious 2020.0.5. We determined the computation results using the composition of the base call at the nucleotide positions of interest ("variable nucleotide position"). Since the products could contain registers with and without mismatches at the variable nucleotide position, Sanger sequencing results could show mixed signal traces at the locations. In these cases, we interpreted the height of base call peaks in the raw trace to represent the relative proportions of each base in that population. Thus, although Sanger sequencing is intended for use with homogeneous samples to produce qualitative sequencing outputs, it can nonetheless indicate the relative amount of samples at the variable nucleotide position.

#### S5.2 Next-generation sequencing

Python code used for analysis of the NGS data is available via GitHub (`https://github.com/SiyuanSWang/simddna`).

The reads for the HeLa genomic DNA sample (approximately 50%) and PhiX genomic DNA that were intended to boost base diversity were excluded for analysis. The reads for the SIMD||DNA products accounted for at least 30% of total reads. For the same register type, reads were expected to have nearly identical sequences except for the register barcode (identifier for input value), the NGS barcodes (sample-specific identifier) and the regions that indicate the initial value and the locations designed with mismatches ("variable nucleotide position").

Initial filters were applied to the raw reads to filter out reads with too many sequencing errors or reads whose sample-specific barcodes could not be identified. First, 6 quality regions (sequence regions between variable nucleotide positions and/or barcodes ranging from around 15 to 75 nts in which no sequence changes are expected) were selected for the expected SIMD products of each tested register. Since sequencing quality varies by read cycle and reads may have one long high-quality region flanked by lower quality regions, we applied the criteria that reads with at least 3 *consecutive* passing regions (regions with no more than 1 mutation) were considered for analysis. If one read in a paired set of reads satisfied the above criteria, its partner read would also be included regardless of whether it independently passed the filter. This filtering process also identifies the register type of the read for sequencing runs in which PCR products from multiple registers were combined. Second, since one NGS run could contain samples from multiple samples, the sample-specific and register-specific barcodes were identified in read pairs that passed the initial filter. Reads with no more than 2 mutations to the NGS sample barcode *and* no more than 1 mutation in the SIMD||DNA barcode were included in the final analysis. Viable reads were then organized by their NGS sample barcodes and SIMD||DNA barcodes (indicating the initial values).

To determine the results of SIMD||DNA computation, each read in a qualified read pair was locally aligned to each expected cell sequence using pairwise2 from Biopython with "match", "mismatch", "gap opening", and "gap extending" scores of 1, -0.5, -0.5, and -0.5, respectively. If the aligned nucleotide at a variable nucleotide position neither matched the original sequence nor was the intended mismatch from both the forward and reverse read, the corresponding digit would be marked as undefined for that read. It is possible that results from a pair of reads at the variable nucleotide position do not match due to sequencing-related errors. In this case, the read in the pair with the greater read quality score at the variable nucleotide position was used to call the bit.

### S6 Characterizing register locations on the M13mp18 plasmid for subsequent experiments

We narrowed down our selection of register locations on the M13mp18 plasmid by performing the binary counting program on the 9 initially chosen locations with the initial values 0010 and 0111. These initial values were chosen because incrementing them involves both changing a 0 to a 1 and changing a 1 to a 0 with carryover. Recall that registers were designed with either weak, medium or strong domains by adjusting the length of the sequence allocated to each domain. Based on the binding strength, computation was performed on these registers at different temperatures: (1) "Weak" registers (M13.1, M13.2, M13.3): 40 °C for Instruction 1, 25 °C for all other instructions; (2) "Medium" registers (M13.4, M13.5, M13.6): 30 °C for all instructions; (3) "Strong" registers (M13.7, M13.8, M13.9): 40 °C for all instructions. The idea is that increasing the temperature helps eliminate spurious interactions (which involve imperfect complementarity). Since desired binding in registers with stronger domains is less susceptible to temperature, we could increase the temperature more for these registers.

In the following experiments, since homogeneous pools of registers were used (each test tube containing the same input), we relied on Sanger sequencing as a low-cost method of readout with a faster turnaround time compared to NGS. As shown in Figure S3, when temperatures are smaller than or equal to 40 °C, M13.3 and M13.8 registers updated correctly with both initial values. For registers that updated 0010 correctly, but not 0111 (e.g. M13.4, M13.5, M13.7, M13.9 registers), we hypothesize that the long cascade in instruction 1 did not complete, which logically corresponds to an error in the propagation of the carry bit. For "weak" registers that did not update 0010 correctly (e.g. M13.1, M13.2 registers), we hypothesize that some intermediate strand configurations were not stable enough (see Section S17 for more details).

In order to investigate the ability to compute on data in multiple register locations simultaneously, we need to find the reaction condition suitable for multiple locations on the same M13. Observe from Figure S3 that all of the "strong" registers updated 0010 correctly at 40 °C, but not 0111. Since we hypothesized that incorrect output was caused by Instruction 1 not completing, we increased the reaction temperature to 44 °C and 48 °C hoping to speed up toehold dissociation, which must repeatedly occur in Instruction 1. Increasing the temperature from 40 °C to 44 °C and 48 °C, the signal of the leftmost bit on input 0111 for M13.7 and M13.9 registers increased, suggesting greater propagation of the carry signal consistent with our hypothesis. Nonetheless, M13.9 registers didn't show correct computation even at 48 °C, suggesting that an even higher temperature may be needed. On the other hand, the rightmost bit of M13.8 registers failed to update on input 0010 at higher temperatures, which we attributed to the instability of some intermediate strand configurations. Thus to balance the stability and completion of the strand displacement cascade, we subsequently proceeded with M13.7 and M13.9 registers at 50 °C.

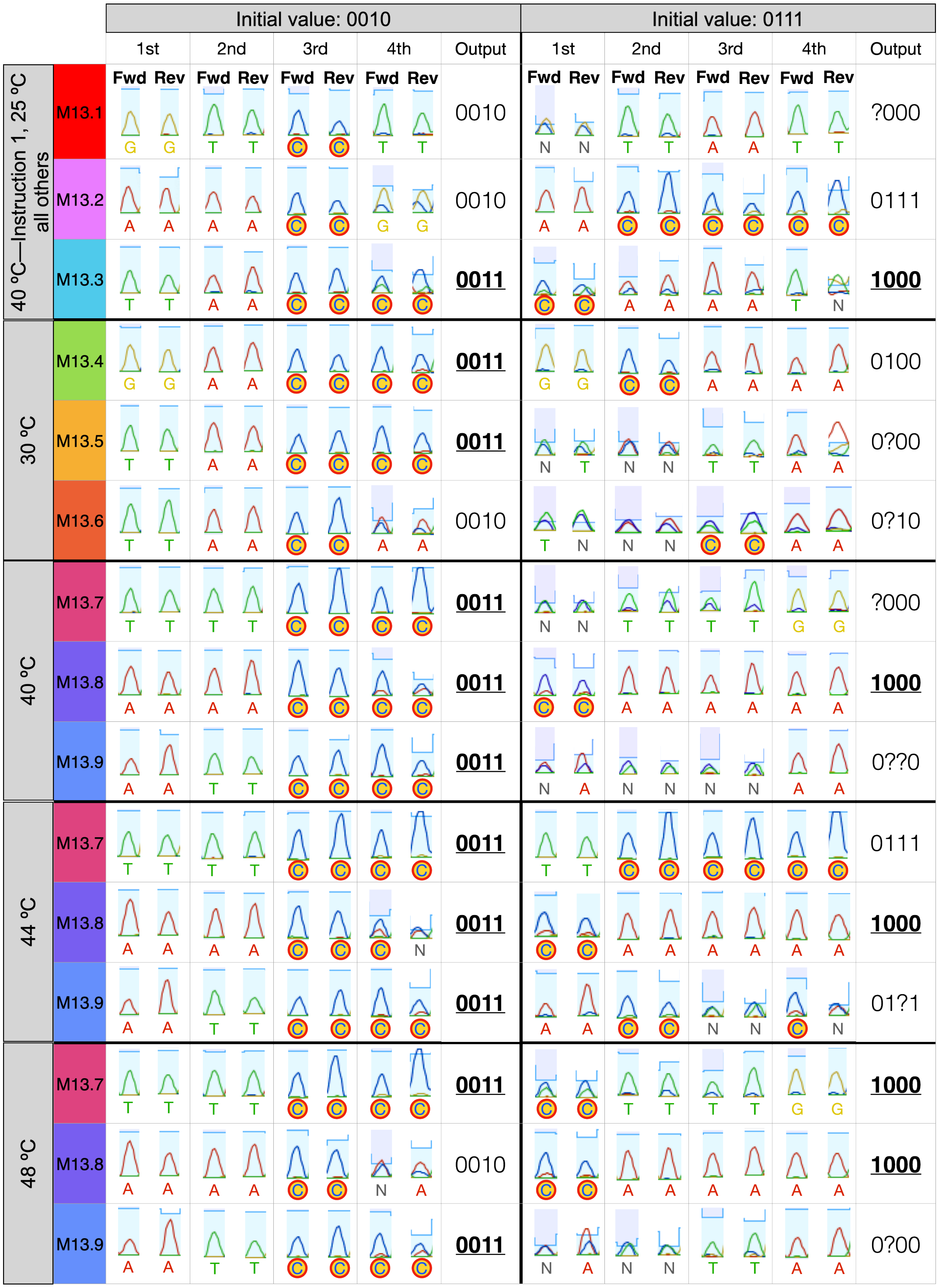

**Figure S3:** Sanger sequencing results of single data binary counting computation at 9 addresses on M13 with the initial values 0010 and 0111. Signals of mismatched nucleotides representing bit 1 are indicated by red circles. "Fwd" and "Rev" indicate forward and reverse reads respectively. "?" represents undetermined since two bases show peaks of similar height which indicates equal abundance. Correct outputs are shown in **bold** and <u>underlined</u>.

### S7 Binary counting: single instruction single data results for M13.8 and M13.3 registers

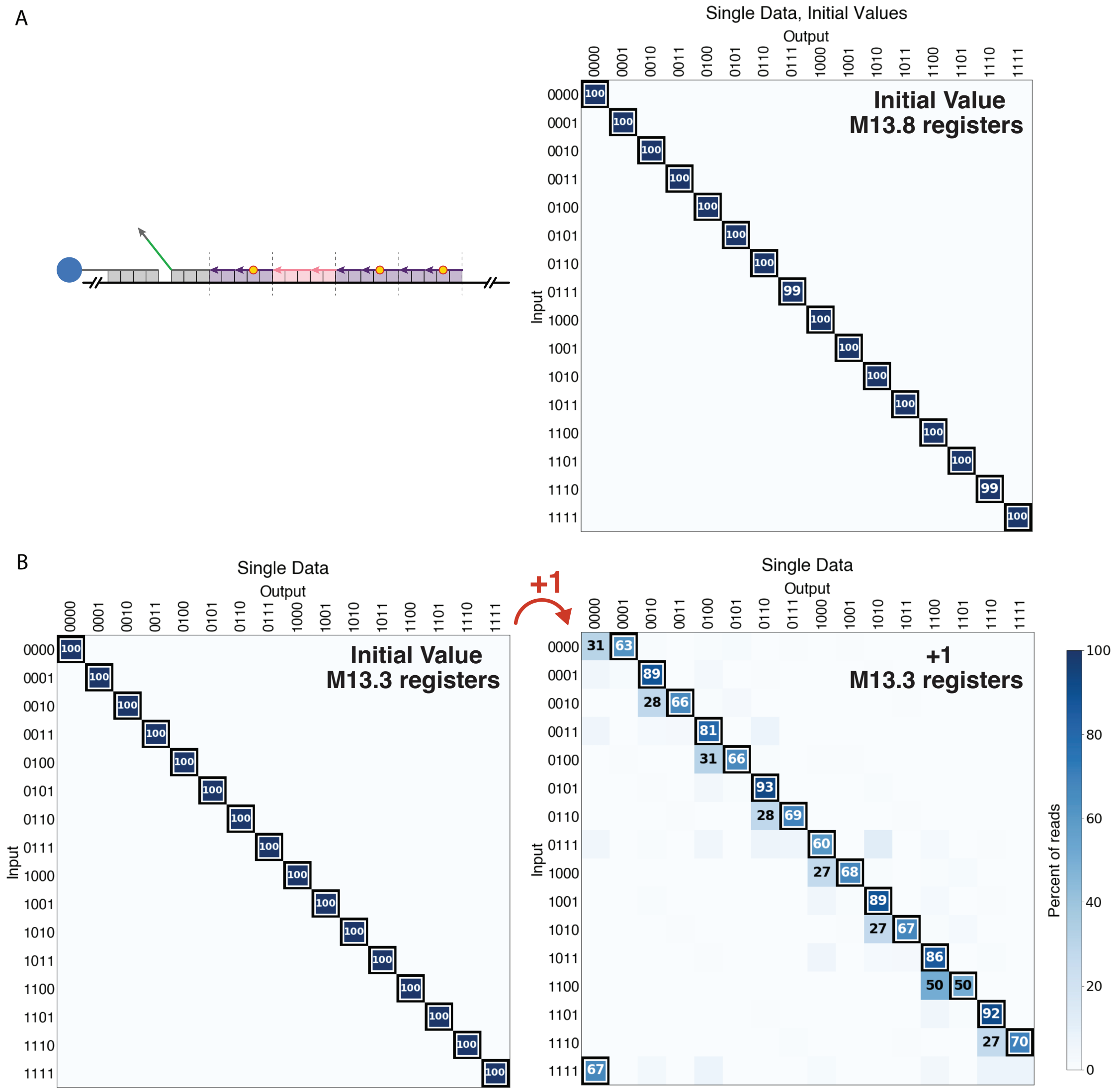

**Figure S4:** Single data binary counting on M13.8 and M13.3 registers. (A) NGS results of the initial values on M13.8 registers (computation shown in Figure 2C). (B) NGS results of the binary counting program on M13.3 registers prior to (left) and after (right) computation. Computation was performed at 40 °C on M13.8 registers. Computation was performed at 40 °C for Instruction 1, and 25 °C for other instructions on M13.3 registers.

#### S8 Binary counting: single instruction multiple data results for M13.7, M13.8 and M13.9 registers

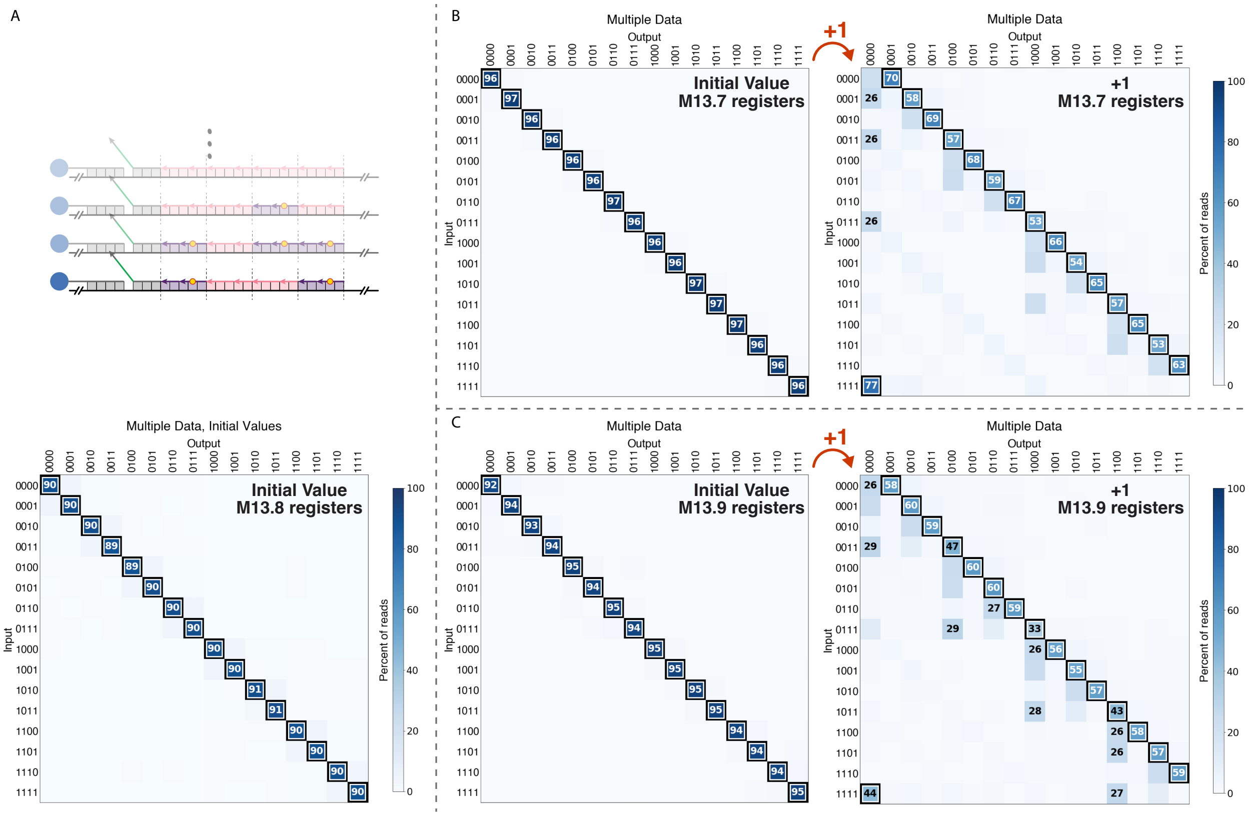

**Figure S5:** NGS results for multiple data binary counting on registers in locations (A) M13.8 (computation shown in Figure 2D) and (B) M13.7 and (C) M13.9 prior to (left) and after (right) computation. Computation was performed at 40 °C for M13.8. Computation was performed at 50 °C for M13.7 and M13.9.

#### S9 Binary counting: single instruction multiple data initial values for M13.7 and M13.9 registers on the same M13 molecules

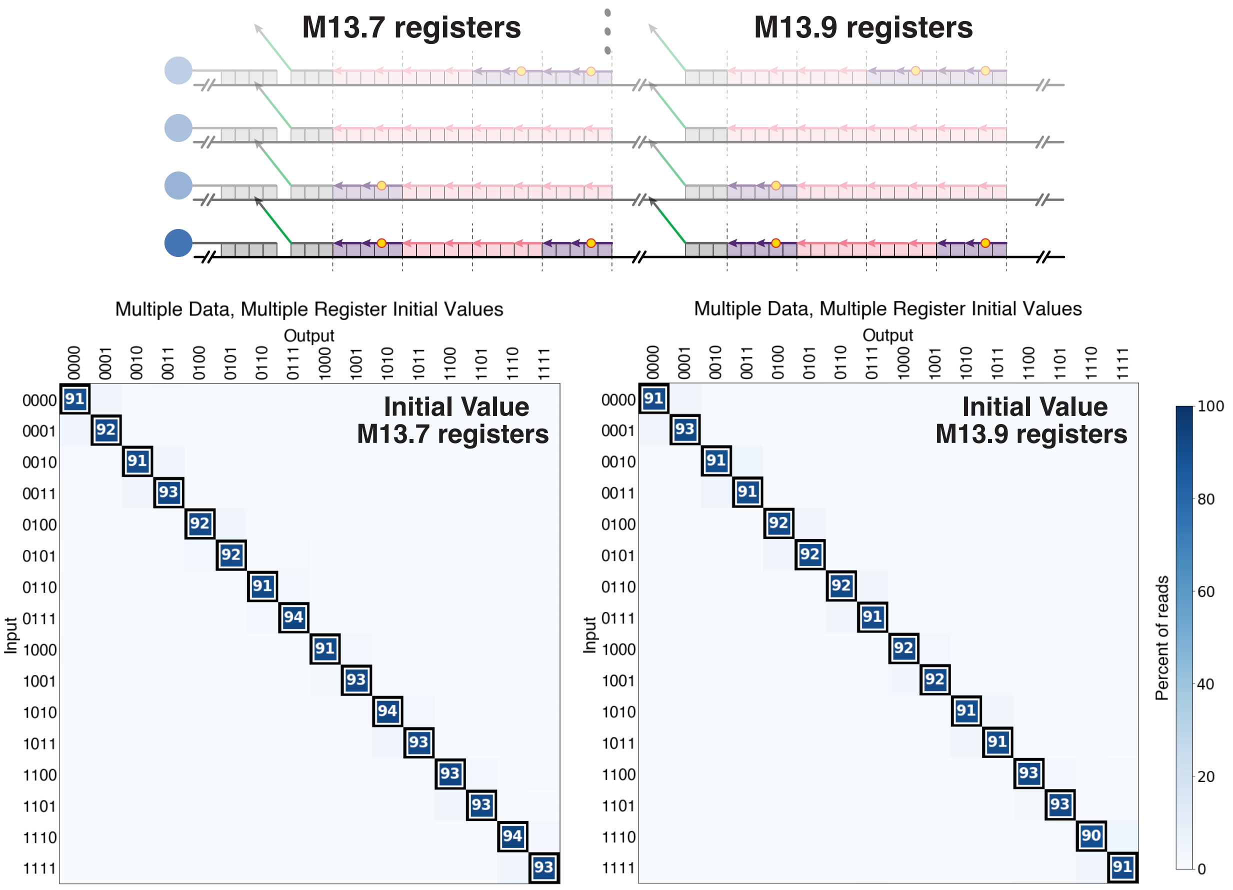

**Figure S6:** NGS results of the initial values of the multiple data binary counting on M13.7 and M13.9 registers assembled on the same M13. The results for computation are shown in Figure 2E.

#### S10 Rule 110 boundary implementation

Cellular automaton Rule 110 updates a cell according to the state of the cell itself and its neighboring cells. Therefore, cells at the two ends of a register need to perform a modified update rule which proceeds as if the missing neighbor had a defined value. We chose this boundary value to be always 0 (Figure S7, left). In principle the boundary condition could be different for each instruction by selective addition of instruction strands: To simulate the boundary condition that the bit to left of the leftmost bit is 1, Instruction 2 and 3 each has an additional strand (Figure S7, right) compared to boundary bits being 0. To simulate the boundary condition where the bit to right of the rightmost bit is 1, Instruction 1, 2, 3 and 5 each has an additional strand (Figure S7, right) compared to boundary bits being 0.

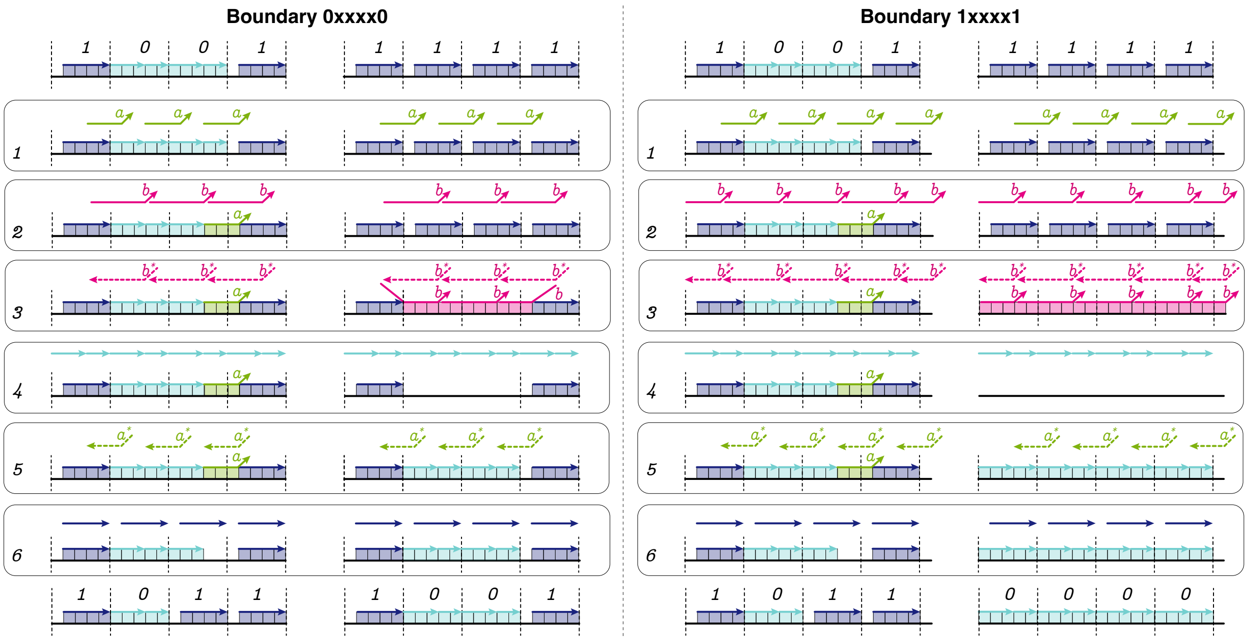

**Figure S7:** Rule 110 programs simulating constant boundary conditions 0 (left) and 1 (right). We experimentally implemented the the 0 boundary condition.

### S11 Rule 110: single instruction multiple data

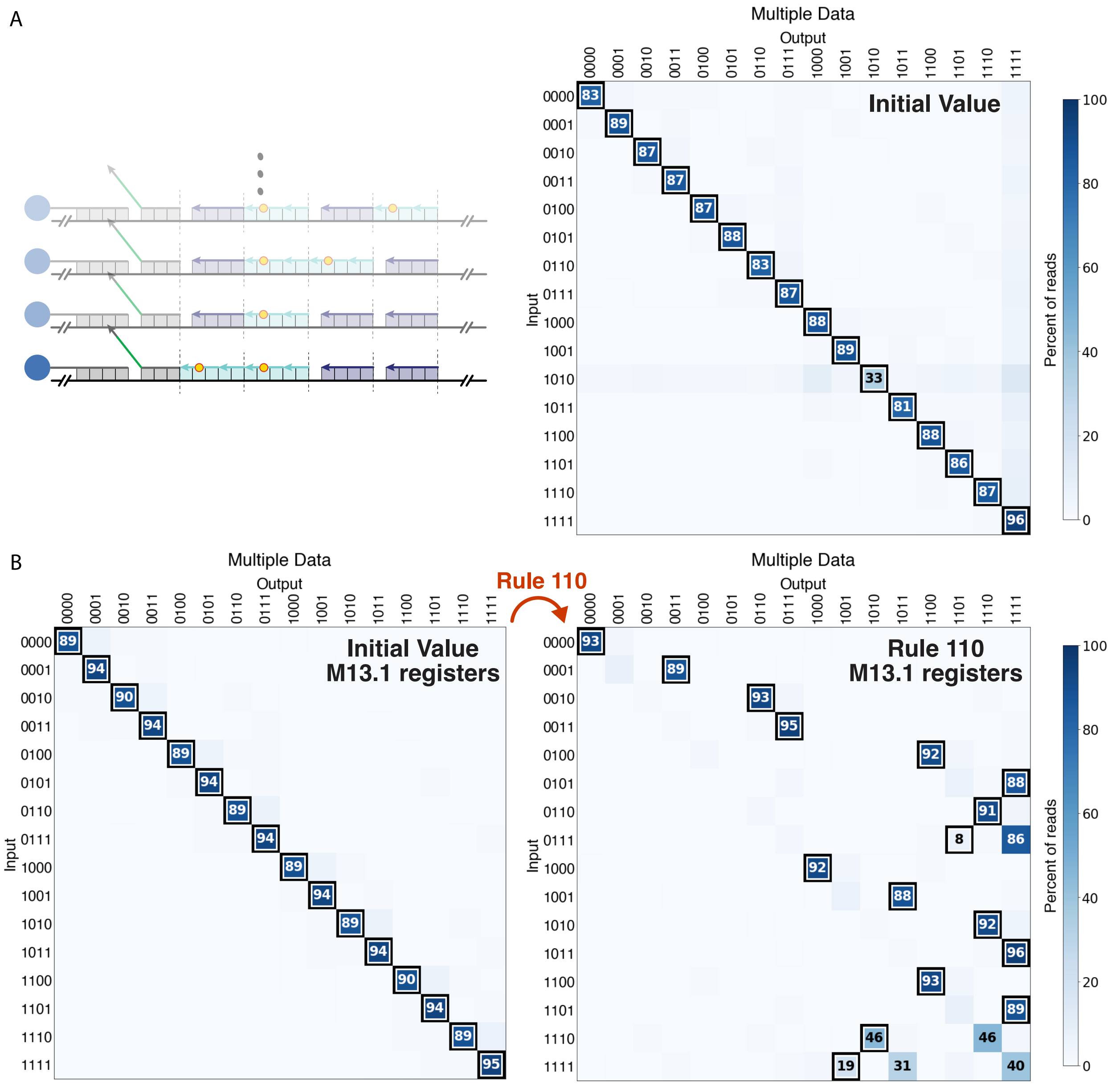

**Figure S8:** NGS results of the multiple data Rule 110 computation on (A) artificially designed registers (computation shown in Figure 3) and (B) M13.1 registers prior to (left) and after (right) computation. Computation was performed at 25 °C.

#### S12 Random-access specificity and selective erasure

##### S12.1 Random access

To evaluate random-access specificity, we analyzed both whether the target registers were queried successfully from the register mix and whether the queried registers had the correct output values. For the Rule 110 computation and queries shown in (Figure 4A), we confirmed that the registers queried from the register mix had the expected barcodes (Figure S9). Figure 4A shows the distribution of output values on these registers.

To further investigate the specificity of random access and the fidelity of binary counting computation, we performed a separate experiment shown in Figure S10. We mixed registers with all 16 input values, performed the binary counting computation, and finally queried barcodes corresponding to 14 inputs in parallel, confirming that all of them can be accessed with high specificity.

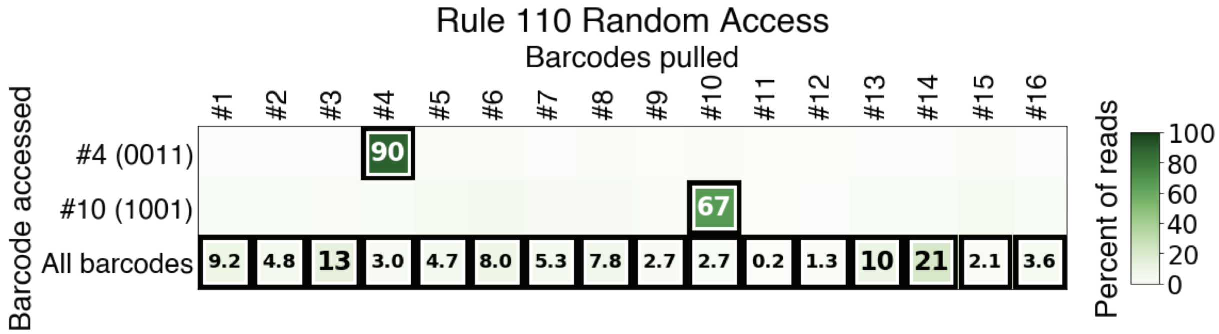

**Figure S9:** NGS results of the percentage of the read out registers with the corresponding barcodes for the sequential random-access experiments after the Rule 110 program. The registers sequentially pulled out are shown in Figure 4. The query strand concentration is lower than the estimated register concentration, and thus displacement achieves partial extraction of registers.

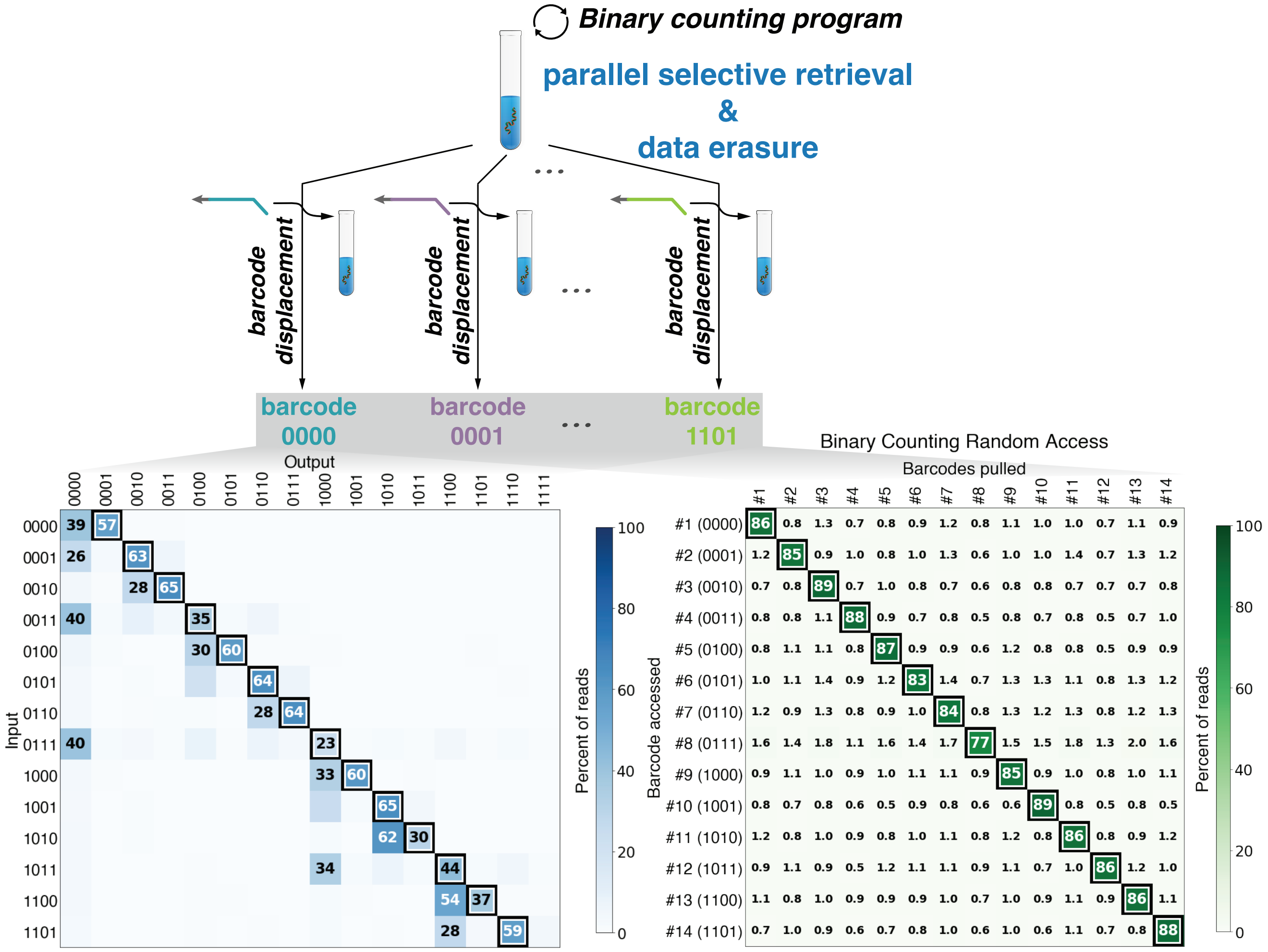

**Figure S10:** NGS results of parallel random-access experiments for the binary counting program. To distinguish between the register values after measurement and register-specific barcodes (each is associated with a unique initial value), we use blue to indicate the measured bit values encoded on reads and green to indicate barcodes associated with reads. Computation was performed at 40 °C for Instruction 1, and 25 °C for other instructions.

#### S12.2 Selective erasure

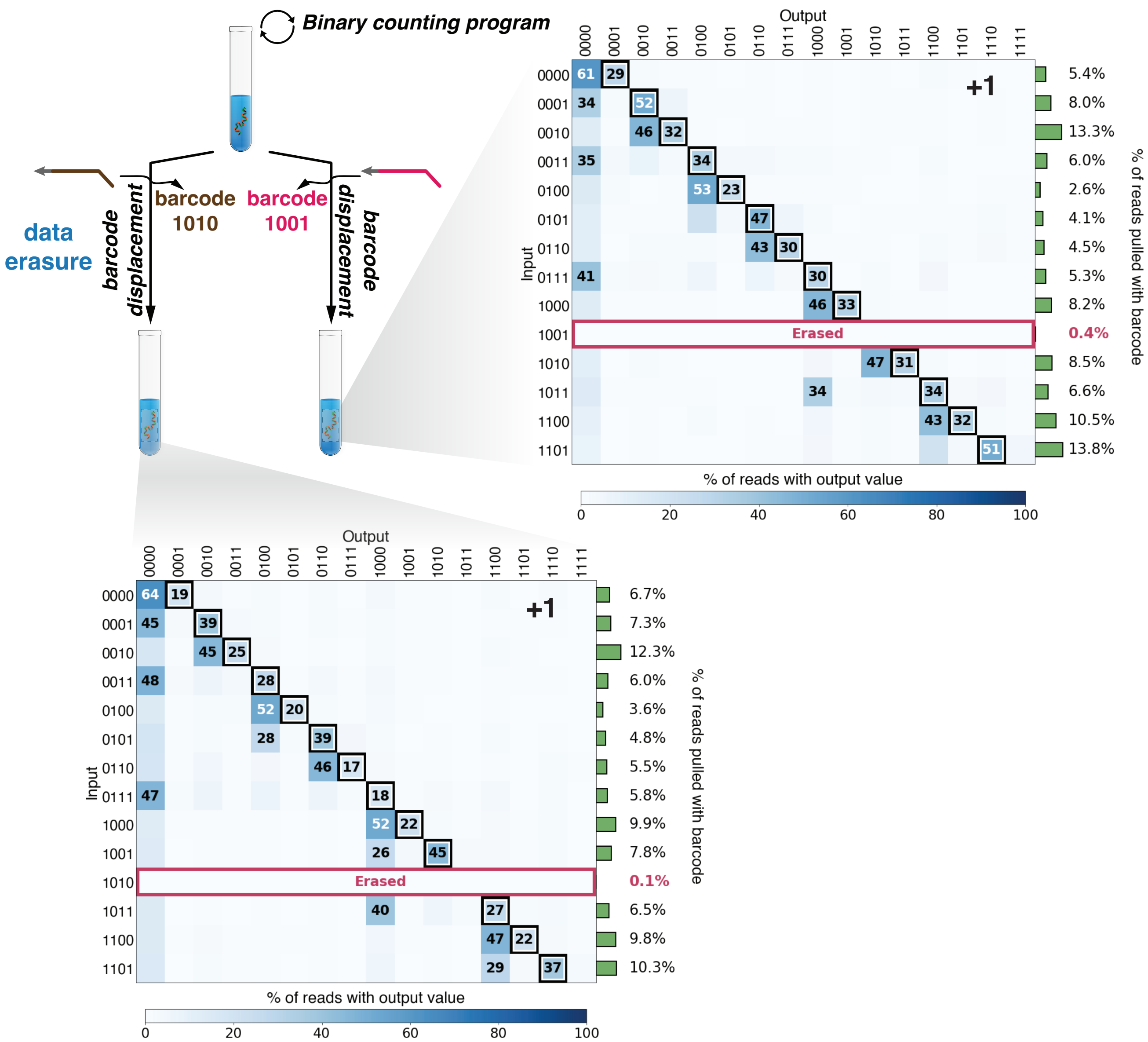

**Figure S11:** NGS results of the remaining registers after selective erasure. The erased registers are different from Figure 4B (query strand 1000 vs 1010). The query strand concentration is higher than the estimated register concentration, and thus displacement achieves full erasure of registers. The registers are composed of chemically synthesized DNA. The binary counting computation was performed at 40 °C for Instruction 1, and 25 °C for other instructions.

### S13 Multiple rounds of binary counting program

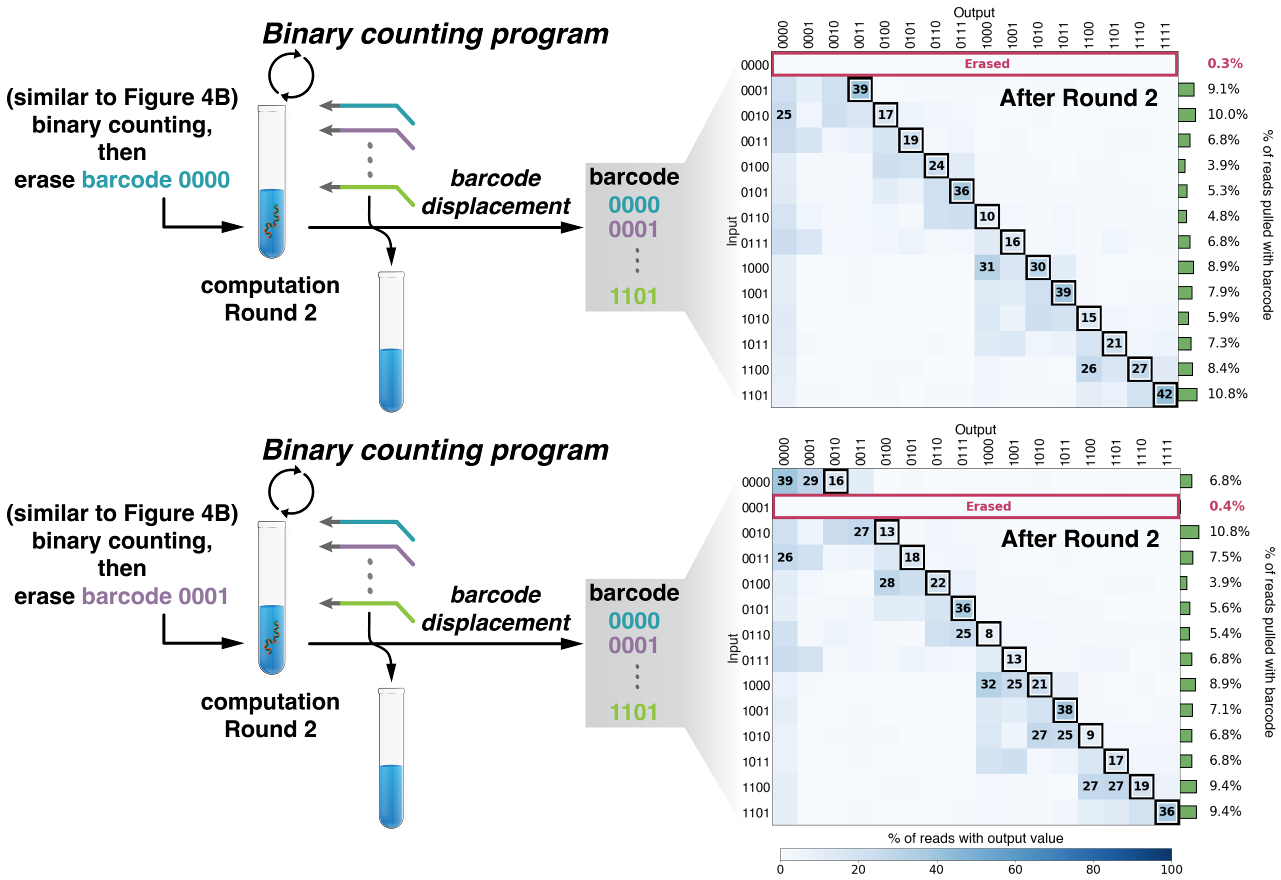

**Figure S12:** NGS results after two rounds of binary counting. The mix of inputs went through one round of computation, and then selective erasure, and finally another round of binary counting computation (40 °C for Instruction 1, and 25 °C for other instructions). The registers are composed of chemically synthesized DNA.

#### S14 Washing efficiency

To determine the number of washing steps needed, we incubated registers with a non-interacting fluorophore labeled strands and then measured the fluorescence signals after multiple steps of washing. As shown in Figure S13, 2 steps of washing was decided for all the experiments.

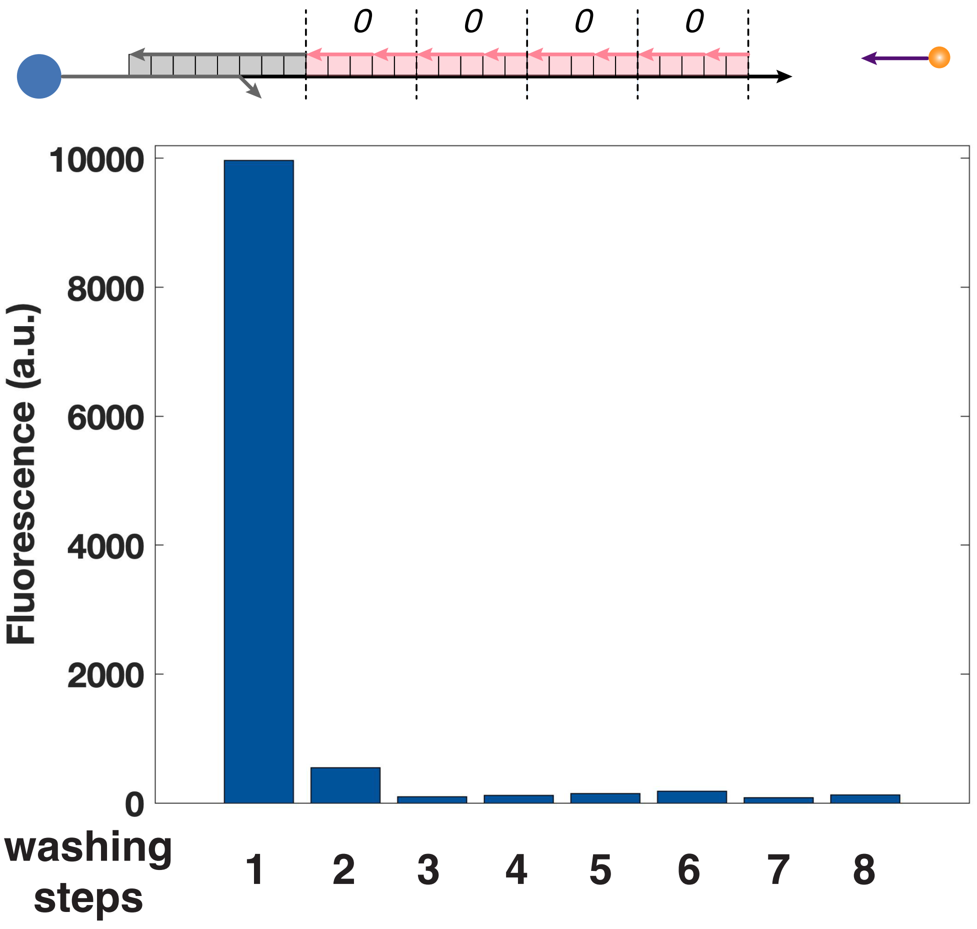

**Figure S13:** Fluorescence signal after multiple washing steps. The concentration of the fluorophore labeled strand was the same as the highest concentration of the instruction strands. The background fluorescence signal from magnetic beads was set to be the baseline and subtracted. After washing twice, fluorescence signal significantly dropped close to background.

#### S15 Investigating register loss during SIMD||DNA computation

There are three factors contributing to the loss of SIMD||DNA products that is not captured by measuring the read fraction of correct output: (1) bead loss during washing, (2) ligation failure due to incorrect computation (desired strands labeled with phosphate are not attached), incomplete ligation, or (3) incomplete displacement of the registers from the bead. Since registers can be fully erased as shown in Figure 4B, S11 and S12, (3) may not contribute significantly to product loss.

To break down the sources of product loss from (1) and (2), we compared the amount of products after the process of one round of computation and the process of washing with the same number of washing steps as computation.

The amount of products were quantified using a combination of qPCR and quantitative electrophoretic techniques. In qPCR, the signal strength depends on the sample concentration and it doubles at each cycle. $C_q$ is defined as the cycle number at which fluorescence signal from the sample diverges from background. We used $C_{product}$ and $C_{initial}$ to represent $C_q$ values for product samples and control samples with no computation or washing treatment respectively. Yield was calculated as $2^{C_{initial}-C_{product}}$. As shown in Figure S14A, the majority of product loss is contributed by bead loss during washing steps: Washing accounts for 76% of total product loss

$$\frac{\texttt{Amount lost after washing}}{\texttt{Amount lost after washing + computation}} = (1-0.53)/(1-0.381) = 76\%$$

The qPCR results were corroborated with gel quantification (Agilent 2100 BioAnalyzer). We observed a similar yield with multi-round Rule 110 computation, with an average of about 28% per round (i.e. $(0.063/0.789)^{1/n}$ = 0.283, for $n$ = 2 rounds of computation).

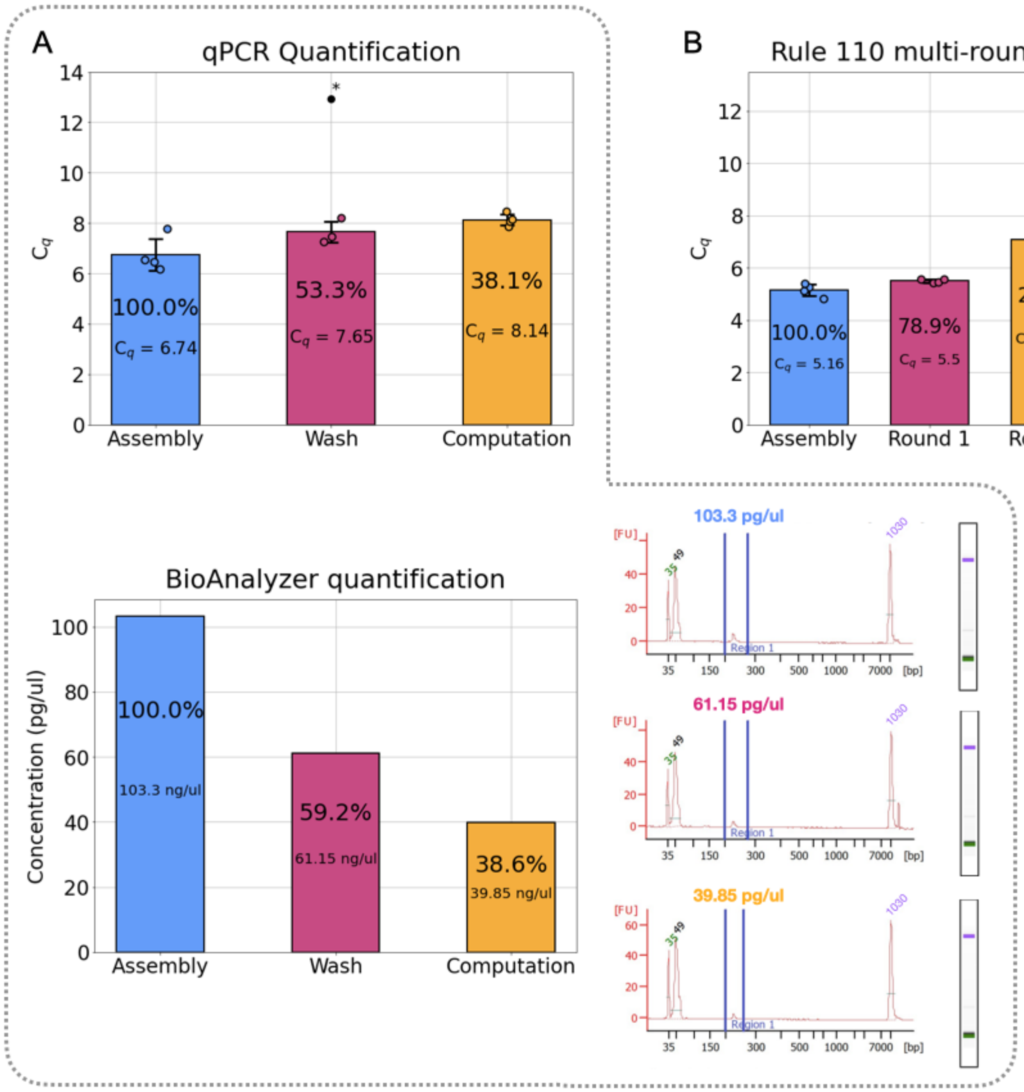

**Figure S14:** Quantifying the yield of SIMD||DNA computation. (A) The amount of SIMD||DNA products were quantified by both qPCR (top) and electrophoresis (bottom). The samples were treated in the following ways: registers were annealed only ("Assembly"), one round of Rule 110 computation was performed ("Computation"), or registers were washed as many times as in the computation condition, but without adding instruction strands ("Wash"). The calculated percent yield is shown in text, as well as the $C_q$ and concentration as determined by qPCR and the Bio-Analyzer, respectively. Asterisk (*) denotes an outlier. (B) Yield quantified by qPCR for multiple rounds of Rule 110 computation. "Assembly" refers to registers that were annealed only. In combination with the results from (A), each round of computation resulted in about 25% to 40% remaining product (e.g. 38.1% product remaining in part A Round 1, 33.3% in part B Round 1 to Round 2, 24.0% in part B Round 2 to Round 3). Note that there is a discrepancy between (A) and (B) in the yield from the initial amount (assembled registers) to the first round of computation. This could be caused by the differences in reaction conditions and washing procedures: Experiments in (A) were performed on registers with 16 inputs (total 50 nM) accessed by 20 nM of query strands, while experiments in (B) were performed on registers with 5 inputs accessed by 30 nM of query strands. Data points represent the $C_q$'s of each SIMD||DNA product aliquots (collected from the same sample) as determined by qPCR.

#### S16 Investigation of instruction completeness through fluorescence

Fluorescence experiments guided our initial choice of reaction conditions. Higher temperature and reaction time increased the completion of instruction 1. Since reaction time affected completion by less than 10%, we still chose the reaction time to be 10 min to reduce the total time cost. Instructions 2, 3, 5 and 6 completed at 10 mins and at their corresponding instruction strand concentrations. Instructions 4 and 7 are not shown because we were not able to successfully prepare the expected register state prior to the instruction. (We hypothesize that annealing these register states directly was hindered by spurious interactions on long single regions on the bottom strand. Note that when annealing the initial register state for the other instructions, most of the register bottom strand was double-stranded.)

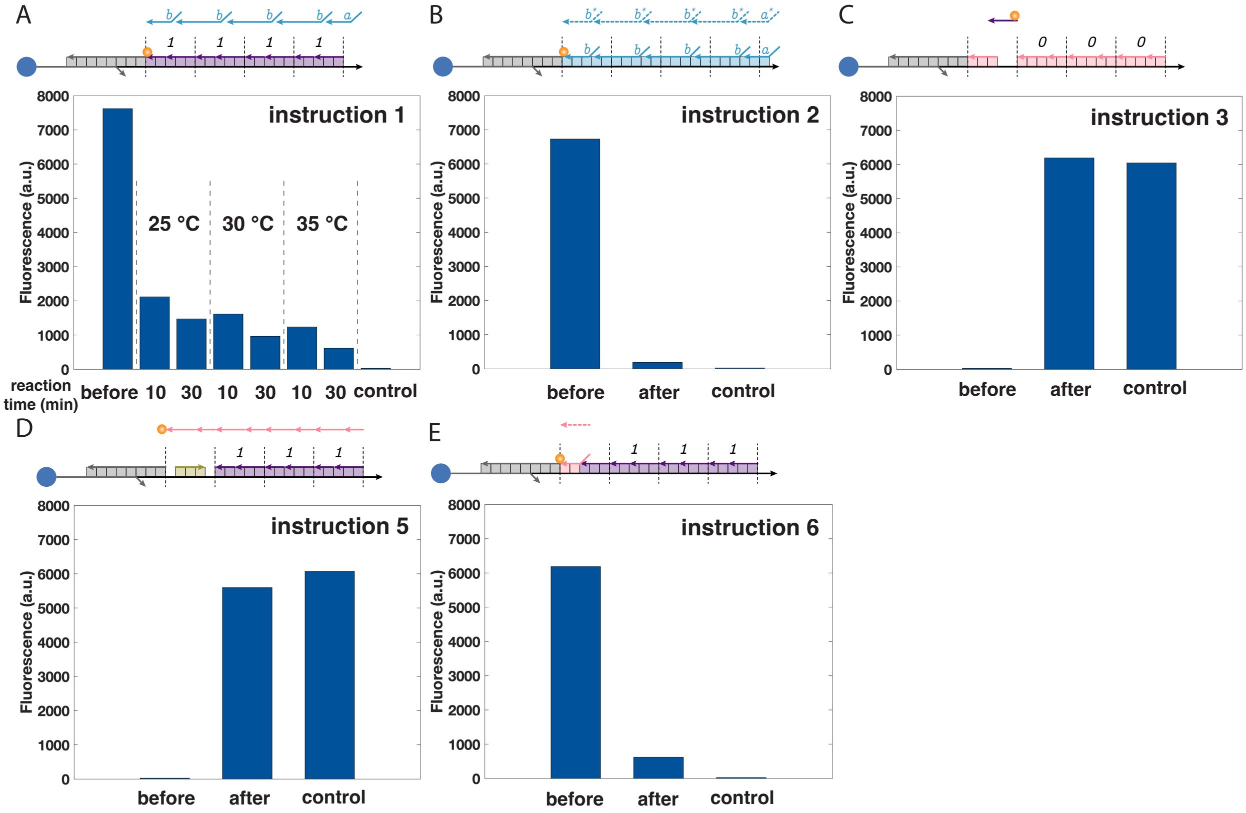

**Figure S15:** Fluorescence results on individual instructions for the binary counting program. The concentrations were 3 $\mu$M for (A) Instruction 1, 0.5 $\mu$M for Instruction (B) 2, (C) 3 and (E) 6, 1 $\mu$M for (D) Instruction 5. Other than Instruction 1, all other instructions were performed at 25 °C. The register shown on the top of the figure was initially prepared and measured. The registers assembled as would be expected after the instruction were measured as the control signals. Fluorescence signal was measured after reaction incubation and washing. The background fluorescence signal from magnetic beads was set to be the baseline and subtracted.

#### S17 Hypotheses about error

From results for both the binary counting and Rule 110 programs presented in this paper, we observed errors in computation. In generally, the current design is not robust to toeless displacement (leak)—undesired displacement reactions that initiate from fraying at nicks—which may account for some of the errors [59]. Since leak increases with thermodynamic bias and the duration of the reaction, we limited fuel concentrations and reaction time——which, in turn, may have led to lower completion levels. Indeed, in troubleshooting we observed that some instructions did not reach full completion (Section S16). The recently published "leakless" design [60, 59] reduces leak by utilizing redundancy, similar to error correction strategies in communications and electrical engineering [61]. Restricting to toehold-exchange reactions alone can also reduce leak [62]. By incorporating such leak reduction strategies and increasing the thermodynamic bias through larger concentrations and longer toeholds, may increase completion without increasing leak. In addition to preventing errors at the strand displacement level, it may be possible to address errors at the "software level" by employing error correction codes in the data and error correction schemes in the instructions.

Below we present a few specific hypotheses about the cause of error and their role in the computation.

For the binary counting program, we notice that some 0's fail to change to 1's. This could be due to a lack of "protection" on the cell that is supposed to change to 1 after Instruction 3, so that strands at Instruction 5 can bind to the cell and "lock" the bit to 0. There are two possible causes for the failed protection: One error pathway is that the intermediate configuration after Instruction 3 is not stable (Figure S16). Since the domain binding stability was not designed with consideration of mismatches, the purple strand with a mismatch may be more likely to dissociate. Another error pathway can be caused by spurious binding of Instruction 3 strands to locations other than the desired protection location. The spurious binding may occur because there exists a long unbound region on the register bottom strand after Instruction 2, and Instruction 3 may spuriously bind to the unbound region.

For the cells that fail to change to 0, the strand displacement cascade in Instruction 1 may not reach completion (Figure S15).

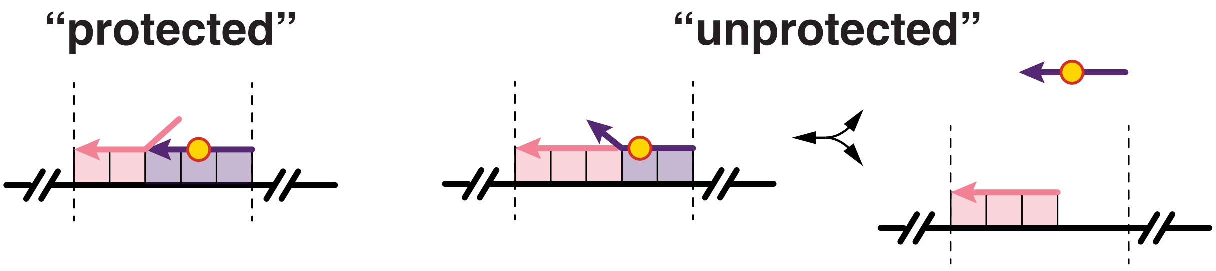

**Figure S16:** Example of one possible pathway for failed protection on a cell that is supposed to change to 1.

For the Rule 110 program, we notice that some 0 fails to change to 1. This could be due to that the strands from Instruction 3 spuriously bind with the domain $a$ and displace the Instruction 1 strand by mistake, and thus the cell changes to 0 in Instruction 4 and stays at 0. For the cells that fail to change to 0, it is likely due to incompletion of the cooperative displacement in Instruction 2.

#### S18 The number of strand displacement steps performed on naturally-occurring sequences

| Binary counting initial values | Instructions | | | | | | | Sum |
|---|---|---|---|---|---|---|---|---|
| | 1 | 2 | 3 | 4 | 5 | 6 | 7 | |
| 0000 | 1 | 1 | 1 | | | 1 | | 4 |
| 0001 | 2 | 2 | 1 | 1 | 1 | 1 | | 8 |
| 0010 | 1 | 1 | 1 | | | 1 | | 4 |
| 0011 | 3 | 3 | 1 | 2 | 2 | 1 | | 12 |
| 0100 | 1 | 1 | 1 | | | 1 | | 4 |
| 0101 | 2 | 2 | 1 | 1 | 1 | 1 | | 8 |
| 0110 | 1 | 1 | 1 | | | 1 | | 4 |
| 0111 | 4 | 4 | 1 | 3 | 3 | 1 | | 16 |
| 1000 | 1 | 1 | 1 | | | 1 | | 4 |
| 1001 | 2 | 2 | 1 | 1 | 1 | 1 | | 8 |
| 1010 | 1 | 1 | 1 | | | 1 | | 4 |
| 1011 | 3 | 3 | 1 | 2 | 2 | 1 | | 12 |
| 1100 | 1 | 1 | 1 | | | 1 | | 4 |
| 1101 | 2 | 2 | 1 | 1 | 1 | 1 | | 8 |
| 1110 | 1 | 1 | 1 | | | 1 | | 4 |
| 1111 | 5 | 5 | | 4 | 4 | | | 18 |
| Total | | | | | | | | 122 |

**Table S3:** Number of strand displacement steps of the binary counting program performed on naturally-occurring sequences.

#### S19 Algorithmic query

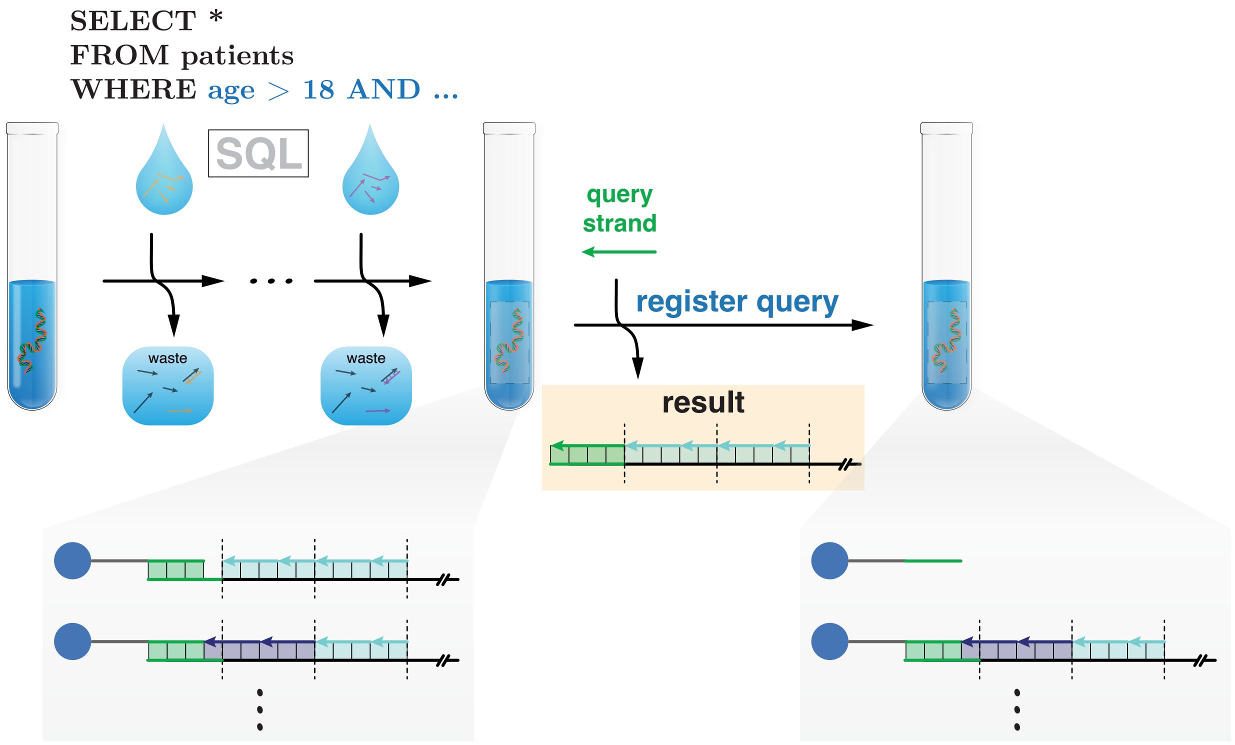

**Figure S17:** More advanced querying similar to the Structured Query Language (SQL) could be used to select data satisfying certain criteria from a database. Such algorithmic queries could be practically realized via SIMD||DNA programs that control the accessibility of the toehold for the query strand as the output of the computation.

### S20 Sequences

##### Artificially designed adaptor and displacement sequences

| | |
|---|---|
| bottom | ACAACTC TCACTTC CAACTCA CACATAC CTTTACA TCTCCT ACACACA CATTTAC TCAAACA TCTATCA ACAAAC CTTTCTC CTTCAAC CTAACTC TCTCTAC TCCACT ACATAAC CAAACAC CAATCTC ACAATCA TCATTC CATTTCA CAAAACT TCAAATC CTTAATC |
| biotin1 | /5Biosg/AAAAAAAAAAAAAAAAAAAA CATCCATTCCACTCA CAAAACAAAACCTCA TTACCCT |
| biotin2 | /5Biosg/AAAAAAAAAAAAAAAAAAAA CAACATATCAATTCA CACTAACATACAACA TACACTC |
| biotin3 | /5Biosg/AAAAAAAAAAAAAAAAAAAA CAATATCCATAACCA CATCAATCAACACCA ATCCAAC |
| biotin4 | /5Biosg/AAAAAAAAAAAAAAAAAAAA CACAACTCATTACCA CATTATTCAAACCCA CCATTAC |
| biotin5 | /5Biosg/AAAAAAAAAAAAAAAAAAAA CACACTATAATTCCA CACTTCATAAATCCA CTACTTC |
| biotin6 | /5Biosg/AAAAAAAAAAAAAAAAAAAA CAACCATACTAAACA CATTCCTACATTTCA CTCAATC |
| biotin7 | /5Biosg/AAAAAAAAAAAAAAAAAAAA CAAATCTTCATCCCA CACTCATCCTTTACA TCTTTCC |
| biotin8 | /5Biosg/AAAAAAAAAAAAAAAAAAAA CAACTCAAACATACA CACATAACAAAACCA CCTAAAC |
| biotin9 | /5Biosg/AAAAAAAAAAAAAAAAAAAA CACTATACACACCCA CAATACAAATCCACA CCTCCTT |
| biotin10 | /5Biosg/AAAAAAAAAAAAAAAAAAAA CAACAAACCATTACA CACTTTTCACTATCA CCTATCA |
| biotin11 | /5Biosg/AAAAAAAAAAAAAAAAAAAA CACCCAAAACCCACA CAAACCCAACTCACA ACATACC |
| biotin12 | /5Biosg/AAAAAAAAAAAAAAAAAAAA CACATAACAACCACA CACCCTAAAATCTCA ACTCTAC |
| biotin13 | /5Biosg/AAAAAAAAAAAAAAAAAAAA CATTACCAACCACCA CATCCTTAACTCCCA CAAATCC |
| biotin14 | /5Biosg/AAAAAAAAAAAAAAAAAAAA CACTTCACAACTACA CACAAACTACATCCA AAACCAC |
| biotin15 | /5Biosg/AAAAAAAAAAAAAAAAAAAA CATCATACCTACTCA CAACTCCTAATATCA TTCATCC |
| biotin16 | /5Biosg/AAAAAAAAAAAAAAAAAAAA CAATTCACTCAATCA CACCACCAAACTTCA CATTCTC |
| leftadaptor1 | /5Phos/TGAGTTGGAAGTGAGAGTTGT TGAGGTTTTGTTTTG TGAGTGGAATGGATG CCGAGCCCACGAGAC |
| leftadaptor2 | /5Phos/TGAGTTGGAAGTGAGAGTTGT TGTTGTATGTTAGTG TGAATTGATATGTTG CCGAGCCCACGAGAC |
| leftadaptor3 | /5Phos/TGAGTTGGAAGTGAGAGTTGT TGGTGTTGATTGATG TGGTTATGGATATTG CCGAGCCCACGAGAC |
| leftadaptor4 | /5Phos/TGAGTTGGAAGTGAGAGTTGT TGGGTTTGAATAATG TGGTAATGAGTTGTG CCGAGCCCACGAGAC |
| leftadaptor5 | /5Phos/TGAGTTGGAAGTGAGAGTTGT TGGATTTATGAAGTG TGGAATTATAGTGTG CCGAGCCCACGAGAC |
| leftadaptor6 | /5Phos/TGAGTTGGAAGTGAGAGTTGT TGAAATGTAGGAATG TGTTTAGTATGGTTG CCGAGCCCACGAGAC |
| leftadaptor7 | /5Phos/TGAGTTGGAAGTGAGAGTTGT TGTAAAGGATGAGTG TGGGATGAAGATTTG CCGAGCCCACGAGAC |
| leftadaptor8 | /5Phos/TGAGTTGGAAGTGAGAGTTGT TGGTTTTGTTATGTG TGTATGTTTGAGTTG CCGAGCCCACGAGAC |
| leftadaptor9 | /5Phos/TGAGTTGGAAGTGAGAGTTGT TGTGGATTTGTATTG TGGGTGTGTATAGTG CCGAGCCCACGAGAC |
| leftadaptor10 | /5Phos/TGAGTTGGAAGTGAGAGTTGT TGATAGTGAAAAGTG TGTAATGGTTTGTTG CCGAGCCCACGAGAC |
| leftadaptor11 | /5Phos/TGAGTTGGAAGTGAGAGTTGT TGTGAGTTGGGTTTG TGTGGGTTTTGGGTG CCGAGCCCACGAGAC |
| leftadaptor12 | /5Phos/TGAGTTGGAAGTGAGAGTTGT TGAGATTTTAGGGTG TGTGGTTGTTATGTG CCGAGCCCACGAGAC |
| leftadaptor13 | /5Phos/TGAGTTGGAAGTGAGAGTTGT TGGGAGTTAAGGATG TGGTGGTTGGTAATG CCGAGCCCACGAGAC |
| leftadaptor14 | /5Phos/TGAGTTGGAAGTGAGAGTTGT TGGATGTAGTTTGTG TGTAGTTGTGAAGTG CCGAGCCCACGAGAC |
| leftadaptor15 | /5Phos/TGAGTTGGAAGTGAGAGTTGT TGATATTAGGAGTTG TGAGTAGGTATGATG CCGAGCCCACGAGAC |
| leftadaptor16 | /5Phos/TGAGTTGGAAGTGAGAGTTGT TGAAGTTTGGTGGTG TGATTGAGTGAATTG CCGAGCCCACGAGAC |
| rightAdaptor | TCGTCGGCAGCGTC GATTAAG GATTTGA |

| | |
|---|---|
| displacement1 | AGGGTAA TGAGGTTTTGTTTTG TGAGTGGAATGGATG |
| displacement2 | GAGTGTA TGTTGTATGTTAGTG TGAATTGATATGTTG |
| displacement3 | GTTGGAT TGGTGTTGATTGATG TGGTTATGGATATTG |
| displacement4 | GTAATGG TGGGTTTGAATAATG TGGTAATGAGTTGTG |
| displacement5 | GAAGTAG TGGATTTATGAAGTG TGGAATTATAGTGTG |
| displacement6 | GATTGAG TGAAATGTAGGAATG TGTTTAGTATGGTTG |
| displacement7 | GGAAAGA TGTAAAGGATGAGTG TGGGATGAAGATTTG |
| displacement8 | GTTTAGG TGGTTTTGTTATGTG TGTATGTTTGAGTTG |
| displacement9 | AAGGAGG TGTGGATTTGTATTG TGGGTGTGTATAGTG |
| displacement10 | TGATAGG TGATAGTGAAAAGTG TGTAATGGTTTGTTG |
| displacement11 | GGTATGT TGTGAGTTGGGTTTG TGTGGGTTTTGGGTG |
| displacement12 | GTAGAGT TGAGATTTTAGGGTG TGTGGTTGTTATGTG |
| displacement13 | GGATTTG TGGGAGTTAAGGATG TGGTGGTTGGTAATG |
| displacement14 | GTGGTTT TGGATGTAGTTTGTG TGTAGTTGTGAAGTG |
| displacement15 | GGATGAA TGATATTAGGAGTTG TGAGTAGGTATGATG |
| displacement16 | GAGAATG TGAAGTTTGGTGGTG TGATTGAGTGAATTG |

**Note that the cell number and the instruction strands are named from left to right as shown in the scheme. Nucleotides marked in color are mismatches compared to the bottom strand, which comprise our secondary encoding of register value (Red for binary counting, Blue for Rule 110).**

**Artificially designed sequences for the binary counting program**

| | |
|---|---|
| cell1ZeroLeft | /5Phos/AGGAGA TGTAAAG GTATGTG |
| cell1ZeroRight | /5Phos/GTAAATG TGTGTGT |
| cell1OneLeft | /5Phos/TGTAAAG GTATGTG |
| cell1OneRight | /5Phos/GTAAAGG TGTGTGT AGGAGA |
| cell2ZeroLeft | /5Phos/GTTTGT TGATAGA TGTTTGA |
| cell2ZeroRight | /5Phos/GTTGAAG GAGAAAG |
| cell2OneLeft | /5Phos/TGATAGA TGTTTGA |
| cell2OneRight | /5Phos/GTTGGAG GAGAAAG GTTTGT |
| cell3ZeroLeft | /5Phos/AGTGGA GTAGAGA GAGTTAG |
| cell3ZeroRight | /5Phos/GTGTTTG GTTATGT |
| cell3OneLeft | /5Phos/GTAGAGA GAGTTAG |
| cell3OneRight | /5Phos/GTGTTTG GGTATGT AGTGGA |
| cell4ZeroLeft | /5Phos/GAATGA TGATTGT GAGATTG |
| cell4ZeroRight | /5Phos/AGTTTTG TGAAATG |
| cell4OneLeft | /5Phos/TGATTGT GAGATTG |
| cell4OneRight | /5Phos/AGTTTTG GGAAATG GAATGA |
| instru11 | AGGATGA AGGAGA TGTAAAG GTATGTG |
| instru12 | AGGATGA GTTTGT TGATAGA TGTTTGA GTAAATG TGTGTGT |
| instru13 | AGGATGA AGTGGA GTAGAGA GAGTTAG GTTGAAG GAGAAAG |
| instru14 | AGGATGA GAATGA TGATTGT GAGATTG GTGTTTG GTTATGT |
| instru15 | AGAAAGA GATTTGA AGTTTTG TGAAATG |
| instru21 | CACATAC CTTTACA TCTCCTT CATCCT |
| instru22 | ACACACA CATTTAC TCAAACA TCTATCA ACAAACT CATCCT |
| instru23 | CTTTCTC CTTCAAC CTAACTC TCTCTAC TCCACTT CATCCT |
| instru24 | ACATAAC CAAACAC CAATCTC ACAATCA TCATTCT CATCCT |
| instru25 | CATTTCA CAAAACT TCAAATC TCTTTCT |
| instru41 | TGTGTGT AGGAGA TGTAAAG |
| instru42 | GAGAAAG GTTTGT TGATAGA |
| instru43 | GTTATGT AGTGGA GTAGAGA |
| instru44 | TGAAATG GAATGA TGATTGT |
| instru61 | CACATAC CTTTACA TCTCCT |
| instru62 | TCAAACA TCTATCA ACAAAC |
| instru63 | CTAACTC TCTCTAC TCCACT |
| instru64 | CAATCTC ACAATCA TCATTC |

**Artificially designed sequences for the Rule 110 program**

Note that the strands for Instruction 2 and 3 in the Rule 110 program are shared with the strands for Instruction 1 and 2 in the binary counting program.

| | |
|---|---|
| cell1zeroLeft | /5Phos/AGGAGA TGGAAAG GTATGTG |
| cell1zeroRight | /5Phos/GTAAATG TGTGTGT |
| cell2zeroLeft | /5Phos/GTTTGT TGAGAGA TGTTTGA |
| cell2zeroRight | /5Phos/GTTGAAG GAGAAAG |
| cell3zeroLeft | /5Phos/AGTGGA GGAGAGA GAGTTAG |
| cell3zeroRight | /5Phos/GTGTTTG GTTATGT |
| cell4zeroLeft | /5Phos/GAATGA TGAGTGT GAGATTG |
| cell4zeroRight | /5Phos/AGTTTTG TGAAATG |
| cell1one | GTAAATG TGTGTGT AGGAGA TGTAAAG |
| cell2one | GTTGAAG GAGAAAG GTTTGT TGATAGA |
| cell3one | GTGTTTG GTTATGT AGTGGA GTAGAGA |
| cell4one | AGTTTTG TGAAATG GAATGA TGATTGT |
| seal-cell1one | /5Phos/GTAAATG TGTGTGT AGGAGA TGTAAAG GTATGTG |
| seal-cell2one | /5Phos/GTTGAAG GAGAAAG GTTTGT TGATAGA TGTTTGA |
| seal-cell3one | /5Phos/GTGTTTG GTTATGT AGTGGA GTAGAGA GAGTTAG |
| seal-cell4one | /5Phos/AGTTTTG TGAAATG GAATGA TGATTGT GAGATTG |
| instru11 | AGAAAGA TGTTTGA GTAAATG TGTGTGT |
| instru12 | AGAAAGA GAGTTAG GTTGAAG GAGAAAG |
| instru13 | AGAAAGA GAGATTG GTGTTTG GTTATGT |
| instru14 | AGAAAGA GATTTGA AGTTTTG TGAAATG |
| instru51 | ACACACA CATTTAC TCAAACA TCTTTCT |
| instru52 | CTTTCTC CTTCAAC CTAACTC TCTTTCT |
| instru53 | ACATAAC CAAACAC CAATCTC TCTTTCT |
| instru54 | CATTTCA CAAAACT TCAAATC TCTTTCT |

#### Sequences for M13 registers

### M13.1

M13 sequence window:

GACTGGTATAATGAGCCAGTTCTTAAAATCGCATAAGGTAATTCACAATGATTAAAGTTGAAATTAAACCA
TCTCAAGCCCAATTTACTACTCGTTCTGGTGTTTCTCGTCAGGGCAAGCCTTATTCACTGAATGAGCAGCT
TTGTTACGTTGATTTGGGTAATGAATATCCGGTTCTTGTCAAGATTACTCTTGATGAAGGTCAGCCAGCCT
ATGCGCCTGGTCTGTACACCGTTCATCTGTCCTCTTTCAAAGTTGGTCAGTT

| | |
|---|---|
| leftadapR1S1 | /5Phos/TAAGAACTGGCTCATTATACCAGTC CAAAACAAAACCTCA CCGAGCCCACGAGAC |
| leftadapR1S2 | /5Phos/TAAGAACTGGCTCATTATACCAGTC CAACATATCAATTCA CCGAGCCCACGAGAC |
| leftadapR1S3 | /5Phos/TAAGAACTGGCTCATTATACCAGTC CATTATTCAAACCCA CCGAGCCCACGAGAC |
| leftadapR1S4 | /5Phos/TAAGAACTGGCTCATTATACCAGTC CACTTCATAAATCCA CCGAGCCCACGAGAC |
| leftadapR1S5 | /5Phos/TAAGAACTGGCTCATTATACCAGTC CAACTCCTAATATCA CCGAGCCCACGAGAC |
| leftadapR1S6 | /5Phos/TAAGAACTGGCTCATTATACCAGTC CAACCATACTAAACA CCGAGCCCACGAGAC |
| leftadapR1S7 | /5Phos/TAAGAACTGGCTCATTATACCAGTC CATTCCTACATTTCA CCGAGCCCACGAGAC |
| leftadapR1S8 | /5Phos/TAAGAACTGGCTCATTATACCAGTC CAAATCTTCATCCCA CCGAGCCCACGAGAC |
| leftadapR1S9 | /5Phos/TAAGAACTGGCTCATTATACCAGTC CAAACACTCTATTCA CCGAGCCCACGAGAC |
| leftadapR1S10 | /5Phos/TAAGAACTGGCTCATTATACCAGTC CAACTCAAACATACA CCGAGCCCACGAGAC |
| leftadapR1S11 | /5Phos/TAAGAACTGGCTCATTATACCAGTC CATACCCTTTTCTCA CCGAGCCCACGAGAC |
| leftadapR1S12 | /5Phos/TAAGAACTGGCTCATTATACCAGTC CATCATACCTACTCA CCGAGCCCACGAGAC |
| leftadapR1S13 | /5Phos/TAAGAACTGGCTCATTATACCAGTC CAAAACTCTCTCTCA CCGAGCCCACGAGAC |
| leftadapR1S14 | /5Phos/TAAGAACTGGCTCATTATACCAGTC CAAACCCAACTCACA CCGAGCCCACGAGAC |
| leftadapR1S15 | /5Phos/TAAGAACTGGCTCATTATACCAGTC CATTCTCCCACCTCA CCGAGCCCACGAGAC |
| leftadapR1S16 | /5Phos/TAAGAACTGGCTCATTATACCAGTC CATATCTAATCTCCA CCGAGCCCACGAGAC |
| rightadapor | TCGTCGGCAGCGTC AGAACCGGATATTCAT |

**M13.1 sequences for the binary counting program**

| | |
|---|---|
| reg1Cell1OneLeft | /5Phos/ACCTTAT GCGATTT |
| reg1Cell1OneRight | /5Phos/TCAAGTT TAATCATT GTGAATT |
| reg1Cell1ZeroLeft | /5Phos/GTGAATT ACCTTAT GCGATTT |
| reg1Cell1ZeroRight | /5Phos/TCAACTT TAATCATT |
| reg1Cell2OneLeft | /5Phos/TGAGATG GTTTAATT |
| reg1Cell2OneRight | /5Phos/ACGAGT GGTAAATT GGGCT |
| reg1Cell2ZeroLeft | /5Phos/GGGCT TGAGATG GTTTAATT |
| reg1Cell2ZeroRight | /5Phos/ACGAGT AGTAAATT |
| reg1Cell3OneLeft | /5Phos/GAGAAAC ACCAGA |
| reg1Cell3OneRight | /5Phos/TGAAGAAG GCTTGC CCTGAC |
| reg1Cell3ZeroLeft | /5Phos/CCTGAC GAGAAAC ACCAGA |
| reg1Cell3ZeroRight | /5Phos/TGAATAAG GCTTGC |
| reg1Cell4OneLeft | /5Phos/GCTGCT CATTCAG |
| reg1Cell4OneRight | /5Phos/TACCCA AGTCAAC GTAACAAA |
| reg1Cell4ZeroLeft | /5Phos/GTAACAAA GCTGCT CATTCAG |
| reg1Cell4ZeroRight | /5Phos/TACCCA AATCAAC |
| reg1Instru11 | CTTATAACAC GTGAATT ACCTTAT GCGATTT |
| reg1Instru12 | CTTATAACAC GGGCT TGAGATG GTTTAATT TCAACTT TAATCATT |
| reg1Instru13 | CTTATAACAC CCTGAC GAGAAAC ACCAGA ACGAGT AGTAAATT |
| reg1Instru14 | CTTATAACAC GTAACAAA GCTGCT CATTCAG TGAATAAG GCTTGC |
| reg1Instru15 | CTCAATATTC GGATATTCAT TACCCA AATCAAC |
| reg1Instru21 | AAATCGC ATAAGGT AATTCAC GTGTTATAAG |
| reg1Instru22 | AATGATTA AAGTTGA AATTAAAC CATCTCA AGCCC GTGTTATAAG |
| reg1Instru23 | AATTTACT ACTCGT TCTGGT GTTTCTC GTCAGG GTGTTATAAG |
| reg1Instru24 | GCAAGC CTTATTCA CTGAATG AGCAGC TTTGTTAC GTGTTATAAG |
| reg1Instru25 | GTTGATT TGGGTA ATGAATATCC GAATATTGAG |
| reg1Instru41 | TAATCATT GTGAATT ACCTTAT |
| reg1Instru42 | AGTAAATT GGGCT TGAGATG |
| reg1Instru43 | GCTTGC CCTGAC GAGAAAC |
| reg1Instru44 | AATCAAC GTAACAAA GCTGCT |
| reg1Instru61 | AAATCGC ATAAGGT AATTCAC |
| reg1Instru62 | AATTAAAC CATCTCA AGCCC |
| reg1Instru63 | TCTGGT GTTTCTC GTCAGG |
| reg1Instru64 | CTGAATG AGCAGC TTTGTTAC |

**M13.1 sequences for the rule110 program**

| | |
|---|---|
| 110R1Cell1One | TCAACTT TAATCATT GTGAATT ACCTTAT |
| 110R1Cell1ZeroLeft | /5Phos/GTGAATT ACCTTGT GCGATTT |
| 110R1Cell1ZeroRight | /5Phos/TCAACTT TAATCATT |
| 110R1Cell2One | ACGAGT AGTAAATT GGGCT TGAGATG |
| 110R1Cell2ZeroLeft | /5Phos/GGGCT TGAGGTG GTTTAATT |
| 110R1Cell2ZeroRight | /5Phos/ACGAGT AGTAAATT |
| 110R1Cell3One | TGAATAAG GCTTGC CCTGAC GAGAAAC |
| 110R1Cell3ZeroLeft | /5Phos/CCTGAC GGGAAAC ACCAGA |
| 110R1Cell3ZeroRight | /5Phos/TGAATAAG GCTTGC |
| 110R1Cell4One | TACCCA AATCAAC GTAACAAA GCTGCT |
| 110R1Cell4ZeroLeft | /5Phos/GTAACAAA GGTGCT CATTCAG |
| 110R1Cell4ZeroRight | /5Phos/TACCCA AATCAAC |
| sealR1Cell1 | /5Phos/TCAACTT TAATCATT GTGAATT ACCTTAT GCGATTT |
| sealR1Cell2 | /5Phos/ACGAGT AGTAAATT GGGCT TGAGATG GTTTAATT |
| sealR1Cell3 | /5Phos/TGAATAAG GCTTGC CCTGAC GAGAAAC ACCAGA |
| sealR1Cell4 | /5Phos/TACCCA AATCAAC GTAACAAA GCTGCT CATTCAG |
| 110R1Instru11 | TCTATCAATC GTTTAATT TCAACTT TAATCATT |
| 110R1Instru12 | TCTATCAATC ACCAGA ACGAGT AGTAAATT |
| 110R1Instru13 | TCTATCAATC CATTCAG TGAATAAG GCTTGC |
| 110R1Instru14 | TCTATCAATC GGATATTCAT TACCCA AATCAAC |
| 110R1Instru51 | AATGATTA AAGTTGA AATTAAAC GATTGATAGA |
| 110R1Instru52 | AATTTACT ACTCGT TCTGGT GATTGATAGA |
| 110R1Instru53 | GCAAGC CTTATTCA CTGAATG GATTGATAGA |
| 110R1Instru54 | GTTGATT TGGGTA ATGAATATCC GATTGATAGA |

**M13.2**

M13 sequence window:

CCTCTGTAGCCGTTGCTACCCTCGTTCCGATGCTGTCTTTCGCTGCTGAGGGTGACGATCCCGCAAAAGCG
GCCTTTAACTCCCTGCAAGCCTCAGCGACCGAATATATCGGTTATGCGTGGGCGATGGTTGTTGTCATTGT
CGGCGCAACTATCGGTATCAAGCTGTTTAAGAAATTCACCTCGAAAGCAAGCTGATAAACCGATACAATTA
AAGGCTCCTTTTGGAGCCTTTTTTTTGGAGATTTTCAACGTGAAAAAATTAT

| | |
|---|---|
| leftadapR2S1 | /5Phos/ACGAGGGTAGCAACGGCTACAGAGG CAAAACAAAACCTCA CCGAGCCCACGAGAC |
| leftadapR2S2 | /5Phos/ACGAGGGTAGCAACGGCTACAGAGG CAACATATCAATTCA CCGAGCCCACGAGAC |
| leftadapR2S3 | /5Phos/ACGAGGGTAGCAACGGCTACAGAGG CATTATTCAAACCCA CCGAGCCCACGAGAC |
| leftadapR2S4 | /5Phos/ACGAGGGTAGCAACGGCTACAGAGG CACTTCATAAATCCA CCGAGCCCACGAGAC |
| leftadapR2S5 | /5Phos/ACGAGGGTAGCAACGGCTACAGAGG CAACTCCTAATATCA CCGAGCCCACGAGAC |
| leftadapR2S6 | /5Phos/ACGAGGGTAGCAACGGCTACAGAGG CAACCATACTAAACA CCGAGCCCACGAGAC |
| leftadapR2S7 | /5Phos/ACGAGGGTAGCAACGGCTACAGAGG CATTCCTACATTTCA CCGAGCCCACGAGAC |
| leftadapR2S8 | /5Phos/ACGAGGGTAGCAACGGCTACAGAGG CAAATCTTCATCCCA CCGAGCCCACGAGAC |
| leftadapR2S9 | /5Phos/ACGAGGGTAGCAACGGCTACAGAGG CAAACACTCTATTCA CCGAGCCCACGAGAC |
| leftadapR2S10 | /5Phos/ACGAGGGTAGCAACGGCTACAGAGG CAACTCAAACATACA CCGAGCCCACGAGAC |
| leftadapR2S11 | /5Phos/ACGAGGGTAGCAACGGCTACAGAGG CATACCCTTTTCTCA CCGAGCCCACGAGAC |
| leftadapR2S12 | /5Phos/ACGAGGGTAGCAACGGCTACAGAGG CATCATACCTACTCA CCGAGCCCACGAGAC |
| leftadapR2S13 | /5Phos/ACGAGGGTAGCAACGGCTACAGAGG CAAAACTCTCTCTCA CCGAGCCCACGAGAC |
| leftadapR2S14 | /5Phos/ACGAGGGTAGCAACGGCTACAGAGG CAAACCCAACTCACA CCGAGCCCACGAGAC |
| leftadapR2S15 | /5Phos/ACGAGGGTAGCAACGGCTACAGAGG CATTCTCCCACCTCA CCGAGCCCACGAGAC |
| leftadapR2S16 | /5Phos/ACGAGGGTAGCAACGGCTACAGAGG CATATCTAATCTCCA CCGAGCCCACGAGAC |
| rightadapor | TCGTCGGCAGCGTC GCTTGATACCGATAGT |

**M13.2 sequences for the binary counting program**

| | |
|---|---|
| reg2Cell1OneLeft | /5Phos/GACAGC ATCGGA |
| reg2Cell1OneRight | /5Phos/TCACCC GCAGCA GCGAAA |
| reg2Cell1ZeroLeft | /5Phos/GCGAAA GACAGC ATCGGA |
| reg2Cell1ZeroRight | /5Phos/TCACCC TCAGCA |
| reg2Cell2OneLeft | /5Phos/TTTGCG GGATCG |
| reg2Cell2OneRight | /5Phos/AGGGAG GTAAAGG CCGCT |
| reg2Cell2ZeroLeft | /5Phos/CCGCT TTTGCG GGATCG |
| reg2Cell2ZeroRight | /5Phos/AGGGAG TTAAAGG |
| reg2Cell3OneLeft | /5Phos/GCTGAG GCTTGC |
| reg2Cell3OneRight | /5Phos/GCATAAG CGATATAT TCGGTC |
| reg2Cell3ZeroLeft | /5Phos/TCGGTC GCTGAG GCTTGC |
| reg2Cell3ZeroRight | /5Phos/GCATAAC CGATATAT |
| reg2Cell4OneLeft | /5Phos/ACCATC GCCCAC |
| reg2Cell4OneRight | /5Phos/TGCGG CGACAAT GACAACA |
| reg2Cell4ZeroLeft | /5Phos/GACAACA ACCATC GCCCAC |
| reg2Cell4ZeroRight | /5Phos/TGCGC CGACAAT |
| reg2Instru11 | ACTTTCTACC GCGAAA GACAGC ATCGGA |
| reg2Instru12 | ACTTTCTACC CCGCT TTTGCG GGATCG TCACCC TCAGCA |
| reg2Instru13 | ACTTTCTACC TCGGTC GCTGAG GCTTGC AGGGAG TTAAAGG |
| reg2Instru14 | ACTTTCTACC GACAACA ACCATC GCCCAC GCATAAC CGATATAT |
| reg2Instru15 | CCTTAACTAT TACCGATAGT TGCGC CGACAAT |
| reg2Instru21 | TCCGAT GCTGTC TTTCGC GGTAGAAAGT |
| reg2Instru22 | TGCTGA GGGTGA CGATCC CGCAAA AGCGG GGTAGAAAGT |
| reg2Instru23 | CCTTTAA CTCCCT GCAAGC CTCAGC GACCGA GGTAGAAAGT |
| reg2Instru24 | ATATATCG GTTATGC GTGGGC GATGGT TGTTGTC GGTAGAAAGT |
| reg2Instru25 | ATTGTCG GCGCA ACTATCGGTA ATAGTTAAGG |
| reg2Instru41 | TCAGCA GCGAAA GACAGC |
| reg2Instru42 | TTAAAGG CCGCT TTTGCG |
| reg2Instru43 | CGATATAT TCGGTC GCTGAG |
| reg2Instru44 | CGACAAT GACAACA ACCATC |
| reg2Instru61 | TCCGAT GCTGTC TTTCGC |
| reg2Instru62 | CGATCC CGCAAA AGCGG |
| reg2Instru63 | GCAAGC CTCAGC GACCGA |
| reg2Instru64 | GTGGGC GATGGT TGTTGTC |

##### M13.2 sequences for the rule110 program

| | |
|---|---|
| 110R2Cell1One | TCACCC TCAGCA GCGAAA GACAGC |
| 110R2Cell1ZeroLeft | /5Phos/GCGAAA GACGGC ATCGGA |
| 110R2Cell1ZeroRight | /5Phos/TCACCC TCAGCA |
| 110R2Cell2One | AGGGAG TTAAAGG CCGCT TTTGCG |
| 110R2Cell2ZeroLeft | /5Phos/CCGCT GTTGCG GGATCG |
| 110R2Cell2ZeroRight | /5Phos/AGGGAG TTAAAGG |
| 110R2Cell3One | GCATAAC CGATATAT TCGGTC GCTGAG |
| 110R2Cell3ZeroLeft | /5Phos/TCGGTC GGTGAG GCTTGC |
| 110R2Cell3ZeroRight | /5Phos/GCATAAC CGATATAT |
| 110R2Cell4One | TGCGC CGACAAT GACAACA ACCATC |
| 110R2Cell4ZeroLeft | /5Phos/GACAACA ACCGTC GCCCAC |
| 110R2Cell4ZeroRight | /5Phos/TGCGC CGACAAT |
| sealR2Cell1 | /5Phos/TCACCC TCAGCA GCGAAA GACAGC ATCGGA |
| sealR2Cell2 | /5Phos/AGGGAG TTAAAGG CCGCT TTTGCG GGATCG |
| sealR2Cell3 | /5Phos/GCATAAC CGATATAT TCGGTC GCTGAG GCTTGC |
| sealR2Cell4 | /5Phos/TGCGC CGACAAT GACAACA ACCATC GCCCAC |
| 110R2Instru11 | CATCTATACA GGATCG TCACCC TCAGCA |
| 110R2Instru12 | CATCTATACA GCTTGC AGGGAG TTAAAGG |
| 110R2Instru13 | CATCTATACA GCCCAC GCATAAC CGATATAT |
| 110R2Instru14 | CATCTATACA TACCGATAGT TGCGC CGACAAT |
| 110R2Instru51 | TGCTGA GGGTGA CGATCC TGTATAGATG |
| 110R2Instru52 | CCTTTAA CTCCCT GCAAGC TGTATAGATG |
| 110R2Instru53 | ATATATCG GTTATGC GTGGGC TGTATAGATG |
| 110R2Instru54 | ATTGTCG GCGCA ACTATCGGTA TGTATAGATG |

**M13.3**

M13 sequence window:

TATTCGCAATTCCTTTAGTTGTTCCTTTCTATTCTCACTCCGCTGAAACTGTTGAAAGTTGTTTAGCAAAA
TCCCATACAGAAAATTCATTTACTAACGTCTGGAAAGACGACAAAACTTTAGATCGTTACGCTAACTATGA
GGGCTGTCTGTGGAATGCTACAGGCGTTGTAGTTTGTACTGGTGACGAAACTCAGTGTTACGGTACATGGG
TTCCTATTGGGCTTGCTATCCCTGAAAATGAGGGTGGTGGCTCTGAGGGTGG

| | |
|---|---|
| leftadapR3S1 | /5Phos/GGAACAACTAAAGGAATTGCGAATA CAAAACAAAACCTCA CCGAGCCCACGAGAC |
| leftadapR3S2 | /5Phos/GGAACAACTAAAGGAATTGCGAATA CAACATATCAATTCA CCGAGCCCACGAGAC |
| leftadapR3S3 | /5Phos/GGAACAACTAAAGGAATTGCGAATA CATTATTCAAACCCA CCGAGCCCACGAGAC |
| leftadapR3S4 | /5Phos/GGAACAACTAAAGGAATTGCGAATA CACTTCATAAATCCA CCGAGCCCACGAGAC |
| leftadapR3S5 | /5Phos/GGAACAACTAAAGGAATTGCGAATA CAACTCCTAATATCA CCGAGCCCACGAGAC |
| leftadapR3S6 | /5Phos/GGAACAACTAAAGGAATTGCGAATA CAACCATACTAAACA CCGAGCCCACGAGAC |
| leftadapR3S7 | /5Phos/GGAACAACTAAAGGAATTGCGAATA CATTCCTACATTTCA CCGAGCCCACGAGAC |
| leftadapR3S8 | /5Phos/GGAACAACTAAAGGAATTGCGAATA CAAATCTTCATCCCA CCGAGCCCACGAGAC |
| leftadapR3S9 | /5Phos/GGAACAACTAAAGGAATTGCGAATA CAAACACTCTATTCA CCGAGCCCACGAGAC |
| leftadapR3S10 | /5Phos/GGAACAACTAAAGGAATTGCGAATA CAACTCAAACATACA CCGAGCCCACGAGAC |
| leftadapR3S11 | /5Phos/GGAACAACTAAAGGAATTGCGAATA CATACCCTTTTCTCA CCGAGCCCACGAGAC |
| leftadapR3S12 | /5Phos/GGAACAACTAAAGGAATTGCGAATA CATCATACCTACTCA CCGAGCCCACGAGAC |
| leftadapR3S13 | /5Phos/GGAACAACTAAAGGAATTGCGAATA CAAAACTCTCTCTCA CCGAGCCCACGAGAC |
| leftadapR3S14 | /5Phos/GGAACAACTAAAGGAATTGCGAATA CAAACCCAACTCACA CCGAGCCCACGAGAC |
| leftadapR3S15 | /5Phos/GGAACAACTAAAGGAATTGCGAATA CATTCTCCCACCTCA CCGAGCCCACGAGAC |
| leftadapR3S16 | /5Phos/GGAACAACTAAAGGAATTGCGAATA CATATCTAATCTCCA CCGAGCCCACGAGAC |
| rightadapor | TCGTCGGCAGCGTC CAAACTACAACGCCTG |

##### M13.3 sequences for the binary counting program

| | |
|---|---|
| reg3Cell1OneLeft | /5Phos/GAGTGAG AATAGAAA |
| reg3Cell1OneRight | /5Phos/ACTTTCG ACAGTTT CAGCG |
| reg3Cell1ZeroLeft | /5Phos/CAGCG GAGTGAG AATAGAAA |
| reg3Cell1ZeroRight | /5Phos/ACTTTCA ACAGTTT |
| reg3Cell2OneLeft | /5Phos/GATTTTG CTAAACA |
| reg3Cell2OneRight | /5Phos/TAGTAAAG GAATTTTC TGTATGG |
| reg3Cell2ZeroLeft | /5Phos/TGTATGG GATTTTG CTAAACA |
| reg3Cell2ZeroRight | /5Phos/TAGTAAAT GAATTTTC |
| reg3Cell3OneLeft | /5Phos/TCTTTCC AGACGT |
| reg3Cell3OneRight | /5Phos/TAACGAG CTAAAGTT TTGTCG |
| reg3Cell3ZeroLeft | /5Phos/TTGTCG TCTTTCC AGACGT |
| reg3Cell3ZeroRight | /5Phos/TAACGAT CTAAAGTT |
| reg3Cell4OneLeft | /5Phos/CTCATAG TTAGCG |
| reg3Cell4OneRight | /5Phos/TAGCGTT CCACAG ACAGCC |
| reg3Cell4ZeroLeft | /5Phos/ACAGCC CTCATAG TTAGCG |
| reg3Cell4ZeroRight | /5Phos/TAGCATT CCACAG |
| reg3Instru11 | TAATCTTCCA CAGCG GAGTGAG AATAGAAA |
| reg3Instru12 | TAATCTTCCA TGTATGG GATTTTG CTAAACA ACTTTCA ACAGTTT |
| reg3Instru13 | TAATCTTCCA TTGTCG TCTTTCC AGACGT TAGTAAAT GAATTTTC |
| reg3Instru14 | TAATCTTCCA ACAGCC CTCATAG TTAGCG TAACGAT CTAAAGTT |
| reg3Instru15 | TTATACCTCT ACAACGCCTG TAGCATT CCACAG |
| reg3Instru21 | TTTCTATT CTCACTC CGCTG TGGAAGATTA |
| reg3Instru22 | AAACTGT TGAAAGT TGTTTAG CAAAATC CCATACA TGGAAGATTA |
| reg3Instru23 | GAAAATTC ATTTACTA ACGTCT GGAAAGA CGACAA TGGAAGATTA |
| reg3Instru24 | AACTTTAG ATCGTTA CGCTAA CTATGAG GGCTGT TGGAAGATTA |
| reg3Instru25 | CTGTGG AATGCTA CAGGCGTTGT AGAGGTATAA |
| reg3Instru41 | ACAGTTT CAGCG GAGTGAG |
| reg3Instru42 | GAATTTTC TGTATGG GATTTTG |
| reg3Instru43 | CTAAAGTT TTGTCG TCTTTCC |
| reg3Instru44 | CCACAG ACAGCC CTCATAG |
| reg3Instru61 | TTTCTATT CTCACTC CGCTG |
| reg3Instru62 | TGTTTAG CAAAATC CCATACA |
| reg3Instru63 | ACGTCT GGAAAGA CGACAA |
| reg3Instru64 | CGCTAA CTATGAG GGCTGT |

**M13.3 sequences for the rule110 program**

| | |
|---|---|
| 110R3Cell1One | ACTTTCA ACAGTTT CAGCG GAGTGAG |
| 110R3Cell1ZeroLeft | /5Phos/CAGCG GAGGGAG AATAGAAA |
| 110R3Cell1ZeroRight | /5Phos/ACTTTCA ACAGTTT |
| 110R3Cell2One | TAGTAAAT GAATTTTC TGTATGG GATTTTG |
| 110R3Cell2ZeroLeft | /5Phos/TGTATGG GAGTTTG CTAAACA |
| 110R3Cell2ZeroRight | /5Phos/TAGTAAAT GAATTTTC |
| 110R3Cell3One | TAACGAT CTAAAGTT TTGTCG TCTTTCC |
| 110R3Cell3ZeroLeft | /5Phos/TTGTCG TCTTTGC AGACGT |
| 110R3Cell3ZeroRight | /5Phos/TAACGAT CTAAAGTT |
| 110R3Cell4One | TAGCATT CCACAG ACAGCC CTCATAG |
| 110R3Cell4ZeroLeft | /5Phos/ACAGCC CTCAGAG TTAGCG |
| 110R3Cell4ZeroRight | /5Phos/TAGCATT CCACAG |
| sealR3Cell1 | /5Phos/ACTTTCA ACAGTTT CAGCG GAGTGAG AATAGAAA |
| sealR3Cell2 | /5Phos/TAGTAAAT GAATTTTC TGTATGG GATTTTG CTAAACA |
| sealR3Cell3 | /5Phos/TAACGAT CTAAAGTT TTGTCG TCTTTCC AGACGT |
| sealR3Cell4 | /5Phos/TAGCATT CCACAG ACAGCC CTCATAG TTAGCG |
| 110R3Instru11 | CTTTTCCAAT CTAAACA ACTTTCA ACAGTTT |
| 110R3Instru12 | CTTTTCCAAT AGACGT TAGTAAAT GAATTTTC |
| 110R3Instru13 | CTTTTCCAAT TTAGCG TAACGAT CTAAAGTT |
| 110R3Instru14 | CTTTTCCAAT ACAACGCCTG TAGCATT CCACAG |
| 110R3Instru51 | AAACTGT TGAAAGT TGTTTAG ATTGGAAAAG |
| 110R3Instru52 | GAAAATTC ATTTACTA ACGTCT ATTGGAAAAG |
| 110R3Instru53 | AACTTTAG ATCGTTA CGCTAA ATTGGAAAAG |
| 110R3Instru54 | CTGTGG AATGCTA CAGGCGTTGT ATTGGAAAAG |

**M13.4**

M13 sequence window:

TTTCCGTCAATATTTACCTTCCCTCCCTCAATCGGTTGAATGTCGCCCTTTTGTCTTTGGCGCTGGTAAAC
CATATGAATTTTCTATTGATTGTGACAAAATAAACTTATTCCGTGGTGTCTTTGCGTTTCTTTTATATGTT
GCCACCTTTATGTATGTATTTTCTACGTTTGCTAACATACTGCGTAATAAGGAGTCTTAATCATGCCAGTT
CTTTTGGGTATTCCGTTATTATTGCGTTTCCTCGGTTTCCTTCTGGTAACTT

| | |
|---|---|
| leftadapR4S1 | /5Phos/GAGGGAAGGTAAATATTGACGGAAA CAAAACAAAACCTCA CCGAGCCCACGAGAC |
| leftadapR4S2 | /5Phos/GAGGGAAGGTAAATATTGACGGAAA CAACATATCAATTCA CCGAGCCCACGAGAC |
| leftadapR4S3 | /5Phos/GAGGGAAGGTAAATATTGACGGAAA CATTATTCAAACCCA CCGAGCCCACGAGAC |
| leftadapR4S4 | /5Phos/GAGGGAAGGTAAATATTGACGGAAA CACTTCATAAATCCA CCGAGCCCACGAGAC |
| leftadapR4S5 | /5Phos/GAGGGAAGGTAAATATTGACGGAAA CAACTCCTAATATCA CCGAGCCCACGAGAC |
| leftadapR4S6 | /5Phos/GAGGGAAGGTAAATATTGACGGAAA CAACCATACTAAACA CCGAGCCCACGAGAC |
| leftadapR4S7 | /5Phos/GAGGGAAGGTAAATATTGACGGAAA CATTCCTACATTTCA CCGAGCCCACGAGAC |
| leftadapR4S8 | /5Phos/GAGGGAAGGTAAATATTGACGGAAA CAAATCTTCATCCCA CCGAGCCCACGAGAC |
| leftadapR4S9 | /5Phos/GAGGGAAGGTAAATATTGACGGAAA CAAACACTCTATTCA CCGAGCCCACGAGAC |
| leftadapR4S10 | /5Phos/GAGGGAAGGTAAATATTGACGGAAA CAACTCAAACATACA CCGAGCCCACGAGAC |
| leftadapR4S11 | /5Phos/GAGGGAAGGTAAATATTGACGGAAA CATACCCTTTTCTCA CCGAGCCCACGAGAC |
| leftadapR4S12 | /5Phos/GAGGGAAGGTAAATATTGACGGAAA CATCATACCTACTCA CCGAGCCCACGAGAC |
| leftadapR4S13 | /5Phos/GAGGGAAGGTAAATATTGACGGAAA CAAAACTCTCTCTCA CCGAGCCCACGAGAC |
| leftadapR4S14 | /5Phos/GAGGGAAGGTAAATATTGACGGAAA CAAACCCAACTCACA CCGAGCCCACGAGAC |
| leftadapR4S15 | /5Phos/GAGGGAAGGTAAATATTGACGGAAA CATTCTCCCACCTCA CCGAGCCCACGAGAC |
| leftadapR4S16 | /5Phos/GAGGGAAGGTAAATATTGACGGAAA CATATCTAATCTCCA CCGAGCCCACGAGAC |
| rightadapor | TCGTCGGCAGCGTC ATACCCAAAAGAACTG |

**M13.4 sequences for the binary counting program**

| | |
|---|---|
| reg4Cell1OneLeft | /5Phos/CGACATTCAA CCGATTGAGG |
| reg4Cell1OneRight | /5Phos/ACCAGC GGCAAAGACA AAAGGG |
| reg4Cell1ZeroLeft | /5Phos/AAAGGG CGACATTCAA CCGATTGAGG |
| reg4Cell1ZeroRight | /5Phos/ACCAGC GCCAAAGACA |
| reg4Cell2OneLeft | /5Phos/TCAATAGAAAAT TCATATGGTTT |
| reg4Cell2OneRight | /5Phos/CACGGA AGAAGTTTATTTT GTCACAA |
| reg4Cell2ZeroLeft | /5Phos/GTCACAA TCAATAGAAAAT TCATATGGTTT |
| reg4Cell2ZeroRight | /5Phos/CACGGA ATAAGTTTATTTT |
| reg4Cell3OneLeft | /5Phos/TATAAAAGAAAC GCAAAGACAC |
| reg4Cell3OneRight | /5Phos/AAATACAG ACATAAAGGTG GCAACA |
| reg4Cell3ZeroLeft | /5Phos/GCAACA TATAAAAGAAAC GCAAAGACAC |
| reg4Cell3ZeroRight | /5Phos/AAATACAT ACATAAAGGTG |
| reg4Cell4OneLeft | /5Phos/CAGTATGTTAG CAAACGTAGA |
| reg4Cell4OneRight | /5Phos/GCATGA GTAAGACTCCT TATTACG |
| reg4Cell4ZeroLeft | /5Phos/TATTACG CAGTATGTTAG CAAACGTAGA |
| reg4Cell4ZeroRight | /5Phos/GCATGA TTAAGACTCCT |
| reg4Instru11 | CCTTACAAAA AAAGGG CGACATTCAA CCGATTGAGG |
| reg4Instru12 | CCTTACAAAA GTCACAA TCAATAGAAAAT TCATATGGTTT ACCAGC GCCAAAGACA |
| reg4Instru13 | CCTTACAAAA GCAACA TATAAAAGAAAC GCAAAGACAC CACGGA ATAAGTTTATTTT |
| reg4Instru14 | CCTTACAAAA TATTACG CAGTATGTTAG CAAACGTAGA AAATACAT ACATAAAGGTG |
| reg4Instru15 | AATCCTACTC AAAAGAACTG GCATGA TTAAGACTCCT |
| reg4Instru21 | CCTCAATCGG TTGAATGTCG CCCTTT TTTTGTAAGG |
| reg4Instru22 | TGTCTTTGGC GCTGGT AAACCATATGA ATTTTCTATTGA TTGTGAC TTTTGTAAGG |
| reg4Instru23 | AAAATAAACTTAT TCCGTG GTGTCTTTGC GTTTCTTTTATA TGTTGC TTTTGTAAGG |
| reg4Instru24 | CACCTTTATGT ATGTATTT TCTACGTTTG CTAACATACTG CGTAATA TTTTGTAAGG |
| reg4Instru25 | AGGAGTCTTAA TCATGC CAGTTCTTTT GAGTAGGATT |
| reg4Instru41 | GCCAAAGACA AAAGGG CGACATTCAA |
| reg4Instru42 | AAAAGTTTATTTT GTCACAA TCAATAGAAAAT |
| reg4Instru43 | ACATAAAGGTG GCAACA TATAAAAGAAAC |
| reg4Instru44 | TTAAGACTCCT TATTACG CAGTATGTTAG |
| reg4Instru61 | CCTCAATCGG TTGAATGTCG CCCTTT |
| reg4Instru62 | AAACCATATGA ATTTTCTATTGA TTGTGAC |
| reg4Instru63 | GTGTCTTTGC GTTTCTTTTATA TGTTGC |
| reg4Instru64 | TCTACGTTTG CTAACATACTG CGTAATA |

**M13.5**

M13 sequence window:

TGTTCGGCTATCTGCTTACTTTTCTTAAAAAGGGCTTCGGTAAGATAGCTATTGCTATTTCATTGTTTCTT
GCTCTTATTATTGGGCTTAACTCAATTCTTGTGGGTTATCTCTCTGATATTAGCGCTCAATTACCCTCTGA
CTTTGTTCAGGGTGTTCAGTTAATTCTCCCGTCTAATGCGCTTCCCTGTTTTTATGTTATTCTCTCTGTAA
AGGCTGCTATTTTCATTTTTGACGTTAAACAAAAAATCGTTTCTTATTTGGA

| | |
|---|---|
| leftadapR5S1 | /5Phos/AGAAAAGTAAGCAGATAGCCGAACA CAAAACAAAACCTCA CCGAGCCCACGAGAC |
| leftadapR5S2 | /5Phos/AGAAAAGTAAGCAGATAGCCGAACA CAACATATCAATTCA CCGAGCCCACGAGAC |
| leftadapR5S3 | /5Phos/AGAAAAGTAAGCAGATAGCCGAACA CATTATTCAAACCCA CCGAGCCCACGAGAC |
| leftadapR5S4 | /5Phos/AGAAAAGTAAGCAGATAGCCGAACA CACTTCATAAATCCA CCGAGCCCACGAGAC |
| leftadapR5S5 | /5Phos/AGAAAAGTAAGCAGATAGCCGAACA CAACTCCTAATATCA CCGAGCCCACGAGAC |
| leftadapR5S6 | /5Phos/AGAAAAGTAAGCAGATAGCCGAACA CAACCATACTAAACA CCGAGCCCACGAGAC |
| leftadapR5S7 | /5Phos/AGAAAAGTAAGCAGATAGCCGAACA CATTCCTACATTTCA CCGAGCCCACGAGAC |
| leftadapR5S8 | /5Phos/AGAAAAGTAAGCAGATAGCCGAACA CAAATCTTCATCCCA CCGAGCCCACGAGAC |
| leftadapR5S9 | /5Phos/AGAAAAGTAAGCAGATAGCCGAACA CAAACACTCTATTCA CCGAGCCCACGAGAC |
| leftadapR5S10 | /5Phos/AGAAAAGTAAGCAGATAGCCGAACA CAACTCAAACATACA CCGAGCCCACGAGAC |
| leftadapR5S11 | /5Phos/AGAAAAGTAAGCAGATAGCCGAACA CATACCCTTTTCTCA CCGAGCCCACGAGAC |
| leftadapR5S12 | /5Phos/AGAAAAGTAAGCAGATAGCCGAACA CATCATACCTACTCA CCGAGCCCACGAGAC |
| leftadapR5S13 | /5Phos/AGAAAAGTAAGCAGATAGCCGAACA CAAAACTCTCTCTCA CCGAGCCCACGAGAC |
| leftadapR5S14 | /5Phos/AGAAAAGTAAGCAGATAGCCGAACA CAAACCCAACTCACA CCGAGCCCACGAGAC |
| leftadapR5S15 | /5Phos/AGAAAAGTAAGCAGATAGCCGAACA CATTCTCCCACCTCA CCGAGCCCACGAGAC |
| leftadapR5S16 | /5Phos/AGAAAAGTAAGCAGATAGCCGAACA CATATCTAATCTCCA CCGAGCCCACGAGAC |
| rightadapor | TCGTCGGCAGCGTC ATAGCAGCCTTTACAG |

**M13.5 sequences for the binary counting program**

| | |
|---|---|
| reg5Cell1OneLeft | /5Phos/TCTTACCGAA GCCCTTTTTA |
| reg5Cell1OneRight | /5Phos/AGAAACG ATGAA ATAGCA ATAGCTA |
| reg5Cell1ZeroLeft | /5Phos/ATAGCTA TCTTACCGAA GCCCTTTTTA |
| reg5Cell1ZeroRight | /5Phos/AGAAACA ATGAA ATAGCA |
| reg5Cell2OneLeft | /5Phos/TTAAGCCCAA TAATAAGAGCA |
| reg5Cell2OneRight | /5Phos/GAGAGAGA ACCCACAAG AATTGAG |
| reg5Cell2ZeroLeft | /5Phos/AATTGAG TTAAGCCCAA TAATAAGAGCA |
| reg5Cell2ZeroRight | /5Phos/GAGAGATA ACCCACAAG |
| reg5Cell3OneLeft | /5Phos/GGGT AATTGAG CGCTAATATCA |
| reg5Cell3OneRight | /5Phos/CTGAACG CCCTGAACAA AGTCAGA |
| reg5Cell3ZeroLeft | /5Phos/AGTCAGA GGGT AATTGAG CGCTAATATCA |
| reg5Cell3ZeroRight | /5Phos/CTGAACA CCCTGAACAA |
| reg5Cell4OneLeft | /5Phos/CGCATTAGAC GGGAGAATTAA |
| reg5Cell4OneRight | /5Phos/AGAGAAGA ACATAAAAACA GGGAAG |
| reg5Cell4ZeroLeft | /5Phos/GGGAAG CGCATTAGAC GGGAGAATTAA |
| reg5Cell4ZeroRight | /5Phos/AGAGAATA ACATAAAAACA |
| reg5Instru11 | TTCTCCTATA ATAGCTA TCTTACCGAA GCCCTTTTTA |
| reg5Instru12 | TTCTCCTATA AATTGAG TTAAGCCCAA TAATAAGAGCA AGAAACA ATGAAATAGCA |
| reg5Instru13 | TTCTCCTATA AGTCAGA GGGTAATTGAG CGCTAATATCA GAGAGATA ACCCACAAG |
| reg5Instru14 | TTCTCCTATA GGGAAG CGCATTAGAC GGGAGAATTAA CTGAACA CCCTGAACAA |
| reg5Instru15 | TCCCATAATT GCCTTTACAG AGAGAATA ACATAAAAACA |
| reg5Instru21 | TAAAAAGGGC TTCGGTAAGA TAGCTAT TATAGGAGAA |
| reg5Instru22 | TGCTATTTCAT TGTTTCT TGCTCTTATTA TTGGGCTTAA CTCAATT TATAGGAGAA |
| reg5Instru23 | CTTGTGGGT TATCTCTC TGATATTAGCG CTCAATTACCC TCTGACT TATAGGAGAA |
| reg5Instru24 | TTGTTCAGGG TGTTCAG TTAATTCTCCC GTCTAATGCG CTTCCC TATAGGAGAA |
| reg5Instru25 | TGTTTTTATGT TATTCTCT CTGTAAAGGC AATTATGGGA |
| reg5Instru41 | ATGAAATAGCA ATAGCTA TCTTACCGAA |
| reg5Instru42 | ACCCACAAG AATTGAG TTAAGCCCAA |
| reg5Instru43 | CCCTGAACAA AGTCAGA GGGTAATTGAG |
| reg5Instru44 | ACATAAAAACA GGGAAG CGCATTAGAC |
| reg5Instru61 | TAAAAAGGGC TTCGGTAAGA TAGCTAT |
| reg5Instru62 | TGCTCTTATTA TTGGGCTTAA CTCAATT |
| reg5Instru63 | TGATATTAGCG CTCAATTACCC TCTGACT |
| reg5Instru64 | TTAATTCTCCC GTCTAATGCG CTTCCC |

**M13.6**

M13 sequence window:

TTGGGATAAATAATATGGCTGTTTATTTTGTAACTGGCAAATTAGGCTCTGGAAAGACGCTCGTTAGCGTT
GGTAAGATTCAGGATAAAATTGTAGCTGGGTGCAAAATAGCAACTAATCTTGATTTAAGGCTTCAAAACCT
CCCGCAAGTCGGGAGGTTCGCTAAAACGCCTCGCGTTCTTAGAATACCGGATAAGCCTTCTATATCTGATT
TGCTTGCTATTGGGCGCGGTAATGATTCCTACGATGAAAATAAAAACGGCTT

| | |
|---|---|
| leftadapR6S1 | /5Phos/TAAACAGCCATATTATTTATCCCAA CAAAACAAAACCTCA CCGAGCCCACGAGAC |
| leftadapR6S2 | /5Phos/TAAACAGCCATATTATTTATCCCAA CAACATATCAATTCA CCGAGCCCACGAGAC |
| leftadapR6S3 | /5Phos/TAAACAGCCATATTATTTATCCCAA CATTATTCAAACCCA CCGAGCCCACGAGAC |
| leftadapR6S4 | /5Phos/TAAACAGCCATATTATTTATCCCAA CACTTCATAAATCCA CCGAGCCCACGAGAC |
| leftadapR6S5 | /5Phos/TAAACAGCCATATTATTTATCCCAA CAACTCCTAATATCA CCGAGCCCACGAGAC |
| leftadapR6S6 | /5Phos/TAAACAGCCATATTATTTATCCCAA CAACCATACTAAACA CCGAGCCCACGAGAC |
| leftadapR6S7 | /5Phos/TAAACAGCCATATTATTTATCCCAA CATTCCTACATTTCA CCGAGCCCACGAGAC |
| leftadapR6S8 | /5Phos/TAAACAGCCATATTATTTATCCCAA CAAATCTTCATCCCA CCGAGCCCACGAGAC |
| leftadapR6S9 | /5Phos/TAAACAGCCATATTATTTATCCCAA CAAACACTCTATTCA CCGAGCCCACGAGAC |
| leftadapR6S10 | /5Phos/TAAACAGCCATATTATTTATCCCAA CAACTCAAACATACA CCGAGCCCACGAGAC |
| leftadapR6S11 | /5Phos/TAAACAGCCATATTATTTATCCCAA CATACCCTTTTCTCA CCGAGCCCACGAGAC |
| leftadapR6S12 | /5Phos/TAAACAGCCATATTATTTATCCCAA CATCATACCTACTCA CCGAGCCCACGAGAC |
| leftadapR6S13 | /5Phos/TAAACAGCCATATTATTTATCCCAA CAAAACTCTCTCTCA CCGAGCCCACGAGAC |
| leftadapR6S14 | /5Phos/TAAACAGCCATATTATTTATCCCAA CAAACCCAACTCACA CCGAGCCCACGAGAC |
| leftadapR6S15 | /5Phos/TAAACAGCCATATTATTTATCCCAA CATTCTCCCACCTCA CCGAGCCCACGAGAC |
| leftadapR6S16 | /5Phos/TAAACAGCCATATTATTTATCCCAA CATATCTAATCTCCA CCGAGCCCACGAGAC |
| rightadapor | TCGTCGGCAGCGTC AATCAGATATAGAAGG |

**M13.6 sequences for the binary counting program**

| | |
|---|---|
| reg6Cell1OneLeft | /5Phos/CCTAATTTGC CAGTTACAAAA |
| reg6Cell1OneRight | /5Phos/GCTAACG GGCGTCTTT CCAGAG |
| reg6Cell1ZeroLeft | /5Phos/CCAGAG CCTAATTTGC CAGTTACAAAA |
| reg6Cell1ZeroRight | /5Phos/GCTAACG AGCGTCTTT |
| reg6Cell2OneLeft | /5Phos/TTTATCCTGAA TCTTACCAAC |
| reg6Cell2OneRight | /5Phos/GCTAGTT TGCACCCAG CTACAAT |
| reg6Cell2ZeroLeft | /5Phos/CTACAAT TTTATCCTGAA TCTTACCAAC |
| reg6Cell2ZeroRight | /5Phos/GCTATTT TGCACCCAG |
| reg6Cell3OneLeft | /5Phos/AAGCCTTAAAT CAAGATTAGTT |
| reg6Cell3OneRight | /5Phos/CCCGG CTTGCGGGA GGTTTTG |
| reg6Cell3ZeroLeft | /5Phos/GGTTTTG AAGCCTTAAAT CAAGATTAGTT |
| reg6Cell3ZeroRight | /5Phos/CCCGA CTTGCGGGA |
| reg6Cell4OneLeft | /5Phos/AGGCGTTTT AGCGAACCT |
| reg6Cell4OneRight | /5Phos/CTTAGCC GGTATTCTAAGA ACGCG |
| reg6Cell4ZeroLeft | /5Phos/ACGCG AGGCGTTTT AGCGAACCT |
| reg6Cell4ZeroRight | /5Phos/CTTATCC GGTATTCTAAGA |
| reg6Instru11 | CCATTTACTC CCAGAG CCTAATTTGC CAGTTACAAAA |
| reg6Instru12 | CCATTTACTC CTACAAT TTTATCCTGAA TCTTACCAAC GCTAACG AGCGTCTTT |
| reg6Instru13 | CCATTTACTC GGTTTTG AAGCCTTAAAT CAAGATTAGTT GCTATTT TGCACCCAG |
| reg6Instru14 | CCATTTACTC ACGCG AGGCGTTTT AGCGAACCT CCCGA CTTGCGGGA |
| reg6Instru15 | TACTAATCAC ATATAGAAGG CTTATCC GGTATTCTAAGA |
| reg6Instru21 | TTTTGTAACTG GCAAATTAGG CTCTGG GAGTAAATGG |
| reg6Instru22 | AAAGACGCT CGTTAGC GTTGGTAAGA TTCAGGATAAA ATTGTAG GAGTAAATGG |
| reg6Instru23 | CTGGGTGCA AAATAGC AACTAATCTTG ATTTAAGGCTT CAAAACC GAGTAAATGG |
| reg6Instru24 | TCCCGCAAG TCGGG AGGTTCGCT AAAACGCCT CGCGT GAGTAAATGG |
| reg6Instru25 | TCTTAGAATACC GGATAAG CCTTCTATAT GTGATTAGTA |
| reg6Instru41 | AGCGTCTTT CCAGAG CCTAATTTGC |
| reg6Instru42 | TGCACCCAG CTACAAT TTTATCCTGAA |
| reg6Instru43 | CTTGCGGGA GGTTTTG AAGCCTTAAAT |
| reg6Instru44 | GGTATTCTAAGA ACGCG AGGCGTTTT |
| reg6Instru61 | TTTTGTAACTG GCAAATTAGG CTCTGG |
| reg6Instru62 | GTTGGTAAGA TTCAGGATAAA ATTGTAG |
| reg6Instru63 | AACTAATCTTG ATTTAAGGCTT CAAAACC |
| reg6Instru64 | AGGTTCGCT AAAACGCCT CGCGT |

**M13.7**

M13 sequence window:

GCTTGTTCTCGATGAGTGCGGTACTTGGTTTAATACCCGTTCTTGGAATGATAAGGAAAGACAGCCGATTA
TTGATTGGTTTCTACATGCTCGTAAATTAGGATGGGATATTATTTTTCTTGTTCAGGACTTATCTATTGTT
GATAAACAGGCGCGTTCTGCATTAGCTGAACATGTTGTTTATTGTCGTCGTCTGGACAGAATTACTTTACC
TTTTGTCGGTACTTTATATTCTCTTATTACTGGCTCGAAAATGCCTCTGCCT

| | |
|---|---|
| leftadapR7S1 | /5Phos/AGTACCGCACTCATCGAGAACAAGC CAAAACAAAACCTCA CCGAGCCCACGAGAC |
| leftadapR7S2 | /5Phos/AGTACCGCACTCATCGAGAACAAGC CAACATATCAATTCA CCGAGCCCACGAGAC |
| leftadapR7S3 | /5Phos/AGTACCGCACTCATCGAGAACAAGC CATTATTCAAACCCA CCGAGCCCACGAGAC |
| leftadapR7S4 | /5Phos/AGTACCGCACTCATCGAGAACAAGC CACTTCATAAATCCA CCGAGCCCACGAGAC |
| leftadapR7S5 | /5Phos/AGTACCGCACTCATCGAGAACAAGC CAACTCCTAATATCA CCGAGCCCACGAGAC |
| leftadapR7S6 | /5Phos/AGTACCGCACTCATCGAGAACAAGC CAACCATACTAAACA CCGAGCCCACGAGAC |
| leftadapR7S7 | /5Phos/AGTACCGCACTCATCGAGAACAAGC CATTCCTACATTTCA CCGAGCCCACGAGAC |
| leftadapR7S8 | /5Phos/AGTACCGCACTCATCGAGAACAAGC CAAATCTTCATCCCA CCGAGCCCACGAGAC |
| leftadapR7S9 | /5Phos/AGTACCGCACTCATCGAGAACAAGC CAAACACTCTATTCA CCGAGCCCACGAGAC |
| leftadapR7S10 | /5Phos/AGTACCGCACTCATCGAGAACAAGC CAACTCAAACATACA CCGAGCCCACGAGAC |
| leftadapR7S11 | /5Phos/AGTACCGCACTCATCGAGAACAAGC CATACCCTTTTCTCA CCGAGCCCACGAGAC |
| leftadapR7S12 | /5Phos/AGTACCGCACTCATCGAGAACAAGC CATCATACCTACTCA CCGAGCCCACGAGAC |
| leftadapR7S13 | /5Phos/AGTACCGCACTCATCGAGAACAAGC CAAAACTCTCTCTCA CCGAGCCCACGAGAC |
| leftadapR7S14 | /5Phos/AGTACCGCACTCATCGAGAACAAGC CAAACCCAACTCACA CCGAGCCCACGAGAC |
| leftadapR7S15 | /5Phos/AGTACCGCACTCATCGAGAACAAGC CATTCTCCCACCTCA CCGAGCCCACGAGAC |
| leftadapR7S16 | /5Phos/AGTACCGCACTCATCGAGAACAAGC CATATCTAATCTCCA CCGAGCCCACGAGAC |
| rightadapor | TCGTCGGCAGCGTC TTTAGGCAGAGGCATT |

**M13.7 sequences for the binary counting program**

| | |
|---|---|
| reg7Cell1OneLeft | /5Phos/TTATCATTCCAAGAA CGGGTATTAAACCA |
| reg7Cell1OneRight | /5Phos/AAACCGA TCAATAATCGGCTG TCTTTCC |
| reg7Cell1ZeroLeft | /5Phos/TCTTTCC TTATCATTCCAAGAA CGGGTATTAAACCA |
| reg7Cell1ZeroRight | /5Phos/AAACCAA TCAATAATCGGCTG |
| reg7Cell2OneLeft | /5Phos/ATATCCCATCCTAAT TTACGAGCATGTAG |
| reg7Cell2OneRight | /5Phos/ACAATGG ATAAGTCCTGAACAA GAAAAATA |
| reg7Cell2ZeroLeft | /5Phos/GAAAAATA ATATCCCATCCTAAT TTACGAGCATGTAG |
| reg7Cell2ZeroRight | /5Phos/ACAATAG ATAAGTCCTGAACAA |
| reg7Cell3OneLeft | /5Phos/GCTAATGCAGAACG CGCCTGTTTATCA |
| reg7Cell3OneRight | /5Phos/AGACGG CGACAATAAACAAC ATGTTCA |
| reg7Cell3ZeroLeft | /5Phos/ATGTTCA GCTAATGCAGAACG CGCCTGTTTATCA |
| reg7Cell3ZeroRight | /5Phos/AGACGA CGACAATAAACAAC |
| reg7Cell4OneLeft | /5Phos/ACCGACAAAAGGT AAAGTAATTCTGTCC |
| reg7Cell4OneRight | /5Phos/TTCGAG GCAGTAATAAGAGAA TATAAAGT |
| reg7Cell4ZeroLeft | /5Phos/TATAAAGT ACCGACAAAAGGT AAAGTAATTCTGTCC |
| reg7Cell4ZeroRight | /5Phos/TTCGAG CCAGTAATAAGAGAA |
| reg7Instru11 | CCTAATTCAA TCTTTCC TTATCATTCCAAGAA CGGGTATTAAACCA |
| reg7Instru12 | CCTAATTCAA GAAAAATA ATATCCCATCCTAAT TTACGAGCATGTAG AAACCAA TCAATAATCGGCTG |
| reg7Instru13 | CCTAATTCAA ATGTTCA GCTAATGCAGAACG CGCCTGTTTATCA ACAATAG ATAAGTCCTGAACAA |
| reg7Instru14 | CCTAATTCAA TATAAAGT ACCGACAAAAGGT AAAGTAATTCTGTCC AGACGA CGACAATAAACAAC |
| reg7Instru15 | ATTCTCACTT CAGAGGCATT TTCGAG CCAGTAATAAGAGAA |
| reg7Instru21 | TGGTTTAATACCCG TTCTTGGAATGATAA GGAAAGA TTGAATTAGG |
| reg7Instru22 | CAGCCGATTATTGA TTGGTTT CTACATGCTCGTAA ATTAGGATGGGATAT TATTTTTC TTGAATTAGG |
| reg7Instru23 | TTGTTCAGGACTTAT CTATTGT TGATAAACAGGCG CGTTCTGCATTAGC TGAACAT TTGAATTAGG |
| reg7Instru24 | GTTGTTTATTGTCG TCGTCT GGACAGAATTACTTT ACCTTTTGTCGGT ACTTTATA TTGAATTAGG |
| reg7Instru25 | TTCTCTTATTACTGG CTCGAA AATGCCTCTG AAGTGAGAAT |
| reg7Instru41 | TCAATAATCGGCTG TCTTTCC TTATCATTCCAAGAA |
| reg7Instru42 | ATAAGTCCTGAACAA GAAAAATA ATATCCCATCCTAAT |
| reg7Instru43 | CGACAATAAACAAC ATGTTCA GCTAATGCAGAACG |
| reg7Instru44 | CCAGTAATAAGAGAA TATAAAGT ACCGACAAAAGGT |
| reg7Instru61 | TGGTTTAATACCCG TTCTTGGAATGATAA GGAAAGA |
| reg7Instru62 | CTACATGCTCGTAA ATTAGGATGGGATAT TATTTTTC |
| reg7Instru63 | TGATAAACAGGCG CGTTCTGCATTAGC TGAACAT |
| reg7Instru64 | GGACAGAATTACTTT ACCTTTTGTCGGT ACTTTATA |

**M13.8**

M13 sequence window:

ATTTTGTTTTCTTGATGTTTGTTTCATCATCTTCTTTTGCTCAGGTAATTGAAATGAATAATTCGCCTCTG
CGCGATTTTGTAACTTGGTATTCAAAGCAATCAGGCGAATCCGTTATTGTTTCTCCCGATGTAAAAGGTAC
TGTTACTGTATATTCATCTGACGTTAAACCTGAAAATCTACGCAATTTCTTTATTTCTGTTTTACGTGCAA
ATAATTTTGATATGGTAGGTTCTAACCCTTCCATTATTCAGAAGTATAATCC

| | |
|---|---|
| leftadapR8S1 | /5Phos/GAAACAAACATCAAGAAAACAAAAT CAAAACAAAACCTCA CCGAGCCCACGAGAC |
| leftadapR8S2 | /5Phos/GAAACAAACATCAAGAAAACAAAAT CAACATATCAATTCA CCGAGCCCACGAGAC |
| leftadapR8S3 | /5Phos/GAAACAAACATCAAGAAAACAAAAT CATTATTCAAACCCA CCGAGCCCACGAGAC |
| leftadapR8S4 | /5Phos/GAAACAAACATCAAGAAAACAAAAT CACTTCATAAATCCA CCGAGCCCACGAGAC |
| leftadapR8S5 | /5Phos/GAAACAAACATCAAGAAAACAAAAT CAACTCCTAATATCA CCGAGCCCACGAGAC |
| leftadapR8S6 | /5Phos/GAAACAAACATCAAGAAAACAAAAT CAACCATACTAAACA CCGAGCCCACGAGAC |
| leftadapR8S7 | /5Phos/GAAACAAACATCAAGAAAACAAAAT CATTCCTACATTTCA CCGAGCCCACGAGAC |
| leftadapR8S8 | /5Phos/GAAACAAACATCAAGAAAACAAAAT CAAATCTTCATCCCA CCGAGCCCACGAGAC |
| leftadapR8S9 | /5Phos/GAAACAAACATCAAGAAAACAAAAT CAAACACTCTATTCA CCGAGCCCACGAGAC |
| leftadapR8S10 | /5Phos/GAAACAAACATCAAGAAAACAAAAT CAACTCAAACATACA CCGAGCCCACGAGAC |
| leftadapR8S11 | /5Phos/GAAACAAACATCAAGAAAACAAAAT CATACCCTTTTCTCA CCGAGCCCACGAGAC |
| leftadapR8S12 | /5Phos/GAAACAAACATCAAGAAAACAAAAT CATCATACCTACTCA CCGAGCCCACGAGAC |
| leftadapR8S13 | /5Phos/GAAACAAACATCAAGAAAACAAAAT CAAAACTCTCTCTCA CCGAGCCCACGAGAC |
| leftadapR8S14 | /5Phos/GAAACAAACATCAAGAAAACAAAAT CAAACCCAACTCACA CCGAGCCCACGAGAC |
| leftadapR8S15 | /5Phos/GAAACAAACATCAAGAAAACAAAAT CATTCTCCCACCTCA CCGAGCCCACGAGAC |
| leftadapR8S16 | /5Phos/GAAACAAACATCAAGAAAACAAAAT CATATCTAATCTCCA CCGAGCCCACGAGAC |
| rightadapor | TCGTCGGCAGCGTC TTGTTTGGATTATACT |

##### M13.8 sequences for the binary counting program

| | |
|---|---|
| reg8Cell1OneLeft | /5Phos/ATTTCAATTACCTGA GCAAAAGAAGATGAT |
| reg8Cell1OneRight | /5Phos/CAAAAGC GCGCAGAGGCG AATTATTC |
| reg8Cell1ZeroLeft | /5Phos/AATTATTC ATTTCAATTACCTGA GCAAAAGAAGATGAT |
| reg8Cell1ZeroRight | /5Phos/CAAAATC GCGCAGAGGCG |
| reg8Cell2OneLeft | /5Phos/CGCCTGATTGCT TTGAATACCAAGTTA |
| reg8Cell2OneRight | /5Phos/TTTTACAG CGGGAGAAACAATAA CGGATT |
| reg8Cell2ZeroLeft | /5Phos/CGGATT CGCCTGATTGCT TTGAATACCAAGTTA |
| reg8Cell2ZeroRight | /5Phos/TTTTACAT CGGGAGAAACAATAA |
| reg8Cell3OneLeft | /5Phos/ACGTCAGATGAATAT ACAGTAACAGTACC |
| reg8Cell3OneRight | /5Phos/TAAAGAAA GTGCGTAGATTTTC AGGTTTA |
| reg8Cell3ZeroLeft | /5Phos/AGGTTTA ACGTCAGATGAATAT ACAGTAACAGTACC |
| reg8Cell3ZeroRight | /5Phos/TAAAGAAA TTGCGTAGATTTTC |
| reg8Cell4OneLeft | /5Phos/CATATCAAAATTATTTG CACGTAAAACAGAAA |
| reg8Cell4OneRight | /5Phos/TCTGAAG AATGGAAGGGTTAG AACCTAC |
| reg8Cell4ZeroLeft | /5Phos/AACCTAC CATATCAAAATTATTTG CACGTAAAACAGAAA |
| reg8Cell4ZeroRight | /5Phos/TCTGAAT AATGGAAGGGTTAG |
| reg8Instru11 | TATCTCTAAC AATTATTC ATTTCAATTACCTGA GCAAAAGAAGATGAT |
| reg8Instru12 | TATCTCTAAC CGGATT CGCCTGATTGCT TTGAATACCAAGTTA CAAAATC GCGCAGAGGCG |
| reg8Instru13 | TATCTCTAAC AGGTTTA ACGTCAGATGAATAT ACAGTAACAGTACC TTTTACAT CGGGAGAAACAATAA |
| reg8Instru14 | TATCTCTAAC AACCTAC CATATCAAAATTATTTG CACGTAAAACAGAAA TAAAGAAA TTGCGTAGATTTTC |
| reg8Instru15 | TTCTATTAC CGGATTATACT TCTGAAT AATGGAAGGGTTAG |
| reg8Instru21 | ATCATCTTCTTTTGC TCAGGTAATTGAAAT GAATAATT GTTAGAGATA |
| reg8Instru22 | CGCCTCTGCGC GATTTTG TAACTTGGTATTCAA AGCAATCAGGCG AATCCG GTTAGAGATA |
| reg8Instru23 | TTATTGTTTCTCCCG ATGTAAAA GGTACTGTTACTGT ATATTCATCTGACGT TAAACCT GTTAGAGATA |
| reg8Instru24 | GAAAATCTACGCAA TTTCTTTA TTTCTGTTTTACGTG CAAATAATTTTGATATG GTAGGTT GTTAGAGATA |
| reg8Instru25 | CTAACCCTTCCATT ATTCAGA AGTATAATCCG GTAATAGAA |
| reg8Instru41 | GCGCAGAGGCG AATTATTC ATTTCAATTACCTGA |
| reg8Instru42 | CGGGAGAAACAATAA CGGATT CGCCTGATTGCT |
| reg8Instru43 | TTGCGTAGATTTTC AGGTTTA ACGTCAGATGAATAT |
| reg8Instru44 | AATGGAAGGGTTAG AACCTAC CATATCAAAATTATTTG |
| reg8Instru61 | ATCATCTTCTTTTGC TCAGGTAATTGAAAT GAATAATT |
| reg8Instru62 | TAACTTGGTATTCAA AGCAATCAGGCG AATCCG |
| reg8Instru63 | GGTACTGTTACTGT ATATTCATCTGACGT TAAACCT |
| reg8Instru64 | TTTCTGTTTTACGTG CAAATAATTTTGATATG GTAGGTT |

**M13.9**

M13 sequence window:

CCTCAATTCCTTTCAACTGTTGATTTGCCAACTGACCAGATATTGATTGAGGGTTTGATATTTGAGGTTCA
GCAAGGTGATGCTTTAGATTTTTCATTTGCTGCTGGCTCTCAGCGTGGCACTGTTGCAGGCGGTGTTAATA
CTGACCGCCTCACCTCTGTTTTATCTTCTGCTGGTGGTTCGTTCGGTATTTTTAATGGCGATGTTTTAGGG
CTATCAGTTCGCGCATTAAAGACTAATAGCCATTCAAAAATATTGTCTGTGC

| | |
|---|---|
| leftadapR9S1 | /5Phos/AATCAACAGTTGAAAGGAATTGAGG CAAAACAAAACCTCA CCGAGCCCACGAGAC |
| leftadapR9S2 | /5Phos/AATCAACAGTTGAAAGGAATTGAGG CAACATATCAATTCA CCGAGCCCACGAGAC |
| leftadapR9S3 | /5Phos/AATCAACAGTTGAAAGGAATTGAGG CATTATTCAAACCCA CCGAGCCCACGAGAC |
| leftadapR9S4 | /5Phos/AATCAACAGTTGAAAGGAATTGAGG CACTTCATAAATCCA CCGAGCCCACGAGAC |
| leftadapR9S5 | /5Phos/AATCAACAGTTGAAAGGAATTGAGG CAACTCCTAATATCA CCGAGCCCACGAGAC |
| leftadapR9S6 | /5Phos/AATCAACAGTTGAAAGGAATTGAGG CAACCATACTAAACA CCGAGCCCACGAGAC |
| leftadapR9S7 | /5Phos/AATCAACAGTTGAAAGGAATTGAGG CATTCCTACATTTCA CCGAGCCCACGAGAC |
| leftadapR9S8 | /5Phos/AATCAACAGTTGAAAGGAATTGAGG CAAATCTTCATCCCA CCGAGCCCACGAGAC |
| leftadapR9S9 | /5Phos/AATCAACAGTTGAAAGGAATTGAGG CAAACACTCTATTCA CCGAGCCCACGAGAC |
| leftadapR9S10 | /5Phos/AATCAACAGTTGAAAGGAATTGAGG CAACTCAAACATACA CCGAGCCCACGAGAC |
| leftadapR9S11 | /5Phos/AATCAACAGTTGAAAGGAATTGAGG CATACCCTTTTCTCA CCGAGCCCACGAGAC |
| leftadapR9S12 | /5Phos/AATCAACAGTTGAAAGGAATTGAGG CATCATACCTACTCA CCGAGCCCACGAGAC |
| leftadapR9S13 | /5Phos/AATCAACAGTTGAAAGGAATTGAGG CAAAACTCTCTCTCA CCGAGCCCACGAGAC |
| leftadapR9S14 | /5Phos/AATCAACAGTTGAAAGGAATTGAGG CAAACCCAACTCACA CCGAGCCCACGAGAC |
| leftadapR9S15 | /5Phos/AATCAACAGTTGAAAGGAATTGAGG CATTCTCCCACCTCA CCGAGCCCACGAGAC |
| leftadapR9S16 | /5Phos/AATCAACAGTTGAAAGGAATTGAGG CATATCTAATCTCCA CCGAGCCCACGAGAC |
| rightadapor | TCGTCGGCAGCGTC ACAATATTTTTGAATG |

**M13.9 sequences for the binary counting program**

| | |
|---|---|
| reg9Cell1OneLeft | /5Phos/CCCTCAATCAATATC TGGTCAGTTGGCA |
| reg9Cell1OneRight | /5Phos/TCACCG TGCTGAACCTCAA ATATCAAA |
| reg9Cell1ZeroLeft | /5Phos/ATATCAAA CCCTCAATCAATATC TGGTCAGTTGGCA |
| reg9Cell1ZeroRight | /5Phos/TCACCT TGCTGAACCTCAA |
| reg9Cell2OneLeft | /5Phos/GCCAGCAGCAAAT GAAAAATCTAAAGCA |
| reg9Cell2OneRight | /5Phos/CGCCT GCGACAGTGCCAC GCTGAGA |
| reg9Cell2ZeroLeft | /5Phos/GCTGAGA GCCAGCAGCAAAT GAAAAATCTAAAGCA |
| reg9Cell2ZeroRight | /5Phos/CGCCT GCAACAGTGCCAC |
| reg9Cell3OneLeft | /5Phos/CAGAGGTGAGGC GGTCAGTATTAACAC |
| reg9Cell3OneRight | /5Phos/GAACGA ACCGCCAGCAGA AGATAAAA |
| reg9Cell3ZeroLeft | /5Phos/AGATAAAA CAGAGGTGAGGC GGTCAGTATTAACAC |
| reg9Cell3ZeroRight | /5Phos/GAACGA ACCACCAGCAGA |
| reg9Cell4OneLeft | /5Phos/AGCCCTAAAACATC GCCATTAAAAATACC |
| reg9Cell4OneRight | /5Phos/GCTAGTAG TCTTTAATGCGCG AACTGAT |
| reg9Cell4ZeroLeft | /5Phos/AACTGAT AGCCCTAAAACATC GCCATTAAAAATACC |
| reg9Cell4ZeroRight | /5Phos/GCTATTAG TCTTTAATGCGCG |
| reg9Instru11 | TTACATATCC ATATCAAA CCCTCAATCAATATC TGGTCAGTTGGCA |
| reg9Instru12 | TTACATATCC GCTGAGA GCCAGCAGCAAAT GAAAAATCTAAAGCA TCACCT TGCTGAACCTCAA |
| reg9Instru13 | TTACATATCC AGATAAAA CAGAGGTGAGGC GGTCAGTATTAACAC CGCCT GCAACAGTGCCAC |
| reg9Instru14 | TTACATATCC AACTGAT AGCCCTAAAACATC GCCATTAAAAATACC GAACGA ACCACCAGCAGA |
| reg9Instru15 | TTCCTTCTTT TTTTTGAATG GCTATTAG TCTTTAATGCGCG |
| reg9Instru21 | TGCCAACTGACCA GATATTGATTGAGGG TTTGATAT GGATATGTAA |
| reg9Instru22 | TTGAGGTTCAGCA AGGTGA TGCTTTAGATTTTTC ATTTGCTGCTGGC TCTCAGC GGATATGTAA |
| reg9Instru23 | GTGGCACTGTTGC AGGCG GTGTTAATACTGACC GCCTCACCTCTG TTTTATCT GGATATGTAA |
| reg9Instru24 | TCTGCTGGTGGT TCGTTC GGTATTTTTAATGGC GATGTTTTAGGGCT ATCAGTT GGATATGTAA |
| reg9Instru25 | CGCGCATTAAAGA CTAATAGC CATTCAAAAA AAAGAAGGAA |
| reg9Instru41 | TGCTGAACCTCAA ATATCAAA CCCTCAATCAATATC |
| reg9Instru42 | GCAACAGTGCCAC GCTGAGA GCCAGCAGCAAAT |
| reg9Instru43 | ACCACCAGCAGA AGATAAAA CAGAGGTGAGGC |
| reg9Instru44 | TCTTTAATGCGCG AACTGAT AGCCCTAAAACATC |
| reg9Instru61 | TGCCAACTGACCA GATATTGATTGAGGG TTTGATAT |
| reg9Instru62 | TGCTTTAGATTTTTC ATTTGCTGCTGGC TCTCAGC |
| reg9Instru63 | GTGTTAATACTGACCGCC TCACCT CTGTTTTATCT |
| reg9Instru64 | GGTATTTTTAATGGC GATGTTTTAGGGCT ATCAGTT |

##### Sequences used for readout

##### PCR amplification

The forward and reverse primers used for PCR amplification are listed here. The sample-specific NGS barcodes were taken from the "11-2 set" from reference [63]. The list below shows 40 possible forward barcodes, and the reverse barcodes are their corresponding reverse complements.

| | |
|---|---|
| P5.B11-2.X.F | AATGATACGGCGACCACCGAGA TCTACACTCTTTCCCTACACGACGCTCTTCCGATCT <FWD BARCODE> TCGTCGGCAGCGTC |
| P7.B11-2.X.R | CAAGCAGAAGACGGCATACGAGAT GAACAA GTGACTGGAGTTCAGACGTGTGCTCTTCCGATCT <REV BARCODE> GTCTCGTGGGCTCGG |

| | | | | | | | |
|---|---|---|---|---|---|---|---|
| B.1.F | AACAACAACAC | B.2.F | AACAACGGTGG | B.3.F | AACACCTTCTT | B.4.F | AACATTGAGCC |
| B.5.F | AACCGAGAAGT | B.6.F | AACCGTCCACC | B.7.F | AAGAAGCTCCG | B.8.F | AAGACCATAGG |
| B.9.F | AAGCGGTTATA | B.10.F | AAGGTACTGAA | B.11.F | AAGTTCATTCC | B.12.F | AATAGTGTCGG |
| B.13.F | AATTCCGCGCT | B.14.F | ACAATGCGAAT | B.15.F | ACCACAGATTA | B.16.F | ACCGCTATGCC |
| B.17.F | ACCTCCTGTAA | B.18.F | ACGATGTTGTG | B.19.F | ACGGAACCAGC | B.20.F | ACGTAGGATAA |
| B.21.F | ACTATTACTCC | B.22.F | AGAAGAAGAGA | B.23.F | AGCATCTCAAT | B.24.F | AGCTCGGACCT |
| B.25.F | AGGTGAGTCTT | B.26.F | AGTCTAGCGTT | B.27.F | AGTGCGCCAAC | B.28.F | ATGGAGGACGG |
| B.29.F | ATTAACATGCC | B.30.F | ATTGGTGCATA | B.31.F | CAAGCCTGTGG | B.32.F | CAATCTTACAG |
| B.33.F | CACGCATAACG | B.34.F | CACTGGTTCCG | B.35.F | CAGTAATTGGA | B.36.F | CATACGACCGC |
| B.37.F | CCAGAGCACAC | B.38.F | CCAGGAGGTGT | B.39.F | CCATTCGCTAG | B.40.F | CCGCGTCTTAG |

The sequencing primers are:

| | |
|---|---|
| P5.F | AATGATACGGCGACCACCGAGA |
| P7.R | CAAGCAGAAGACGGCATACGAGAT |

##### Fluorescence experiments

| | |
|---|---|
| instru11-F | AGGATGA AGGAGA TGTAAAG GTATGTG A/3Rox_N/ |
| cell1ZeroLeft-F | AGGAGA TGTAAAG GTATGTG A/3Rox_N/ |
| cell1OneLeft-F | TGTAAAG GTATGTG A/3Rox_N/ |
| cell1OneRight-F | /56-ROXN/T GTAAAGG TGTGTGT AGGAGA |
